# One-carbon pathway metabolites are altered in the plasma of subjects with Down syndrome: relation to chromosomal dosage

**DOI:** 10.1101/2021.11.30.470411

**Authors:** Beatrice Vione, Giuseppe Ramacieri, Giacomo Zavaroni, Angela Piano, Giorgia La Rocca, Maria Caracausi, Lorenza Vitale, Allison Piovesan, Caterina Gori, Gian Luca Pirazzoli, Pierluigi Strippoli, Guido Cocchi, Luigi Corvaglia, Chiara Locatelli, Maria Chiara Pelleri, Francesca Antonaros

## Abstract

Down syndrome (DS) is the most common chromosomal disorder and it is caused by trisomy of chromosome 21 (Hsa21). Subjects with DS show a large heterogeneity of phenotypes and the most constant clinical features present are typical facies and intellectual disability (ID). Several studies demonstrated that trisomy 21 causes an alteration in the metabolic profile, involving among all one-carbon cycle. We performed enzyme-linked immunosorbent assays (ELISAs) to identify the concentration of 5 different intermediates of the one-carbon cycle in plasma samples obtained from a total of 164 subjects with DS compared to 54 euploid subjects. We investigated: tetrahydrofolate (THF; DS n=108, control n=41), 5-methyltetrahydrofolate (5-methyl-THF; DS n=140, control n=34), 5-formyltetrahydrofolate (5-formyl-THF; DS n=80, control n=21), S-adenosyl-homocysteine (SAH; DS n=94, control n=20) and S-adenosyl-methionine (SAM; DS n=24, control n=15). Results highlight specific alterations of THF with a median concentration ratio DS/control of 2:3, a decrease of a necessary molecule perfectly consistent with a chromosomal dosage effect. Moreover, SAM and SAH show a ratio DS/control of 1.82:1 and 3.6:1, respectively. The relevance of these results for the biology of intelligence and its impairment in trisomy 21 is discussed, leading to the final proposal of 5-methyl-THF as the best candidate for a clinical trial aimed at restoring the dysregulation of one-carbon cycle in trisomy 21, possibly improving cognitive skills of subjects with DS.

## Introduction

Down syndrome (DS) or trisomy 21 (T21) is the most frequent aneuploidy occurring in human live births (OMIM 190685) and the most common genetic cause of intellectual disability (ID) (Antonarakis, 1991). It was discovered using a phenotype-first approach, recognizing a list of phenotypic features that were linked among themselves (Down, 1866;Romano, 2022) but its association with the presence of an extra copy of a chromosome 21 (Hsa21) was discovered only by Jerôme Lejeune and coll., who was the first to establish a link between ID and a chromosomal abnormality (Lejeune et al., 1959).

Subjects with T21 can present a variety of congenital defects and comorbidities. Only a few features occur in every individual with DS, and they are facial dysmorphology, small and hypocellular brain and cognitive impairment with highly variable severity (Roper and Reeves, 2006).

Lejeune was the first who presented a general analysis of the pathogenesis of metabolic diseases determining intellectual disability, hypothesizing that DS could be also considered a metabolic disease (Lejeune, 1979). Following Lejeune, we conducted a nuclear magnetic resonance (NMR) analysis of the metabolome in plasma and urine samples, comparing trisomic and euploid subjects (siblings of subjects with DS) to study metabolic processes possibly changed in DS as a result of the genetic imbalance (Caracausi et al., 2018). The multivariate analysis of the NMR metabolomic profiles showed a clear difference between the two groups, confirmed in a more recent study in plasma samples of a greater cohort (Antonaros et al., 2020). The univariate analysis showed a significant alteration of some metabolites. Most of the altered concentrations reflect the 3:2 gene dosage model, suggesting effects caused by the presence of three copies of Hsa21. It is interesting to note that several metabolites found increased in this study are involved in central metabolic processes such as fumarate and succinate in the Krebs cycle, pyruvate and lactate in glycolysis, and formate that is an end product of one-carbon metabolism in the mitochondrion.

One-carbon metabolism (Figure 1) is the process by which one-carbon groups at different oxidation states are used in a set of interconnected biochemical pathways driven by folate and homocysteine-methionine cycles, involved in DNA synthesis through purine and thymidylate generation, amino acid homeostasis, antioxidant generation, and epigenetic regulation (Lyon et al., 2020). Remethylation of homocysteine to methionine allows several transmethylation reactions. Folate metabolism plays at least two separate roles. It is critical for one-carbon metabolism. Moreover, reactions of folate metabolism play a role in the catabolism of choline and at least three amino acids: histidine, serine and glycine (Brosnan et al., 2015). These biochemical processes, in turn, support critical cellular functions such as cell proliferation, mitochondrial respiration and epigenetic regulation (Mattson and Shea, 2003).

**Figure 1.**
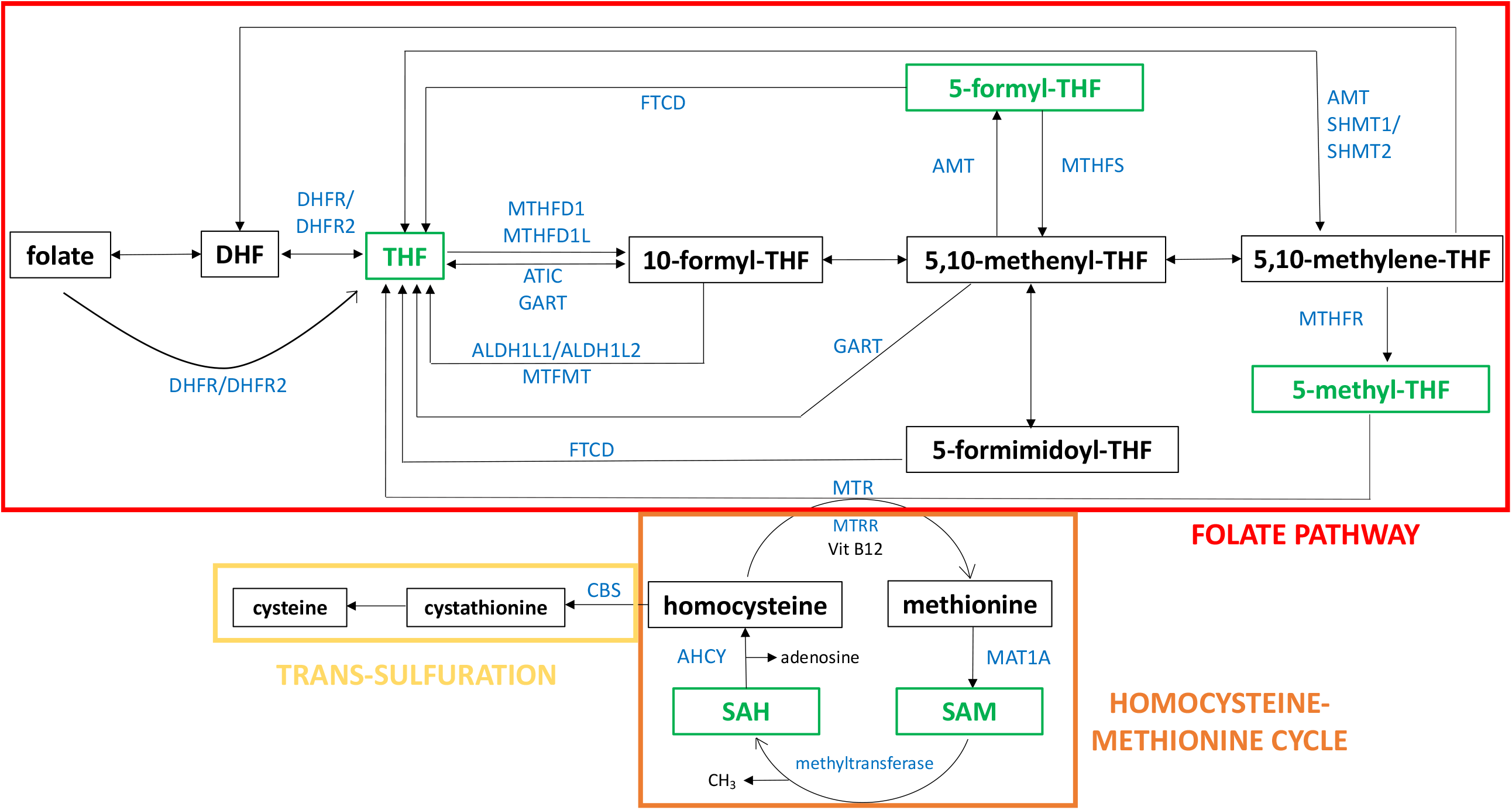
One-carbon pathway. The figure shows a schematic representation of folate pathway (in red), homocysteine-methionine cycle (in orange) and trans-sulfulration pathway (in yellow), that are part of one-carbon pathway. The metabolites analysed in our study are reported in green and the enzymes directly involved in their production or transformation are indicated with the official symbol reported in Gene NCBI (https://www.ncbi.nlm.nih.gov/gene). Alternative forms of enzyme are shown together separated by “/”. The schematic representation was realised starting from “One carbon pool by folate -Homo sapiens” of KEGG pathway database (https://www.genome.jp/kegg/pathway.html).

In his 1979 pivotal study Lejeune, comparing DS with several metabolic diseases, hypothesised that a disturbance of the one-carbon cycle could occur in DS. He thought that trisomic cells were intoxicated by an excess of gene products, which was caused by the presence of the additional Hsa21 (Lejeune, 1990). He investigated the genes located on Hsa21, and he found that several enzymes involved in the one-carbon metabolism were encoded by genes located on this chromosome, in particular: cystathionine β-synthase (*CBS*) and phosphoribosylglycinamide formyltransferase, phosphoribosylglycinamide synthetase, phosphoribosylaminoimidazole synthetase (the three subunits of GART enzyme). Moreover, the gene for the main transporter of folate, solute carrier family 19 member 1 gene (*SLC19A1*), is also located on Hsa21.

Further studies performed by Lejeune and coll. demonstrated that methotrexate (MTX) was twice as toxic in T21 lymphocytes as compared to control cells (Lejeune et al., 1986), in agreement with the observation of Peeters and coll. of an increased toxicity of MTX in the leukaemia therapy in children with DS compared to euploid subjects (Peeters et al., 1986). MTX is a cytotoxic drug used as a chemotherapy agent and immune system suppressant (Chabner et al., 1985;Morgan et al., 1987). MTX is a folate analogue which inhibits the activity of the dihydrofolate reductase (DHFR) enzyme involved in the one-carbon metabolic cycle (Lejeune et al., 1986).

In 2019 we evaluated the rescue effect on MTX toxicity mediated by folate and some of its derivatives in primary human fibroblast cell lines obtained from euploid and T21 donors (Vitale et al., 2019). This study shows that 5-methyltetrahydrofolate (5-methyl-THF) and 5-formyltetrahydrofolate (5-formyl-THF or folinic acid), but not THF, treatments have significant protective effects on both euploid and T21 cells during MTX treatment.

In a recent survey, erythrocyte folate concentrations in children with DS were compared with a cohort of subjects with juvenile arthritis as euploid control, in order to understand if the MTX treatment toxicity in children with DS is related to altered erythrocyte folate concentrations. They reported a lower concentration level of total folate, 5-methyl-THF and 5,10-methenyl-THF in the erythrocytes of subjects with DS. These reductions in erythrocyte folates were also associated with a decrease in short-chain folate polyglutamation 5-methyl-THFGlu3-6 and a corresponding increase in longer chain 5-methyl-THFGlu7-10 (Funk et al., 2020). These data suggest that in trisomic cells there is a biological adaptation to folate deficiency by an increased level of long-chain folate polyglutamation; indeed, increased expression of folylpolyglutamate synthetase (*FPGS*) gene was already reported in *vitro* and in animal models (Ifergan et al., 2004;Pelleri et al., 2018;Funk et al., 2020).

In subjects with DS, one-carbon metabolism was considered to be imbalanced by several authors (Lejeune, 1979;Rosenblatt et al., 1982;Peeters et al., 1995;Song et al., 2015;Funk et al., 2020). In particular, subjects with DS commonly present lower blood levels of vitamin B12 (or cobalamin) and folic acid with increasing age and lower erythrocyte folates and homocysteine (Hcy) compared to healthy controls (Ueland et al., 1990;Obeid et al., 2012;Song et al., 2015;Funk et al., 2020).

In the present study, we conducted enzyme-linked immunosorbent assays (ELISAs) to collect the concentration of 5 different intermediates of one-carbon metabolism which are tetrahydrofolate (THF), 5-methyl-THF, 5-formyl-THF, S-adenosyl-homocysteine (SAH) and S-adenosyl-methionine (SAM) in plasma samples obtained from a total of 164 subjects with DS and 54 euploid subjects as control. Moreover, we investigated the association between metabolite concentrations in the DS group and control group and the correlation with folic acid, vitamin B12 and Hcy in the DS group.

## Materials and Methods Ethics statement

The Independent Ethics Committee of the Hospital - University of Bologna Policlinico S. Orsola-Malpighi Italy has granted the ethical approval for this study (number: 39/2013/U/Tess). We obtained an informed written consent from all participants to collect urine and blood samples and clinical data. Concerning minors, the consent was collected from his/her parents. All procedures were carried out in accordance with the Ethical Principles for Medical Research Involving Human Subjects of the Helsinki Declaration.

### Case selection

A total of 218 subjects were selected for this study, including 164 subjects with DS and 54 subjects as control, selected among siblings of subjects with DS and without evidence of abnormal karyotype. The study has been proposed to all subjects consecutively admitted to the Day Hospital of the Neonatology Unit, IRCCS Sant’Orsola-Malpighi Polyclinic, Bologna, Italy, in the context of routine follow up provided for DS.

For this work, we considered in the DS group subjects with diagnosis of DS with homogeneous or mosaic T21, availability of an adequate amount of plasma to perform at least one ELISA assay and a similar mean age as close as possible to control group. Concerning control group, we considered subjects with an adequate amount of plasma to perform at least one ELISA assay and a similar mean age as close as possible to DS group. The blood samples selected were treated within two hours of collection and plasma samples contaminated by residual erythrocytes after the treatment were excluded.

DS group consists of 95 males (M) and 69 females (F) with a mean age of 11.55 years old (standard deviation, SD= 6.69) and an age range from 3.0 to 37.9; control group consists of 30 M and 24 F with a mean age of 14.53 years old (standard deviation, SD=6.64) and an age range from 3.3 to 31.5. Concerning the control group, 29 subjects were siblings of 23 subjects in the DS group.

For every collected sample, parents filled out a form with information about the current fasting state, last meal, consumed medications (see Supplementary Dataset 1).

### Blood sample preparation and folic acid, vitamin B12, homocysteine detection

Two blood sample aliquots were collected in EDTA-coated blood collection tubes in the Neonatology unit.

The first aliquot was sent to “Laboratorio unico Metropolitano” (LUM) of Maggiore Hospital (Bologna, Italy) for routine blood analyses of the DS group including folic acid and vitamin B12. The concentration levels in serum were obtained by chemiluminescent immunoassay by Beckman Coulter Immunoassay Systems (Beckman Coulter, respectively REF A98032 and REF 33000).

The second aliquot was kept at room temperature and treated within two hours from blood draw. Sample was transferred to a new tube and centrifuged at 1,250 g for 10 min to separate corpuscular fraction from plasma. The plasma was isolated and centrifuged for a second time at 800 g for 30 min to eliminate residual debris. The supernatant was collected and transferred into new tubes avoiding contact with pellets or the bottom of the tube, divided in aliquots and rapidly stored in a -80°C freezer.

Every delayed treatment of the sample following its transfer to the university laboratory (>2 hours) was recorded, as one of the exclusion criteria from the analysis was sample treatment after two hours from the draw. The other one was evident contamination of plasma by hemoglobin due to hemolysis after the treatment. All procedures were conducted carefully to avoid contamination during the different steps and any anomalies like different plasma colour or precipitates after centrifugation were noted and considered in further analysis.

One or more plasma aliquots were used for ELISA assays. One plasma aliquot was sent to LUM for the detection of Hcy plasma level by an automated chemistry analyzer AU 400 Beckman Coulter (Beckman Coulter, REF FHRWAU/FHRWR100/200/1000).

Concerning concentration values of folic acid, vitamin B12 and Hcy of DS group, 25% of data was drawn from a previous work from our group (Antonaros et al., 2021).

### Enzyme-linked immunosorbent assay (ELISA)

We evaluated the quantitative measurement of THF, 5-methyl-THF, 5-formyl-THF and SAH plasma concentrations using specific ELISA kits manufactured by MyBioSource (San Diego, California, USA) and SAM plasma concentration using a specific ELISA kit manufactured by Biovision (Milpitas, California, USA). We performed 6 ELISA assays to obtain concentration values of 5-methyl-THF, 5-formyl-THF and SAH, 5 ELISA assays to obtain concentration values of THF and 1 ELISA assay to collect concentration values of SAM.

“General Tetrahydrofolic Acid (THFA) ELISA assay”, “Human 5-methyltetrahydrofolate ELISA assay” and “Enzyme-linked Immunosorbent assay for 5-formyl-THF (folinic acid)” MyBioSource kits are based on a competitive inhibition enzyme technique.

Regarding THF, specifications were: detection range, 1.23-100 ng/mL; sensitivity, 0.57 ng/mL; intra-assay precision, Coefficient of Variation (CV) <10%; inter-assay precision, CV <12%; no significant cross-reactivity or interference between THFA and analogues was observed.

Regarding 5-methyl-THF, assay specifications were: detection range, 0.156-10 ng/mL; sensitivity, 0.094 ng/mL; intra-assay precision, CV <8%; inter-assay precision, CV <10%; no significant cross-reactivity or interference between 5-methyl-THF and analogues was observed.

Regarding 5-formyl-THF, assay specifications were: detection range, 117.2-30000 pg/mL; sensitivity, <41.5 pg/mL; intra-assay precision, CV <10%; inter-assay precision, CV <12%; no significant cross-reactivity or interference between Folinic Acid (FA) and analogues was observed.

MyBioSource “Human S-Adenosyl-Homocysteine (SAH) ELISA assay” and Biovision “S-Adenosylmethionine (SAM) ELISA” kits employ a double antibody sandwich technique.

Regarding SAH, assay specifications were: detection range, 20 ng/mL-0.312 ng/mL; sensitivity, up to 0.06 ng/mL; intra-assay precision, CV **≤**8%; inter-assay precision, CV **≤**12%; no cross-reaction with other factors.

Regarding SAM, assay specifications were: detection range, 0.39-25 mcg/mL; sensitivity, <0.234 mcg/mL; intra-assay precision, CV <8%; inter-assay precision, CV<10%; no significant cross-reactivity or interference between SAM and analogues was observed.

Ninety-six well plates were used for all the assays and all standard and plasma samples were tested in duplicate. Due to preliminary studies, plasma samples used for the measurement of 5-methyl-THF were diluted 1:10 using Sample Dilution Buffer supplied by the kit and plasma samples used for the measurement of SAH were diluted 1:5 in Plates 1, 2, 3 and 4, and 1:3 in Plates 4 and 5 to avoid excessive dilution of the metabolite, using Sample Diluent supplied by the kit. The standard samples were provided by the kits and were reconstituted and serially diluted as suggested.

In order to perform the assays, the manufacturer instructions of each kit were followed, and the final spectrophotometric reading was carried out by microplate reader (Perkin Elmer Wallac 1420 Victor 2 Multi-Label) set at a wavelength of 450 nm.

In order to create the standard curve for each ELISA assay, standard O.D. mean values were plotted on the x-axis and the known standard concentration values were plotted on the y-axis using Microsoft Excel and following manufacturer instructions. The transformation of standard concentration values in their logarithm (Log_10_) is required to build the standard curves in THF and 5-formyl-THF ELISA assays. The subtraction of the background O.D. mean value (“standard 8”) from the O.D. mean values of the other standards and plasma samples is required in SAH ELISA assay. The transformation of standard O.D. mean values in their inverse (1/O.D. mean) is required to build the standard curves in the case of the SAM assay.

The polynomial trend lines of each plot were created using Excel, and the resulting polynomial equations (y = a + bx + cx^2^) were used to determine metabolite concentrations of plasma samples using interpolation.

### Statistical analyses

Statistical analyses were carried out with SPSS Statistics (IBM, Version 25 for Mac OS X) and were performed using the data available in Supplementary Dataset 1. For all results, a p<0.05 was considered statistically significant. An r<0.4 was considered as weakly correlated, 0.4<r<0.7 as moderately correlated and r>0.7 as strongly correlated.

A first descriptive analysis was made using SPSS Statistics software in order to obtain a general view of the data included in Supplementary Dataset 1. It was performed with SPSS Statistics software as follows: from the leading software Menu, we selected “Analyze” and then “Descriptive analysis”, we then chose “Frequencies”, we included our concentration data in the “variables” box and finally we chose “Mean”, “Median”, “Modal value”, “Standard deviation”, “Variance”, “Range”, “Minimum”, “Maximum” in the “Statistics” box. The presence of strong outliers in concentration level distribution was performed with

SPSS Statistics software as follows: from the leading software Menu we selected “Analyze” and then “Descriptive statistics”, we then chose “Explore”, and included concentration levels in “Dependent List”, finally in “Statistics section” we selected the “Outliers” and “Percentiles” options. SPSS Statistics software indicates strong outliers with an asterisk in the graph.

For each metabolite (THF, 5-methyl-THF, 5-formyl-THF, SAH and SAM) we performed an unpaired student t-test between DS and control group using the “Graph Pad” t-test calculator online (https://www.graphpad.com/quickcalcs/ttest1.cfm). In order to perform the analysis, we chose “Enter or paste up to 2000 rows”, then we chose “Unpaired t-test”, entered for each DS metabolite data in “Group one” column and control data in “Group two” column and in the end, we chose “Calculate now”.

A linear correlation was used to determine if the correlation between age and molecule levels existed. It was performed with SPSS Statistics software as follows: from the leading software Menu, we selected “Analyze” and then “Correlate”, we then chose “Bivariate correlation”, and finally we included our data in the “variables” box.

Unpaired t-test was used to test whether sex and fasting/non-fasting state might affect the main results. Unpaired t-test was performed with SPSS Statistics software as follows: from the leading software Menu we selected “Analyze” and then “Compare means”, we then chose “independent-samples T-test” and finally we included our data in the “test variables” box and inserted “Sex” or “Fasting/non-fasting” in the “grouping variable” box.

SPSS Statistics software was used to perform a linear correlation between the level of each molecule and the levels of all the other molecules. Briefly, from the main Menu of the “SPSS Statistics” software, we selected “Analyze” and then “Correlation”; we chose “Bivariate” and finally we inserted our data in the main box. Partial correlation analyses checked for the effect of chronological age were used to investigate associations between the level of the involved molecules and other molecules. These analyses were performed as follows: from the main Menu of the “SPSS Statistics” software, we selected “Analyze” and then “Correlation”; we chose “Partial correlation” and finally we inserted our data in the main box and inserted “Age” in the “Check by” box. The Heat Map figures representing metabolite correlations (THF, 5-methyl-THF, 5-formyl-THF, SAH and SAM) were generated using JMP Pro software (Version 14 of the SAS System for Mac OS X, SAS Institute Inc., Cary, NC, USA).

## Results

### Enzyme-linked Immunosorbent Assay (ELISA)

We selected 164 subjects with DS and 54 euploid subjects. Concerning subjects with DS, only one has T21 mosaicism at 94%. In the DS group we obtained the concentration values of THF in 108 subjects, of 5-methyl-THF in 140 subjects, of 5-formyl-THF in 79 subjects, of SAH in 94 subjects and of SAM in 24 subjects. In the control group we obtained the concentration values of THF in 41 subjects, of 5-methyl-THF in 34 subjects, of 5-formyl-THF in 21 subjects, of SAH in 20 subjects and of SAM in 15 subjects (see Supplementary Dataset 1). It was not possible to obtain the metabolite dosage in all the selected subjects for several reasons: the amount of plasma was not sufficient, the absorbance (O.D., optical density) mean values of plasma samples were out of the range, or there was a technical problem during the assay. Moreover, it was not possible to have a larger group of SAM concentration results due to technical problems with some of the ELISA kits purchased.

Concerning the O.D. mean values, those that were higher or lower than the O.D. mean values of standard sample ranges were not taken into consideration (they are indicated in red in Supplementary Tables 1, 2, 3 and 4). In total we did not consider the concentration values of 9 plasma samples in THF ELISA assays, 63 plasma samples in 5-formyl-THF assays, 32 plasma samples in SAH ELISA assays and 1 plasma sample in SAM ELISA assay.

In order to assess the intra-wells variability of the assays in duplicate, the Coefficient of Variation of the O.D. values expressed as percentage (%CV) was reported for each sample in Supplementary Tables 1, 2, 3, 4 and 5. The %CV was calculated dividing the standard deviation of the two measurements for the mean of the two measurements and multiplying for 100. The mean of the %CV values also named Intra-Assay CV was 9.55% for THF assays, 6.77% for 5-methyl-THF assays, 6.92% for 5-formyl-THF assays, 10.95% for SAH assays and 8.40% for SAM assay. While the mean of the %CV values was lower than 11% for all five metabolites, individual values could exceed 15% in a part of the measured sample. In particular, out of a total of 189 samples assayed there were 40 samples that ranged between 15.49% and 53.23% of %CV in THF assays; out of a total of 222 samples assayed, there were 17 samples that ranged between 15.03% and 31.15% in 5-methyl-THF assays; out of a total of 143 samples assayed, there were 15 samples that ranged between 15.44% and 36.29% in 5-formyl-THF assays; out of a total of 162 samples assayed, there were 46 samples that ranged between 15.10% and 46.36% in SAH assays; out of a total of 47 samples assayed, there were 8 samples that ranged between 15.12% and 72.55% in SAM assay.

The coefficient of determination (R^2^) of the standard curves ranged between 0.9697 and 0.9881 for THF (see Supplementary Table 1), between 0.922 and 0.9707 for 5-methyl-THF (see Supplementary Table 2), between 0.9048 and 0.9433 for 5-formyl-THF (see Supplementary Table 3), between 0.9998 and 0.9956 for SAH (see Supplementary Table 4) and 0.9996 for SAM (see Supplementary Table 5).

### Statistical analyses

Overall, 3 strong outliers were identified among 5-formyl-THF and vitamin B12 concentration values of the DS group and SAH concentration values of control group that are reported in red in Supplementary Dataset 1.

The main results of descriptive analyses of THF, 5-methyl-THF, 5-formyl-THF, SAH and SAM without strong outliers are shown in Table 1. The detailed results of descriptive analysis with and without strong outliers are shown in Supplementary Table 6.

**Table 1.**
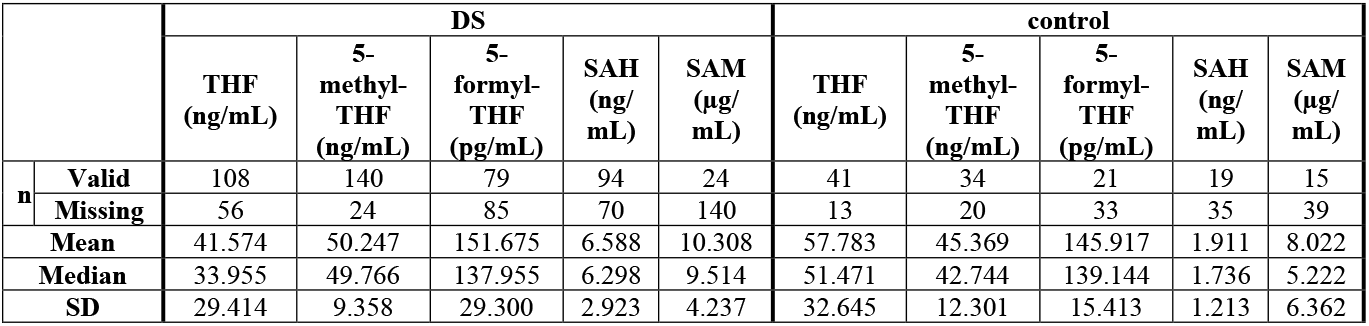
Main results of descriptive analysis of DS and control group excluding strong outliers. The values of subjects with Down syndrome (DS) are reported on the left of the table and the values of normal control subjects (control) are reported on the right of the table. For each metabolite the number (n) of plasma samples valid or missing are reported. Also the mean, median and standard deviation (SD) values are shown. The detailed results of descriptive analysis with and without strong outliers are shown in Supplementary Table 6.

|  |  | DS |  |  |  |  | control |  |  |  |  |
| --- | --- | --- | --- | --- | --- | --- | --- | --- | --- | --- | --- |
| | | THF<br>(ng/mL) | 5-methyl-THF<br>(ng/mL) | 5-formyl-THF<br>(pg/mL) | SAH<br>(ng/<br>mL) | SAM<br>($\mu\text{g}/$<br>mL) | THF<br>(ng/mL) | 5-methyl-THF<br>(ng/mL) | 5-formyl-THF<br>(pg/mL) | SAH<br>(ng/<br>mL) | SAM<br>($\mu\text{g}/$<br>mL) |
| n | Valid | 108 | 140 | 79 | 94 | 24 | 41 | 34 | 21 | 19 | 15 |
|  | Missing | 56 | 24 | 85 | 70 | 140 | 13 | 20 | 33 | 35 | 39 |
|  | Mean | 41.574 | 50.247 | 151.675 | 6.588 | 10.308 | 57.783 | 45.369 | 145.917 | 1.911 | 8.022 |
|  | Median | 33.955 | 49.766 | 137.955 | 6.298 | 9.514 | 51.471 | 42.744 | 139.144 | 1.736 | 5.222 |
|  | SD | 29.414 | 9.358 | 29.300 | 2.923 | 4.237 | 32.645 | 12.301 | 15.413 | 1.213 | 6.362 |

The results of unpaired student t-test between DS and control concentration values of each metabolite showed that the difference between DS and control concentrations is statistically significant for THF (p-value=0.0041), 5-methyl-THF (p-value=0.015) and SAH (p-value=0.0001) when the strong outliers are excluded (see Table 2) or included (see Supplementary Table 7) in the analyses.

Figure 2 shows the variation of THF, 5-methyl-THF, 5-formyl-THF, SAH and SAM concentrations in subjects with DS and normal control subjects excluding strong outliers. In Supplementary Figure 1 the variation of 5-formyl-THF and SAH concentration in subjects with DS and normal control subjects is reported including strong outliers.

**Table 2.**
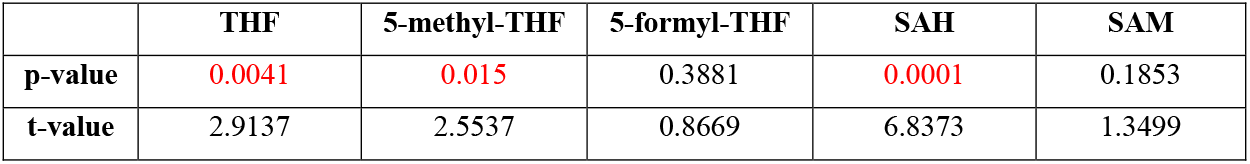
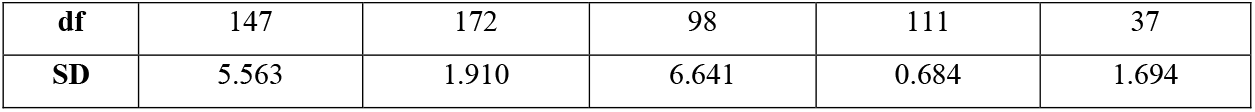
Unpaired student t-test between DS and control group concentration values of each metabolite excluding strong outliers. Significant p-values (<0.05) are reported in red in the first line. T-value, degrees of freedom (df) and the standard deviation (SD) are reported in the other lines. The detailed results of unpaired student t-test with and without strong outliers are shown in Supplementary Table 7.

**Figure 2.**
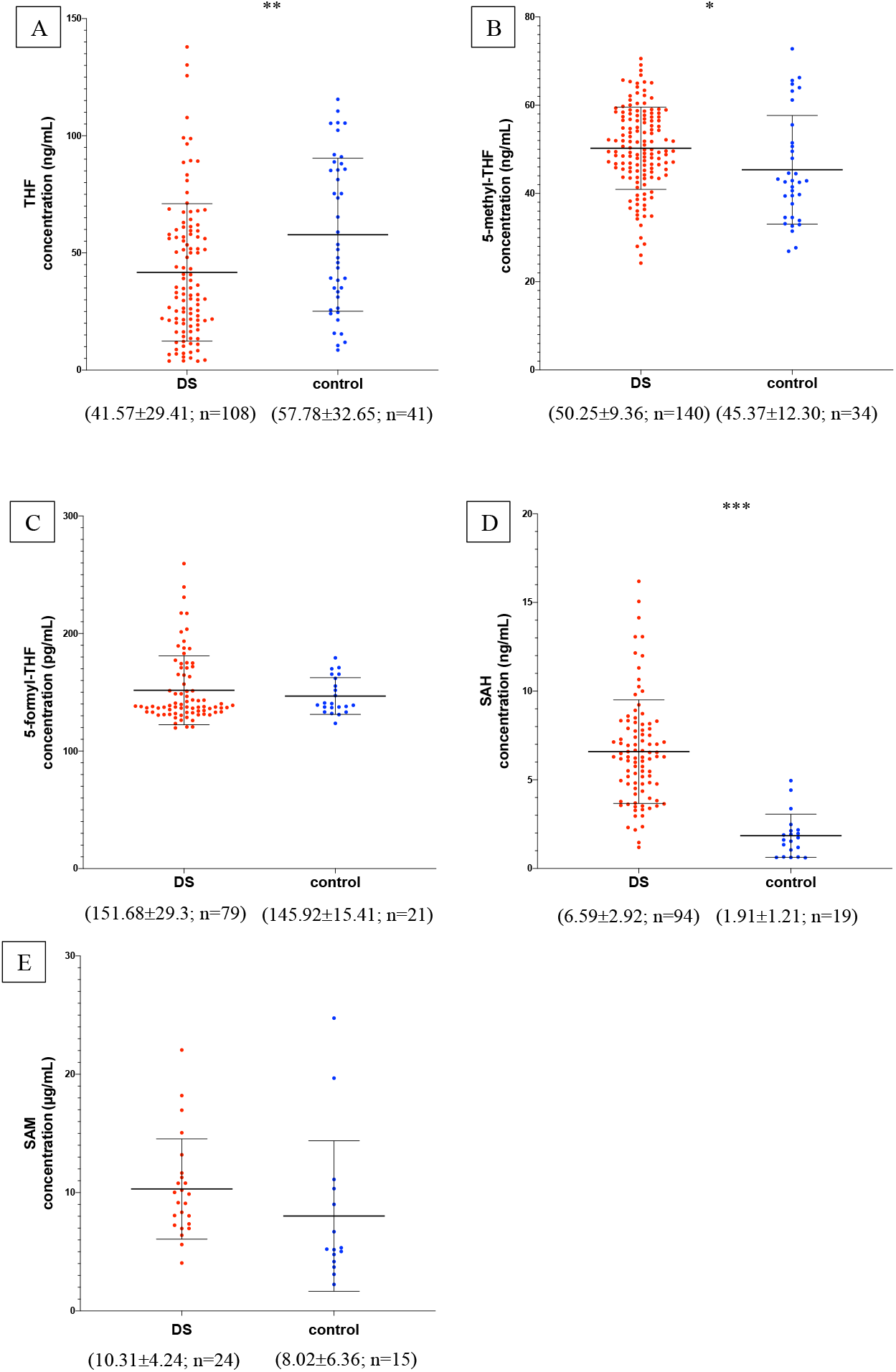
Metabolite concentrations in subjects with DS and normal control subjects. The graphs report metabolite plasma levels of each subject in the study. On the x-axis there is the subdivision of subjects in DS and control groups. Subjects with DS are represented like red dots and normal control subjects are represented like blue dots. On the y-axis the concentration of the metabolite in µg/mL, ng/mL or pg/mL is reported. The asterisks above the graph indicate the level of statistical significance (*=p<0.05; **=p<0.005; ***=p<0.0005). The middle black lines indicate the mean concentration values for each group and the external black lines indicate standard deviation (SD) values. The mean concentration, SD values and the number of subjects (n) are reported below each graph for DS and control groups. Figure 2a shows THF concentrations; Figure 2b shows 5-methyl-THF concentrations; Figure 2c shows 5-formyl-THF concentrations excluding strong outliers; Figure 2d shows SAH concentrations excluding strong outliers; Figure 2e shows SAM concentrations. The graphs were created with GraphPad Prism software v.6.0 (San Diego, CA).

In Table 3 the ratio of mean concentration values and the ratio of median concentration values between DS and control groups is reported.

**Table 3.**
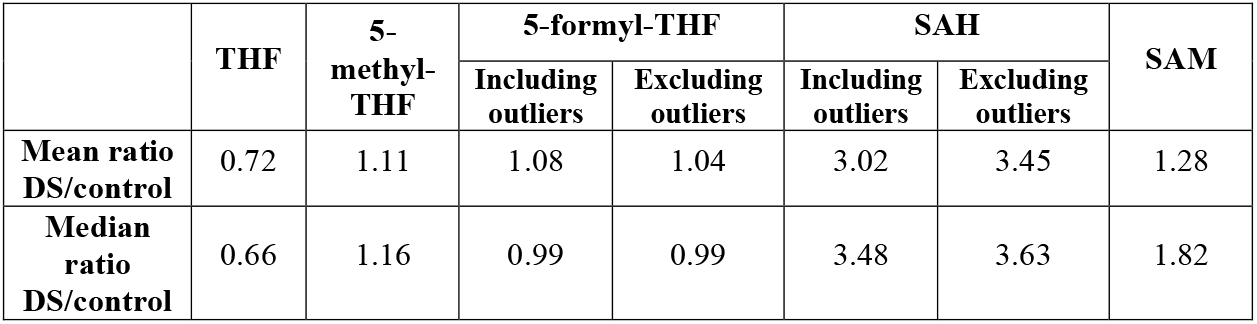
Ratio of mean concentration values and median values between DS and control groups. For each metabolite in the study the ratios of mean concentration levels and median levels between DS and control groups (Mean ratio DS/control and Median ratio DS/control) is reported, following data in Table 1. For 5-formyl-THF and SAH the ratio including and excluding strong outliers are both showed.

|  | THF | 5-methyl-THF | 5-formyl-THF |  | SAH |  | SAM |
| --- | --- | --- | --- | --- | --- | --- | --- |
|  |  |  | Including outliers | Excluding outliers | Including outliers | Excluding outliers |  |
| **Mean ratio DS/control** | 0.72 | 1.11 | 1.08 | 1.04 | 3.02 | 3.45 | 1.28 |
| **Median ratio DS/control** | 0.66 | 1.16 | 0.99 | 0.99 | 3.48 | 3.63 | 1.82 |

A statistically significant weak correlation (r=Pearson correlation coefficient) was found between age and THF concentration levels in control group (r= 0.395 and p-value= 0.011). The significance was lost in DS group. Whether or not the strong outliers were considered did not change the results (see respectively Table 4 and Supplementary Table 8B). A statistically significant moderate correlation was found between age and Hcy concentration levels in the DS group (r=0.593 and p-value < 0.001) (see Supplementary Table 8A and 8B).

**Table 4.** Bivariate correlation between age and each concentration levels in DS and control groups excluding strong outliers. The table reports for each metabolite the results of the statistical analyses in DS and control groups indicated by Pearson correlation coefficient (r) and two-tailed significance (p-value). Significant r (>0.4) and p-values (<0.05) are reported in red. The detailed results of bivariate correlation with and without strong outliers are shown in Supplementary Table 8.

|  | DS |  |  |  |  | control |  |  |  |  |
| --- | --- | --- | --- | --- | --- | --- | --- | --- | --- | --- |
|  | THF | 5-methyl-THF | 5-formyl-THF | SAH | SAM | THF | 5-methyl-THF | 5-formyl-THF | SAH | SAM |
| **r** | 0.089 | -0.085 | 0.128 | 0.152 | 0.099 | 0.395 | 0.239 | 0.193 | 0.102 | 0.115 |
| **P-value** | 0.362 | 0.318 | 0.261 | 0.145 | 0.644 | 0.011 | 0.296 | 0.379 | 0.678 | 0.682 |

The unpaired t-test did not identify differences between males and females concerning the concentration of all the molecules investigated (see Supplementary Table 9).

The unpaired t-tests comparing values in fasting or non-fasting state highlighted a statistically significant difference in vitamin B12 levels in the DS group when the strong outliers are both included (p-value=0.004) (see Supplementary Table 10A) or excluded (p-value=0.005) (see Supplementary Table 10B) in the analyses.

The linear correlation analysis identified a statistically significant moderate negative correlation (r=-0.628 and p-value=0.029) between SAM and vitamin B12 levels in the non-fasting DS group. A statistically significant strong positive correlation (r=0.81 and p-value=0.003) between SAM and 5-formyl-THF levels was found in DS group. The correlation was not maintained in the control group (see Supplementary Table 11, Figure 3 and Supplementary Figure 2).

**Figure 3.**
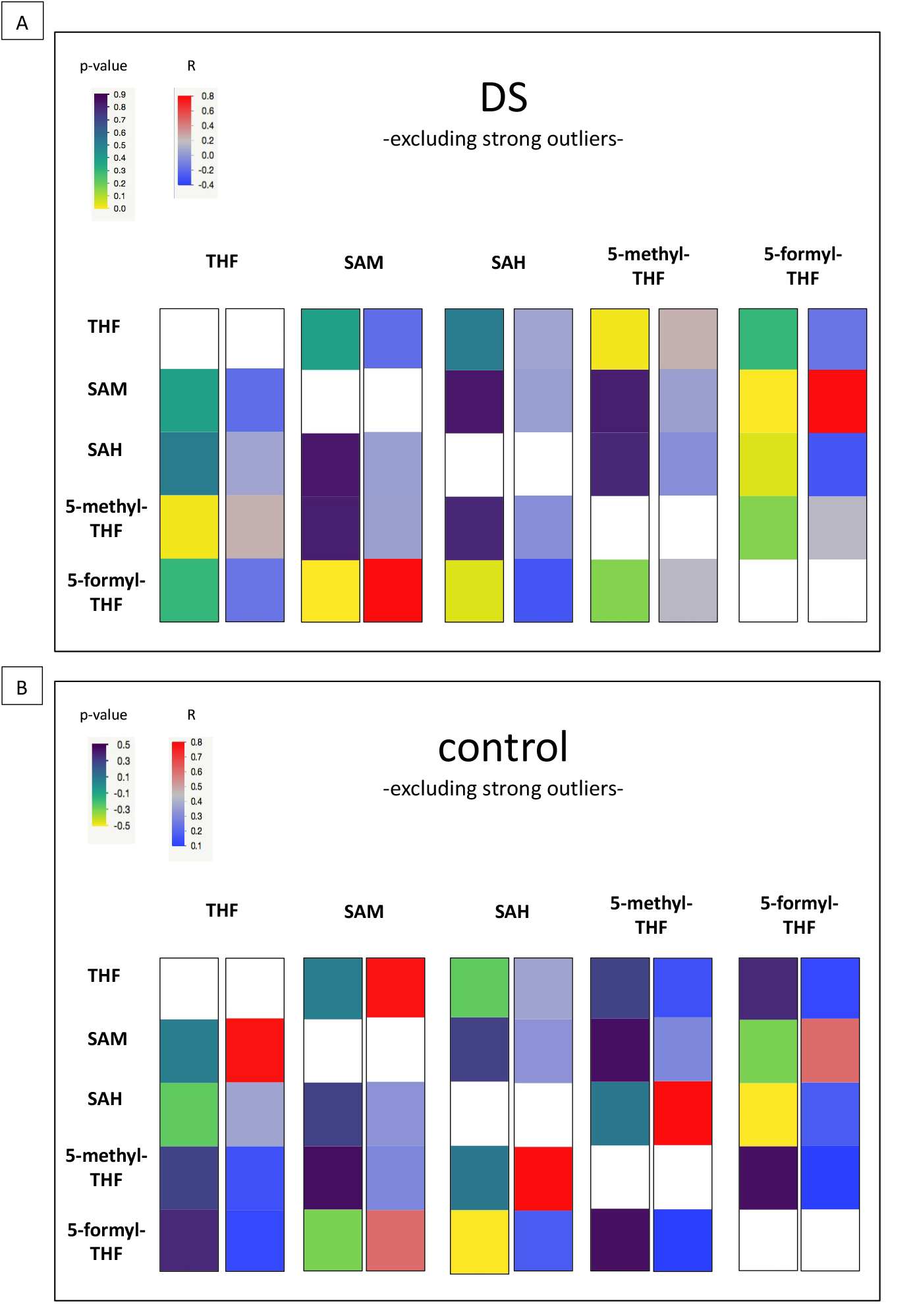
Heat Map figure of bivariate correlation between levels of each metabolite and levels of all the other metabolites excluding strong outliers. Figure 3a presents bivariate correlations in DS group and Figure 3b presents bivariate correlation in the control group (for complete data see Supplementary Table 10). At the top left of the figures the color codes for Pearson correlation coefficient (r) and two tailed-significance (p-value) are reported. For each metabolite indicated above, two bars are shown: the first represents p-values and the second r values. Every bar is divided in five segments corresponding to the five correlations with other metabolites.

## Discussion

Our results highlight plasma level alteration of some intermediates of one-carbon metabolism in a group of subjects with DS compared to a group of euploid subjects as control. We were able to enrol a lower number of control subjects compared to subjects with DS, a common situation (see for example (Caracausi et al., 2018;Orozco et al., 2019;Antonaros et al., 2020;Funk et al., 2020) often due to the difficulty in the enrollment of healthy children in a medical setting. However, all the statistical analyses performed accurately account for any difference in the group size and report the significance of the results after compensation for this factor (Cardinal, 2015). A point of strength of our analysis is the number of healthy controls that are siblings of children with DS included in the compared group because the similarity in genetic background between siblings helps to highlight the differences due to the additional copy of Hsa21 in the trisomic subjects (an extreme of this condition has been in literature with the use of cells derived from two monozygotic twins discordant for trisomy 21, (Hibaoui et al., 2014).

We reported a statistically significant difference of THF (p-value=0.0041), 5-methyl-THF (p-value=0.015) and SAH (p-value=0.0001) plasma levels between DS and control groups (see Table 2). Concerning 5-formyl-THF and SAM, the difference is not statistically significant (respectively p-value=0.3881 and 0.1853) (see Table 2).

THF is the metabolic active form of folates in cells and it serves as one-carbon carrier in most folate-mediated reactions (Zheng and Cantley, 2019). THF is the folate form most interconnected in the folate pathway and it is the product of ten enzymatic reactions as shown in Figure 1. Moreover, Lejeune analyzed the genetic diseases causing biochemical alterations that affect the nervous system and he found out that many biochemical dysregulations had an effect on THF metabolism and its chemical connections (Lejeune, 1979). Our results suggest that THF plasma concentration is lower than normal in subjects with DS and the median concentration ratio between DS and control groups is 0.66 (see Table 3), that is a 2:3 ratio, strongly suggesting a correlation with the imbalanced original event or the presence of a third Hsa21. The starting event leading to the 3:2 or 2:3 metabolite imbalances observed in DS could be the activity of critical gene determinants included in the Hsa21 region associated with diagnosis of DS (highly restricted Down syndrome critical region, HR-DSCR) (Pelleri et al., 2016;Pelleri et al., 2019) and still unknown.

Moreover, THF concentration level shows a statistically significant weak correlation with age in the control group (r=0.395 and p-value=0.011) (see Table 4). Pfeiffer and coll. reported that older age is associated with less bioactive folate (THF) and more biologically inactive folate (Pfeiffer et al., 2015). Interestigly, our results report that the correlation between THF and age is lost in the DS group (r=0.089 and p-value=0.362) suggesting that the plasma level alteration of THF in subjects may be a stable consequence of trisomy 21 masking variation with age.

5-methyl-THF is the most abundant folate form in blood circulation and the only form able to cross the blood-brain barrier (Ducker and Rabinowitz, 2017). Our results suggest that 5-methyl-THF plasma concentration is slightly higher than normal in subjects with DS and the median concentration ratio between DS and control groups is 1.16 (see Table 3), that is a 1:1 ratio.

5-formyl-THF is a reserve of one-carbon units in cells, and it does not play a direct biosynthetic role (Ducker and Rabinowitz, 2017). Our results suggest that 5-formyl-THF plasma concentration is equal in subjects with DS compared to normal control subjects and the median concentration ratio between DS and control groups is 0.99 (see Table 3), that is a 1:1 ratio.

SAH is part of homocysteine-methionine cycle in which the universal methyl donor named S-adenosyl-methionine (SAM) is transformed in SAH by methyltransferases (Mentch and Locasale, 2016) (see Figure 1). Our results suggest that SAH plasma concentration is much higher than normal in subjects with DS and the median concentration ratio between DS and control groups is 3.63 (see Table 3).

Even if there is not a statistically significant difference of SAM plasma levels between the DS and control groups due to the distribution of the values (see Table 2), SAM concentration is higher than normal in subjects with DS and the median concentration ratio between DS and control groups is 1.82 (see Table 3).

In a previous work (Obeid et al., 2012), both SAH and SAM were found to be significantly higher in young individuals with DS compared to control subjects of comparable age, and the median SAM/SAH ratio was lower in individuals with DS. We find here that, following data in Table 1, the SAM/SAH mass median ratio is 1511 in subjects with DS and 3008 in control subjects, suggesting that SAM/SAH median ratio in subjects with DS is half (exactly 0.50) compared to control subjects. SAM/SAH ratio is a well known indicator of cellular methylation capacity and when it is decreased can correlate with reduced methylation potential (Clarke, 2001;Petrossian and Clarke, 2011). An alteration of the methylation capacity is reported in DS and was connected to aging acceleration supporting the notion that DS is a progeroid trait (Franceschi et al., 2019).

Our statistical analyses report a negative correlation between SAM concentration levels and vitamin B12 (when the subjects are in a non-fasting state) (r=-0.628 and p-value=0.029) and a positive correlation 5-formyl-THF (r=0.81 and p-value=0.003) in plasma samples in the DS group (see Figure 3 and Supplementary Table 11). These findings are difficult to interpret but it is interesting to note that the association between SAM and 5-formyl-THF is present only in the DS group and is lost in the control group, suggesting again a selective alteration of one-carbon cycle in subjects with DS. Further studies are necessary to increase the number of SAM concentration values obtained from DS and control groups.

Abnormal folate metabolism has been causally linked with many diseases. Cerebral folate deficiency (CFD), caused by folate receptor autoantibodies or germline mutations such as in *FOLR1* (folate receptor alpha), *SLC46A1* (solute carrier family 46 member 1) or *MTHFR* genes, is manifested in neurological impairments and shows deficiency of 5-methyl-THF in the cerebrospinal fluid (CSF) in the presence of low or undefined peripheral folate levels (Watkins and Rosenblatt, 2012;Pope et al., 2019). Thus, folate metabolism plays a crucial role in the brain, but is still poorly defined.

Recently, we have explored the role of five metabolite levels involved in one-carbon pathway (Hcy, folate, vitamin B12, uric acid, creatinine) in the intellectual impairment of children with DS. The findings highlighted that specific vitamin B12 and Hcy thresholds corresponded respectively with better and worse cognitive scores in children with DS suggesting a role in their cognitive development (Antonaros et al., 2021).

Today the research about DS is focusing on the improvement of cognitive status (Antonarakis, 2017). Several groups have tried to improve cognitive skills of subjects with DS using different compounds in clinical trials (Bianchi et al., 2010;Moon et al., 2010;Braudeau et al., 2011;Das et al., 2013;de la Torre et al., 2016). Even if some of them showed interesting results, as of today none of these compounds are used as effective treatments of cognitive impairment in DS.

It was demonstrated that folate deficiencies contribute to several neuropsychiatric disorders and that can be corrected by folate supplementation (Kronenberg et al., 2009;Nazki et al., 2014;Ramaekers et al., 2016). Three clinical trials tried to use 5-formyl-THF supplementation (alone and together with other compounds) to improve psychomotor and language development in young subjects with DS, but they did not evidence any significant clinical differences compared to placebo group (Ellis et al., 2008;Blehaut et al., 2010;Mircher et al., 2020). The search results for “Down syndrome” [MeSH Terms] AND “folic acid” [MeSH Terms] in Pubmed.gov (https://pubmed.ncbi.nlm.nih.gov/) did not retrieve any other clinical trial, about folates and DS.

Our results confirm that there is a dysregulation of the one-carbon pathway in subjects with DS that could be related to cognitive impairment. In particular, it is remarkable that plasma THF median concentration in subjects with DS appears to be lacking and exactly inversely proportional to the Hsa21 chromosomal dosage. This might imply a direct role of Hsa21 in impairing the progression of the folate/one-carbon cycle and opens the possibility that restoring a normal THF concentration could be important in subjects with DS. To this aim, we could evaluate administration of THF itself, or of the well known folinic acid, or of 5-methyl-THF.

Regarding THF, we have demonstrated that THF, as well as folic acid, is not able to rescue MTX toxicity in both euploid and T21 fibroblast cells, while 5-methyl-THF and 5-formyl-THF treatments have shown much better protective effects during MTX treatment (Vitale et al., 2019). Moreover, the THF node of the one-carbon metabolic network is located far from the steps related to the alteration of the methylation capacity as highlighted by the SAM/SAH altered ratio already documented in DS and confirmed here as discussed above. Finally, it should be noted that to date THF supplement is not available for human administration.

Concerning folinic acid (5-formyl-THF), it has been in use for the longest time for the treatment of CFDs (Irons et al., 1987) and is currently the most used folate form for the treatment of CFDs (Pope et al., 2019). However, its inefficiency in any improvement of the cognitive skills in children with DS has already been clearly demonstrated (Blehaut et al., 2010;Mircher et al., 2020). This finding is again consistent with the position of this metabolite in the folate cycle in that it goes toward more oxidized forms of THF, while Hcy/methionine cycle requires more reduced ones.

The 5-methyl-THF is the most reduced folate form (Nelson, 2017), the biologically active form of folate and it is essential for the formation of SAM, the universal methyl donor, by the regeneration of methionine from Hcy (see Figure 1). SAM is a powerful methylating agent in several biosynthetic reactions, and its methyl group is the most reactive in the one-carbon cycle (Nelson, 2017). Moreover, and remarkably, 5-methyl-THF is the only folate form able to cross the blood-brain barrier (Ducker and Rabinowitz, 2017). It is well absorbed in the intestinal tract and its bioavailability is not influenced by additional enzymatic steps (Pope et al., 2019). 5-methyl-THF has been made available for human administration only since 2000 (Venn et al., 2002) and it is still poorly used. One of the reasons for this is because it costs more than folic acid even though it is has been proposed to possibly be superior than folic acid in allowing the bypass of the pathway needed to generate 5-methyl-THF from folic acid, for example when the methylation step of THF is impaired in the carriers of the *MTHFR* C6777 polymorphism (Greenberg and Bell, 2011).

The use of 5-methyl-THF has more recently been proposed for the treatment of CFD (Li et al., 2008;Knowles et al., 2016) and is considered to be the most efficient way to normalize CSF 5-methyl-THF concentrations (Pope et al., 2019). Although 5-methyl-THF plasma concentration is not decreased in the subjects with DS that we have investigated, from the above discussion it appears to be the best way to restore the THF deficit. In addition, considering that in subjects with CFD the peripheral blood 5-methyl-THF levels may be normal while they are low in the CSF (Pope et al., 2019), a similar condition might be present in DS. Even if a draw of CSF only for research appears unjustified to us, we plan to study 5-methyl-THF levels in samples available following neurosurgery in subjects with DS.

For all these reasons, we propose 5-methyl-THF as the best candidate for a clinical trial that aims to improve cognitive status of subjects with DS. Nevertheless, we advise against administrating 5-methyl-THF until the effective dosage able to correct the altered values is identified in a pilot study.

## Acknowledgements

We wish to sincerely thank all children, families, colleagues, and students participating in our study on trisomy 21. Our heartfelt thanks to all foundations, associations, families, companies, and friends supporting our research by donations and by promoting fund raising initiatives. A complete list of the donors will be released.

