## Supplementary Figure 1 for "One-carbon pathway metabolites are altered in the plasma of subjects with Down syndrome: relation to chromosomal dosage"

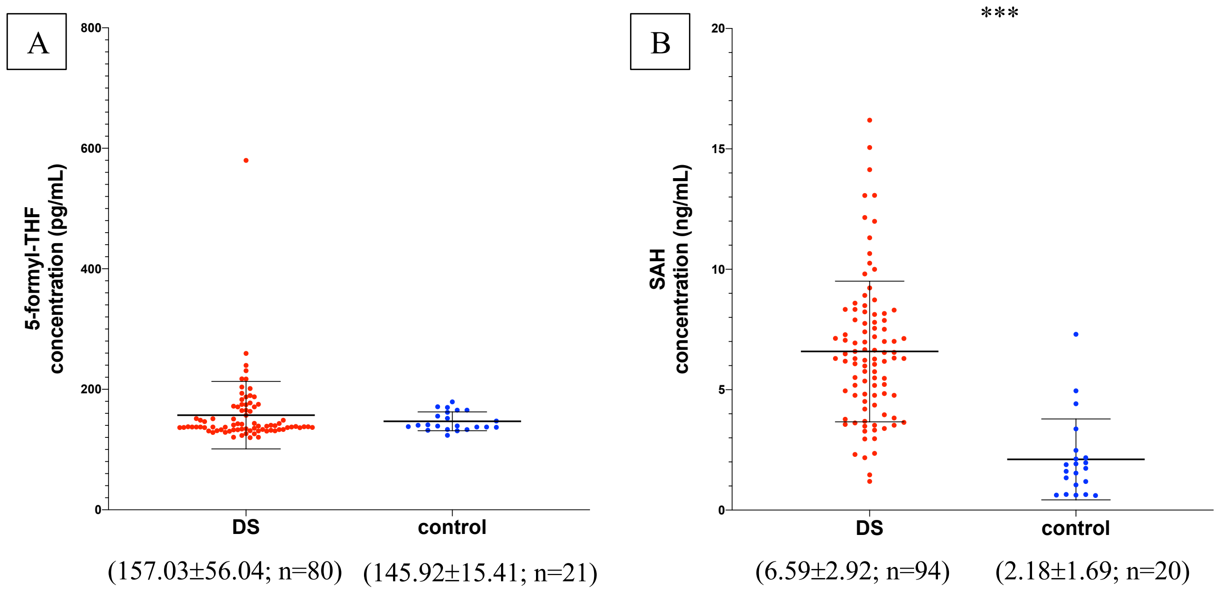


**Supplementary Figure 1. 5-formyl-THF and SAH concentrations in subjects with DS and normal control subjects including strong outliers.** The graphs report 5-formyl-THF plasma concentrations of each subject in the study. On the x-axis there is the subdivision of subjects in DS and control groups. Subjects with DS are represented like red dots and normal control subjects are represented like blue dots. On the y-axis the concentration of the metabolite in ng/mL or pg/mL is reported. The middle black lines indicate the mean concentration values for each group and the external black lines indicate standard deviation (SD) values. The mean concentration, SD values and the number of subjects (n) are reported below the graph for DS and control groups. The asterisks above the graph indicate the level of statistical significance (*=p<0.05; **=p<0.005; ***=p<0.0005). The graph was created with GraphPad Prism software v.6.0 (San Diego, CA).
