## Supplementary Figure 2 for "One-carbon pathway metabolites are altered in the plasma of subjects with Down syndrome: relation to chromosomal dosage"

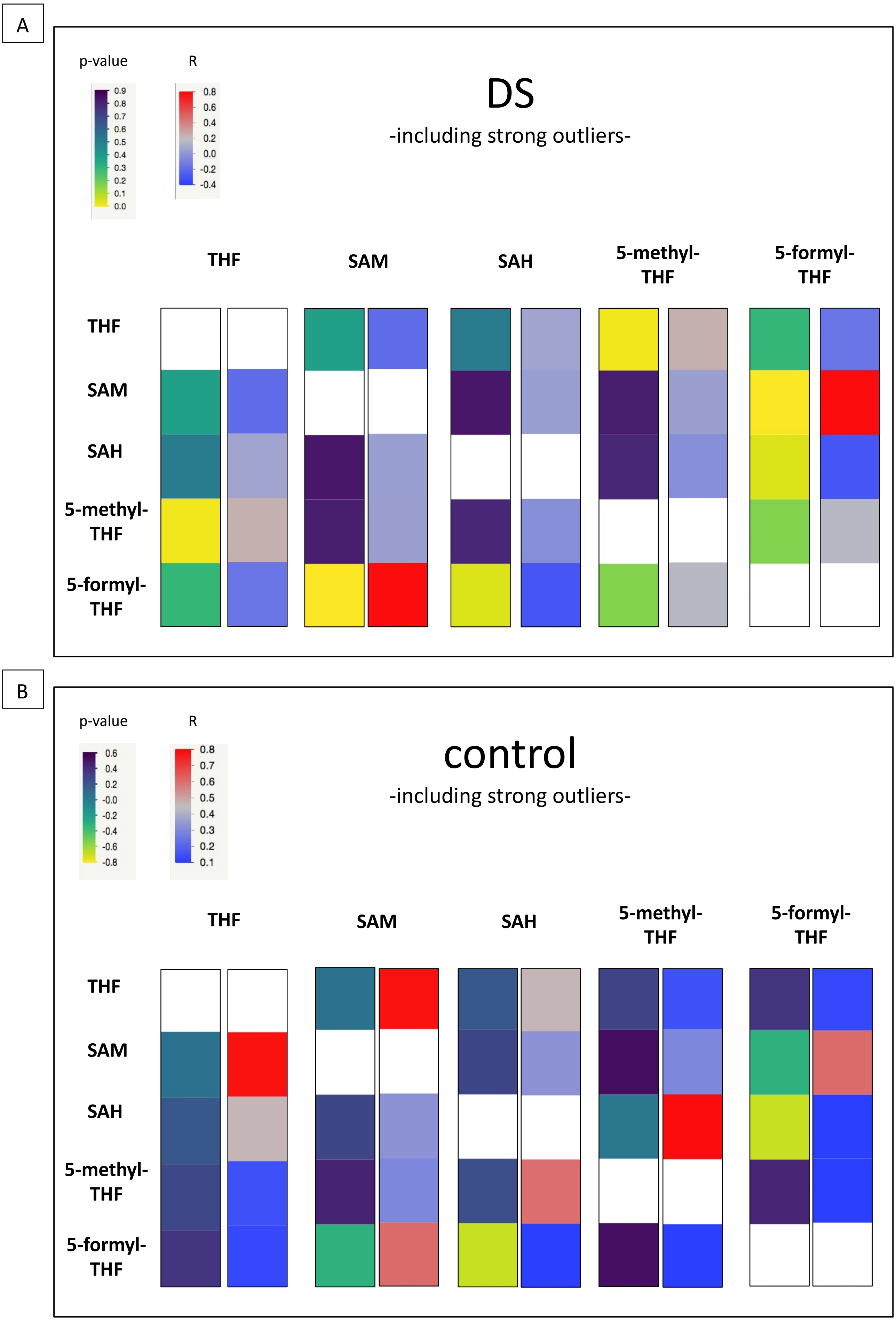
A
p-value
R
DS
-including strong outliers-
THF
SAM
SAH
5-methyl-THF
5-formyl-THF
B
p-value
R
control
-including strong outliers-
THF
SAM
SAH
5-methyl-THF
5-formyl-THF

**Supplementary Figure 2. Heat Map figure of bivariate correlation between levels of each metabolite and levels of all the other metabolites including strong outliers.** Supplementary Figure 2A presents bivariate correlation in DS group and Supplementary Figure 2B presents bivariate correlation in control group (for complete data see Supplementary Table 10). At the top left of the figures the color codes for Pearson correlation coefficient (r) and two tailed-significance (p-value) are reported. For each metabolite indicated above, two bars are shown: the first represents p-values and the second r values. Every bar is divided in five segments corresponding to the five correlations with other metabolites.
