## Supplementary Dataset for "One-carbon pathway metabolites are altered in the plasma of subjects with Down syndrome: relation to chromosomal dosage"

**Supplementary Dataset 1a. Dataset of subjects with DS selected in this study.** Each subject is identified by a code ("subject code") constituted by subject diagnosis that is Down syndrome ("DS") and a unique serial number. Each plasma sample is identified by a code ("sample code") constituted by: subject diagnosis ("DS"), the origin of the sample that is plasma ("P") and a unique serial number. For each subject sex ("M" for male or "F" for female) and age at the time of blood collection is reported. Moreover, fasting status at the time of collection is reported ("YES" if the subject was fasting or "NO" if the subject was not fasting). The "Sibling code sample" column shows the sample code of a subject enrolled in the control group who is familiar with subject with DS. The "Disease" and "drugs" columns show whether subject had pathologies or assumed drugs at the time of blood collection. The last columns report the concentration levels of metabolites obtained by ELISA assays (THF, 5-methyl-THF, 5-formyl-THF, SAH and SAM) and folic acid, vitamin B12 and homocysteine. The strong outliers among concentration values are reported in red. Below the main table, the mean age of subjects and the corresponding standard deviation (SD) are reported.

| Subject code | Sample code | Sex | Age | Fasting | Sibling code sample | Disease | Drugs | THF concentration (ng/mL) | 5-methyl-THF concentration (ng/mL) | 5-formyl-THF concentration (pg/mL) | SAH concentration (ng/mL) | SAM concentration (ng/mL) | Folic Acid concentration (ng/mL) | Vitamin B12 concentration (pg/mL) | Homocysteine concentration (µmol/L) |
|---|---|---|---|---|---|---|---|---|---|---|---|---|---|---|---|
| DS059 | DSP19489 | F | 29,2 | YES | | Basedow's disease | Tapazole 2.5mg | 5,631 | 43,403 | 217,363 | 7,125 | | 13,4 | 376 | 9,6 |
| DS097 | DSP19529 | M | 13,0 | YES | nP19727 | Hypothyroidism | Eutirox 31.25mg | | 28,538 | 132,577 | | | 6,7 | 327 | 12,4 |
| DS004 | DSP19596 | M | 20,3 | YES | | Hypothyroidism | Eutirox 75mg | | 44,944 | | | | 6,1 | 202 | |
| DS088 | DSP19689 | F | 13,0 | YES | nP19691 | NO | NO | | 28,021 | | | | | 164 | 5,8 |
| DS102 | DSP19757 | M | 13,0 | NO | | Hypothyroidism | Eutirox 25mg | 64,294 | | | | | 5,7 | 587 | 8,5 |
| DS135 | DSP19759 | F | 16,6 | NO | | Hypothyroidism | Eutirox | | 34,871 | 189,490 | | | | 588 | 6,1 |
| DS062 | DSP19761 | M | 24,0 | YES | | NO | Movicol | | 47,969 | | | | 7,5 | 378 | 9,2 |
| DS140 | DSP19770 | M | 11,6 | NO | | NO | NO | | 45,243 | 133,341 | | | | 390 | 8,3 |
| DS041 | DSP19776 | M | 9,1 | NO | | NO | NO | | 51,957 | | | | 11,9 | 557 | 8,3 |
| DS109 | DSP19841 | M | 7,0 | YES | | Celiac disease | Citogenex | 11,840 | | | | | 10,7 | 329 | 8,3 |
| DS021 | DSP19843 | F | 14,0 | YES | nP19461 | NO | NO | | 53,202 | | | | 12,0 | 595 | |
| DS128 | DSP19873 | M | 6,1 | NO | | NO | Vitamin D | 10,215 | 60,025 | 230,856 | 2,174 | | 7,7 | 501 | 11,2 |
| DS016 | DSP19891 | M | 11,1 | YES | | NO | Growth Hormone (Enantone) every 3 month | | 41,981 | | | | 10 | 207 | 10,10 |
| DS026 | DSP19895 | F | 23,0 | YES | nP19474; nP19487 | NO | Eutirox | 8,782 | 59,101 | 170,896 | 3,389 | | 7,9 | 331 | 14,1 |
| DS027 | DSP19897 | M | 12,0 | YES | nP19481 | NO | NO | 57,962 | | | | | 8,0 | 269 | 10,8 |
| DS158 | DSP19901 | M | 3,1 | YES | nP19902 | NO | NO | 11,307 | | | | | | 481 | 7,8 |
| DS045 | DSP19913 | M | 5,8 | YES | | Hypothyroidism | Eutirox | 35,073 | 43,668 | | | | 7,3 | 392 | 7,3 |
| DS162 | DSP19916 | F | 15,9 | NO | | NO | NO | 75,757 | 51,827 | | | | 4,4 | 142 | 11,2 |
| DS092 | DSP19939 | F | 18,1 | NO | | Celiac disease | NO | 107,770 | 42,503 | 137,301 | | | 4,7 | 207 | 5,6 |
| DS130 | DSP19944 | M | 14,0 | YES | nP19946 | NO | NO | | 48,790 | | | | 3,6 | 183 | 15,7 |
| DS170 | DSP19948 | F | 16,0 | YES | nP20246 | NO | NO | 17,192 | 48,491 | | 11,994 | | 3,5 | 240 | 13,4 |
| DS078 | DSP19952 | M | 10,0 | NO | | NO | NO | 56,646 | 49,594 | 138,875 | 7,000 | | 6,6 | 301 | 14,4 |
| DS172 | DSP19956 | M | 25,0 | YES | | Hypothyroidism | Eutirox | 51,777 | 44,147 | 175,164 | 6,543 | | 4,5 | 143 | 23,1 |
| DS087 | DSP19957 | M | 5,1 | NO | | Hypothyroidism | Eutirox | 39,090 | 70,575 | | 5,474 | | 4,4 | 287 | 5,2 |
| DS121 | DSP19966 | M | 30,0 | YES | | Hypothyroidism | Eutirox | | 48,611 | | | | 5,1 | 314 | 19,3 |
| DS108 | DSP19970 | F | 5,2 | NO | | Hypothyroidism | Eutirox; iron | 35,389 | 59,361 | | 3,488 | | | 1501 | 2,1 |
| DS174 | DSP19971 | F | 11,6 | NO | | NO | NO | | 34,246 | 126,324 | 1,463 | | 5,3 | 292 | 6,8 |
| DS051 | DSP19981 | F | 13,0 | YES | | NO | NO | | 34,825 | | | | 6,3 | 248 | 8,0 |
| DS077 | DSP19989 | M | 6,1 | NO | | NO | Captopril | | 57,399 | 148,754 | 14,136 | | 20,6 | 478 | 4,1 |
| DS129 | DSP19993 | M | 5,0 | NO | | Hypothyroidism | Eutirox | 48,118 | | | 5,220 | | 9,3 | 524 | 4,5 |
| DS036 | DSP19997 | F | 12,0 | YES | | NO | NO | | 37,345 | 134,494 | | | 7,4 | 479 | 9,6 |
| DS177 | DSP20001 | M | 10,0 | YES | | NO | Lamotrigina | | | 130,635 | | | 7,6 | 320 | 4,9 |
| DS105 | DSP20003 | M | 4,1 | NO | | NO | NO | 31,018 | | | | | 6,4 | 279 | 5,6 |
| DS037 | DSP20008 | M | 28,0 | NO | | Celiac disease | Vitamin D | 59,422 | 58,060 | | 6,490 | | 6,5 | 492 | 13,4 |
| DS179 | DSP20013 | M | 3,0 | YES | | Hypothyroidism | Tirosint | | 45,443 | | | | | | 3,9 |
| DS095 | DSP20018 | M | 10,0 | NO | | NO | NO | 51,801 | 58,642 | | 8,592 | | 6,1 | 150 | 8,4 |
| DS076 | DSP20021 | M | 5,1 | NO | | NO | NO | 20,160 | | | 2,957 | | 9,8 | 558 | 8,1 |
| DS053 | DSP20029 | F | 8,0 | YES | | NO | NO | | 39,588 | 133,360 | | | 8,2 | 352 | 6,3 |
| DS181 | DSP20030 | F | 3,5 | NO | | Hypothyroidism | Eutirox 25mg | 40,902 | | | | | 9,2 | 574 | 7,2 |
| DS089 | DSP20034 | F | 22,9 | YES | | NO | NO | 96,489 | 57,351 | 138,228 | 7,052 | | 17,4 | 208 | 7,0 |
| DS183 | DSP20037 | F | 7,1 | NO | | NO | NO | 56,160 | | 171,857 | 5,488 | | 4,7 | 369 | 8,8 |
| DS058 | DSP20047 | M | 13,0 | YES | | NO | NO | 16,329 | 56,584 | | 7,510 | | 6,5 | 159 | 8,4 |
| DS003 | DSP20062 | F | 20,0 | YES | | NO | NO | 25,456 | 65,085 | | 9,806 | | 3,6 | 188 | 10,00 |
| DS186 | DSP20066 | F | 3,5 | N/A | | NO | NO | | 56,797 | | | | 9,2 | 346 | 8,5 |
| DS137 | DSP20068 | M | 6,1 | NO | | NO | Dibase; Singulair | | 40,311 | | | | 4,2 | 294 | |
| DS010 | DSP20075 | F | 37,9 | YES | | Hypothyroidism | Eutirox; Cipralex | | 49,293 | | | | 8,7 | 326 | |
| DS069 | DSP20077 | F | 12,0 | NO | | Hypothyroidism; Celiac disease | Eutirox | 21,260 | 47,185 | 136,529 | 16,194 | | 6,8 | 342 | |
| DS136 | DSP20078 | M | 13,0 | YES | | NO | NO | 27,966 | 60,453 | 156,928 | 3,643 | | 5,7 | 232 | |
| DS104 | DSP20080 | M | 15,0 | NO | | NO | NO | 33,076 | 62,786 | 136,813 | 9,225 | 9,152 | 12,5 | 345 | |
| DS009 | DSP20082 | M | 20,0 | YES | | Hypothyroidism | Eutirox 125mg | 130,213 | 53,813 | 217,168 | 6,644 | | 13,4 | 148 | |
| DS187 | DSP20085 | M | 3,1 | NO | | NO | Ferritin | 11,014 | | 120,536 | 4,770 | | | | |
| DS023 | DSP20086 | M | 15,0 | YES | nP19568; nP19567 | NO | NO | 44,054 | 56,468 | 137,955 | 6,553 | 9,090 | 5,2 | 1007 | |

| | | | | | | | | | | | | | | |
|---|---|---|---|---|---|---|---|---|---|---|---|---|---|---|
| DS101 | DSP20089 | F | 14,2 | NO | | NO | Cortisone, azathioprine, furosemide, amlodipine, colecalcium, Pantorc | | | | 3,561 | | 4,8 | 277 |
| DS143 | DSP20091 | M | 9,0 | YES | | Hypothyroidism | Eutirox | 21,128 | 36,020 | 137,060 | 8,128 | 6,970 | 4,9 | 443 |
| DS144 | DSP20092 | M | 7,0 | NO | | NO | NO | 125,646 | 61,665 | 146,494 | | | 4,9 | 242 |
| DS188 | DSP20095 | F | 3,1 | YES | | NO | Fluispiral; Equazen; Montelukast; Axil | 38,303 | 64,129 | | 5,186 | | 9,3 | 435 |
| DS011 | DSP20096 | F | 13,0 | YES | | Mild hypothyroidism | NO | 13,534 | 43,639 | 136,582 | 7,552 | 5,610 | 6,4 | 284 |
| DS156 | DSP20098 | F | 8,0 | NO | | NO | Probiotics | | 47,102 | | | | 8,6 | 406 |
| DS191 | DSP20103 | M | 3,1 | NO | | NO | NO | 19,296 | | | 4,841 | | 8,4 | 356 |
| DS146 | DSP20104 | M | 4,0 | NO | | NO | NO | 6,908 | 55,358 | | | | 10,3 | 144 |
| DS147 | DSP20107 | F | 12,0 | YES | | Celiac disease | NO | | 49,327 | 136,427 | | | 5,0 | 349 |
| DS106 | DSP20110 | F | 6,0 | YES | nP19812 | NO | NO | 98,795 | 63,362 | 148,760 | 5,182 | 11,653 | 5,8 | 331 |
| DS193 | DSP20112 | M | 3,1 | NO | | NO | NO | 30,352 | | | 3,959 | | 6,0 | 798 |
| DS194 | DSP20113 | M | 11,6 | NO | | NO | NO | 50,357 | 49,964 | | 7,900 | | 6,5 | 263 |
| DS072 | DSP20115 | M | 7,0 | YES | | Hypothyroidism | Eutirox; Singulair | 18,738 | | 165,173 | 3,323 | | 9,5 | 348 |
| DS012 | DSP20117 | F | 21,1 | YES | | Celiac disease | NO | 61,048 | 43,393 | 137,774 | | | 3,6 | 212 |
| DS151 | DSP20121 | M | 5,2 | NO | | Hypothyroidism | Eutirox | 67,964 | 46,410 | 138,767 | | | 5,0 | 357 |
| DS071 | DSP20123 | F | 24,0 | YES | | Hypothyroidism | Eutirox | 89,390 | 54,138 | | 6,315 | | 4,2 | 218 |
| DS155 | DSP20124 | M | 13,0 | NO | | Hypothyroidism | Eutirox | | 54,313 | 131,926 | | | 12,8 | 338 |
| DS044 | DSP20126 | F | 7,0 | NO | nP19576 | NO | Augmentin; Broncovaleas; Prontidal | | 37,524 | | | | 9,1 | 354 |
| DS196 | DSP20127 | F | 5,6 | NO | | NO | Antihistamine | 61,230 | 65,694 | | 4,818 | 6,964 | 15,7 | 350 |
| DS197 | DSP20128 | M | 13,5 | NO | | NO | NO | 3,820 | 48,937 | 239,589 | 6,076 | | 7,0 | 217 |
| DS199 | DSP20132 | M | 3,2 | NO | | NO | NO | 12,193 | 49,457 | 126,003 | 5,507 | | 10,4 | 390 |
| DS148 | DSP20136 | F | 5,4 | NO | nP19821; nP19822 | NO | NO | 50,071 | 69,121 | | 3,824 | 7,356 | 4,7 | 485 |
| DS017 | DSP20139 | M | 10,0 | NO | | Hypothyroidism | Eutirox; Equazen; Magnesium; Xizal | 43,246 | 65,368 | 143,500 | 13,070 | 8,338 | 6,9 | 368 |
| DS154 | DSP20140 | F | 8,0 | YES | | Hypothyroidism | Eutirox | 43,671 | 46,525 | 137,905 | 13,064 | | 7,1 | 363 |
| DS018 | DSP20143 | M | 12,0 | YES | nP19450 | Hypercholesterolemia | Green tea extract; Omega 3 and 6 | 55,129 | | | | | 13,7 | 210 |
| DS107 | DSP20144 | M | 10,0 | YES | | Hypothyroidism | Eutirox | 29,691 | 53,626 | | 8,916 | | 5,8 | 561 |
| DS150 | DSP20146 | F | 26,0 | YES | | Hypothyroidism | Eutirox 75mg | | 43,510 | 132,946 | | | 11,1 | 235 |
| DS201 | DSP20150 | F | 10,1 | NO | | NO | NO | 51,461 | 55,188 | 137,467 | 10,252 | | 10,3 | 520 |
| DS103 | DSP20155 | F | 17,0 | YES | | NO | NO | 51,364 | 47,918 | | 6,662 | 7,243 | 10,7 | 351 |
| DS141 | DSP20157 | M | 9,0 | NO | | Hypothyroidism | Eutirox | | 29,930 | 130,740 | | | 4,0 | 386 |
| DS152 | DSP20158 | F | 8,9 | NO | | NO | NO | | 45,517 | 136,721 | | | 4,1 | 316 |
| DS157 | DSP20161 | M | 8,0 | NO | | NO | NO | 80,882 | 51,766 | | 6,226 | | 6,9 | 386 |
| DS125 | DSP20162 | M | 11,6 | NO | | NO | NO | | 48,083 | 133,053 | | | 10,8 | 329 |
| DS110 | DSP20164 | F | 7,9 | YES | | NO | Captopril; Gaviscon | 68,395 | 58,741 | 137,490 | 7,877 | 6,395 | 10,2 | 443 |
| DS204 | DSP20168 | M | 11,7 | NO | | NO | NO | | 54,832 | 134,111 | | | 7,3 | 211 |
| DS024 | DSP20174 | M | 23,0 | YES | nP19467 | NO | Relax Cal (supplement) | | | | 4,197 | | 3,2 | 272 |
| DS081 | DSP20178 | F | 11,0 | NO | | NO | NO | 26,746 | 46,873 | | | | 6,1 | 341 |
| DS112 | DSP20183 | F | 27,1 | YES | | Hypothyroidism | Eutirox | 21,509 | 56,897 | | 8,727 | | 7,8 | 324 |
| DS111 | DSP20187 | M | 13,0 | YES | nP19606 | Hypothyroidism; Celiac disease | Eutirox | 67,401 | 54,444 | 140,809 | | | 3,5 | 145 |
| DS113 | DSP20189 | M | 14,0 | YES | | Hypothyroidism | Eutirox | 25,200 | 49,493 | 142,854 | 3,774 | | 5,6 | 106 |
| DS161 | DSP20191 | M | 7,0 | NO | | NO | Nasal spray | 137,922 | 65,265 | 170,757 | 3,623 | | | |
| DS207 | DSP20192 | F | 7,4 | YES | | NO | NO | | | 203,724 | 6,275 | | 7,4 | 424 |
| DS079 | DSP20194 | F | 16,0 | YES | nP19494 | NO | NO | 83,292 | 46,981 | 143,278 | 6,176 | | 4,5 | 180 |
| DS047 | DSP20195 | F | 14,0 | NO | | NO | Enapren | 7,168 | 36,386 | 187,206 | 3,278 | | 5,2 | 467 |
| DS114 | DSP20200 | F | 10,0 | NO | | Hypothyroidism | Eutirox | 24,843 | 46,730 | | 7,132 | 13,200 | 6,7 | 301 |
| DS208 | DSP20203 | M | 4,6 | YES | | NO | NO | 6,666 | 39,042 | | 4,768 | | 5,4 | 282 |
| DS034 | DSP20205 | M | 7,2 | NO | | NO | NO | 59,514 | 58,937 | | | | 3,7 | 226 |
| DS209 | DSP20207 | M | 9,1 | NO | | NO | Risperidone | 22,031 | 49,916 | | 10,656 | | 9 | 345 |
| DS210 | DSP20209 | M | 14,5 | NO | | Celiac disease | NO | | 46,614 | 128,677 | 1,194 | | | |
| DS211 | DSP20212 | M | 13,3 | NO | | NO | NO | | 44,851 | 132,929 | | | | |
| DS173 | DSP20214 | F | 17,1 | YES | | Hypothyroidism | Eutirox; Clobazam | 57,911 | 48,808 | | 6,289 | | 3,7 | 388 |
| DS169 | DSP20216 | F | 5,1 | YES | | Hypothyroidism | Tirosint 7 drops | | 45,818 | 136,167 | | | 10,3 | 197 |
| DS212 | DSP20220 | M | 24,7 | NO | | Hypothyroidism | Eutirox 100mg | 61,941 | 45,681 | 140,262 | 15,055 | | 7,7 | 223 |
| DS214 | DSP20224 | M | 6,2 | YES | | Hypothyroidism | Eutirox | | 43,671 | | | | 5,7 | 175 |
| DS167 | DSP20230 | F | 13,0 | NO | nP20228; nP20231 | NO | Zimox | 22,008 | 51,766 | | 6,298 | | 4,9 | 792 |
| DS166 | DSP20232 | F | 15,1 | YES | | Hypothyroidism | Eutirox 25mg | | 57,717 | | | | 2,6 | 335 |
| DS178 | DSP20235 | M | 4,0 | YES | | Celiac disease (not already diagnosed) | Bactrim | 3,894 | 36,672 | | 5,751 | | 10,8 | 343 |
| DS215 | DSP20236 | M | 20,3 | YES | | NO | NO | 34,835 | 44,194 | 139,671 | 8,333 | | 8,5 | 179 |
| DS042 | DSP20237 | F | 9,0 | NO | | NO | NO | | 48,270 | 131,038 | | | 3,7 | 312 |
| DS117 | DSP20240 | M | 11,0 | YES | | NO | NO | | 38,685 | 131,337 | | | 4,4 | 480 |
| DS085 | DSP20242 | M | 17,1 | YES | | NO | NO | 21,802 | 62,128 | | 7,402 | 10,801 | 4,7 | 315 |
| DS094 | DSP20255 | F | 26,1 | YES | | Hypothyroidism | Eutirox | 3,948 | 38,655 | 177,292 | 2,358 | | 8,0 | 338 |

| | | | | | | | | | | | | | | |
|---|---|---|---|---|---|---|---|---|---|---|---|---|---|---|
| DS171 | DSP20257 | M | 16,1 | YES | nP19968 | NO | NO | 36,268 | 54,699 | 137,314 | | 8,064 | 4,1 | 127 |
| DS219 | DSP20264 | M | 13,1 | NO | | NO | NO | 24,172 | 38,255 | | 7,202 | 10,788 | 9,9 | 149 |
| DS050 | DSP20265 | F | 14,1 | YES | | Hypothyroidism | Eutirox | | 50,993 | 193,446 | | | 12,3 | 409 |
| DS013 | DSP20267 | F | 8,0 | YES | | NO | Bactolis; Aircart | 50,057 | 60,403 | 201,372 | | 22,056 | 9,9 | 564 |
| DS124 | DSP20271 | F | 15,1 | NO | nP20273 | Celiac disease | NO | 71,270 | 48,442 | 142,161 | 8,335 | | 10,0 | 323 |
| DS220 | DSP20274 | F | 3,2 | NO | | NO | NO | 21,747 | 58,414 | 187,636 | 6,185 | | | 439 |
| DS221 | DSP20275 | F | 14,9 | NO | | Autoimmune thyroiditis | Eutirox | 13,383 | 46,850 | | 3,527 | | 15,0 | 287 |
| DS222 | DSP20279 | M | 3,1 | NO | | Strawberry allergy | NO | | 49,104 | 123,434 | 7,800 | | | 600 |
| DS093 | DSP20282 | M | 12,9 | NO | nP20284 | NO | NO | 5,172 | 45,096 | | 2,965 | | 8,2 | 395 |
| DS134 | DSP20286 | M | 6,0 | NO | nP19737 | NO | Clavulin | 32,225 | 67,879 | 174,910 | | | | |
| DS116 | DSP20288 | M | 10,3 | NO | | NO | NO | | 32,781 | | | | 10,3 | 128 |
| DS223 | DSP20290 | M | 6,0 | NO | | Hypothyroidism (not already diagnosed) | NO | 53,453 | 56,476 | 151,387 | 7,752 | | | |
| DS086 | DSP20294 | F | 10,0 | NO | | NO | NO | 88,622 | 56,064 | 163,255 | | 8,042 | 5,1 | 431 |
| DS091 | DSP20296 | M | 14,0 | NO | | NO | Flubason; Zeta Spray; Lipikar AP + | 89,199 | 39,847 | | | | 4,6 | 274 |
| DS122 | DSP20299 | M | 6,0 | NO | nP19660 | Hypothyroidism | Eutirox | 99,118 | 57,668 | | 2,309 | 4,049 | 13,8 | 354 |
| DS090 | DSP20302 | M | 31,0 | YES | | NO | Zolof 50 mg; Serpinol drops | 59,891 | 58,191 | 150,812 | 8,168 | | 4,7 | 164 |
| DS149 | DSP20312 | F | 7,6 | YES | | N/A | N/A | 16,219 | 41,350 | 259,457 | 3,691 | | | 237 |
| DS115 | DSP20316 | F | 8,0 | YES | | NO | NO | | 51,862 | 137,485 | | | | |
| DS068 | DSP20318 | M | 5,9 | YES | | Hypothyroidism | Eutirox | | 53,812 | | | | 7,3 | 345 |
| DS035 | DSP20327 | F | 16,0 | N/A | | N/A | N/A | 23,172 | 58,497 | | 7,284 | 18,197 | 8,0 | 415 |
| DS067 | DSP20329 | M | 6,8 | YES | | NO | Amoxicillin | | 59,744 | | | | 4,6 | 307 |
| DS224 | DSP20330 | M | 6,2 | NO | | Celiac disease (not already diagnosed) | Pediason | 7,619 | | 174,352 | 5,984 | | 6,9 | 298 |
| DS098 | DSP20331 | F | 21,0 | YES | nP19533 | Hypothyroidism | Eutirox | 26,137 | 66,836 | 182,990 | 6,982 | | 2,5 | 159 |
| DS176 | DSP20333 | F | 12,0 | YES | | Celiac disease | Vitamin D | 14,328 | 61,190 | | 4,359 | 11,284 | 9,3 | 259 |
| DS145 | DSP20337 | M | 9,0 | YES | | NO | NO | | 51,655 | 130,851 | | | 8,3 | 275 |
| DS056 | DSP20339 | F | 9,0 | NO | | Hypothyroidism | Eutirox | | 62,213 | | | | 19,1 | 609 |
| DS180 | DSP20347 | F | 7,0 | YES | | NO | NO | | 51,950 | | | | 10,4 | 294 |
| DS226 | DSP20350 | M | 3,0 | YES | | NO | Vitamin B12 | 30,047 | 59,036 | 128,335 | 4,954 | | 12,3 | 407 |
| DS131 | DSP20351 | M | 8,0 | YES | nP19718 | NO | NO | | 52,887 | 130,470 | | | 10,3 | 276 |
| DS182 | DSP20353 | M | 15,7 | NO | | NO | NO | 31,932 | 56,506 | | 12,153 | 9,877 | 6,9 | 344 |
| DS198 | DSP20359 | M | 14,1 | NO | | NO | Resveratrol | | 26,010 | 579,917 | 11,308 | | 6,1 | 225 |
| DS142 | DSP20360 | M | 5,0 | N/A | nP19783 | N/A | N/A | 62,977 | | | 10,002 | | | |
| DS227 | DSP20361 | M | 10,0 | NO | | NO | NO | | 47,426 | 131,216 | | | 8,5 | 374 |
| DS096 | DSP20363 | M | 20,1 | YES | | NO | NO | 32,461 | 64,980 | | 5,761 | | 7,8 | 377 |
| DS039 | DSP20365 | M | 12,0 | NO | | NO | NO | | 57,320 | 135,292 | | | 7,5 | 326 |
| DS217 | DSP20366 | M | 4,0 | NO | | NO | NO | 16,447 | 50,663 | 120,570 | 6,943 | | | 495 |
| DS228 | DSP20367 | F | 3,5 | NO | | NO | NO | 68,730 | | | | | 7,8 | 163 |
| DS229 | DSP20368 | M | 4,0 | NO | | Hypothyroidism | Bactrim; Eutirox 75mg | 19,814 | | | 7,011 | | | 172 |
| DS100 | DSP20371 | F | 13,0 | NO | | Celiac disease | Lasox 30 mg | 4,297 | 45,858 | | 6,299 | | 9,1 | 358 |
| DS118 | DSP20373 | F | 6,1 | YES | nP19645; nP19647 | NO | Fucus; Otichil; Formistin; Bactoblis | 40,764 | 58,934 | 164,691 | 4,511 | | | 216 |
| DS057 | DSP20374 | M | 11,0 | NO | | NO | NO | 67,636 | 54,966 | | 5,357 | 15,059 | 9,7 | 178 |
| DS066 | DSP20376 | M | 15,0 | YES | | Hypothyroidism | Eutirox | 56,851 | 52,173 | | | | 3,8 | 152 |
| DS230 | DSP20379 | M | 15,0 | NO | | NO | Quercetin; Resveratrol | 8,307 | 24,217 | 119,681 | 8,494 | 10,025 | 13,2 | 225 |
| DS231 | DSP20380 | M | 4,8 | NO | | NO | NO | 56,119 | 50,337 | | 8,235 | | 7,3 | 455 |
| DS159 | DSP20381 | F | 14,3 | NO | | Celiac disease | NO | 26,377 | 58,197 | | 6,055 | 10,207 | 4,8 | 345 |
| DS059 | DSP20383 | F | 11,0 | YES | | NO | Eutirox; Fluifort | 30,415 | 49,616 | | 6,592 | 16,963 | 10,0 | 115 |
| DS184 | DSP20384 | F | 5,0 | NO | | NO | NO | | 46,473 | 151,279 | | | 8,9 | 293 |
| DS232 | DSP20386 | F | 4,3 | NO | | NO | NO | | | | 4,768 | | | 551 |
| DS233 | DSP20387 | M | 4,3 | NO | | NO | NO | | | | 8,307 | | 20,6 | 926 |
| DS234 | DSP20388 | M | 9,7 | NO | | NO | Prenole; Vitamin D; Herbs for sleep (passionflower and valerian) | 21,316 | 35,123 | | 3,528 | | 5,7 | 560 |

| | |
|---|---|
| **Mean age** | 11,548 |
| **SD** | 6,690 |

**Supplementary Dataset 1b. Dataset of normal control subjects selected in this study.** Each subject is identified by a code ("subject code") constituted by subject diagnosis that is normal control ("n") and a unique serial number. Each plasma sample is identified by a code ("sample code") constituted by: subject diagnosis ("n"), the origin of the sample that is plasma ("P") and a unique serial number. For each subject sex ("M" for male or "F" for female) and age at the time of blood collection is reported. Moreover, fasting status at the time of collection is reported ("YES" if the subject was fasting or "NO" if the subject was not fasting). The "Sibling code sample" column shows the sample code of a subject enrolled in DS group who is familiar with normal control subject. The "Disease" and "drugs" columns show whether subject had pathologies or assumed drugs at the time of blood collection. The last columns report the concentration levels of metabolites obtained by ELISA assays. The strong outliers among metabolite concentrations are reported in red. Below the main table, the mean age of subjects and the corresponding standard deviation (SD) are reported.

| Subject code | Sample code | Sex | Age | Fasting | Sibling code sample | Disease | Drugs | THF concentration (ng/mL) | 5-methyl-THF concentration (ng/mL) | 5-formyl-THF concentration (pg/mL) | SAH concentration (ng/mL) | SAM concentration (ng/mL) |
|---|---|---|---|---|---|---|---|---|---|---|---|---|
| n016 | nP19365 | F | 16,8 | YES | | NO | NO | | 63,983 | 179,270 | | |
| n017 | nP19367 | M | 15,2 | YES | | N/A | N/A | | | | 2,177 | |
| n020 | nP19415 | F | 18,1 | YES | | NO | NO | 10,514 | 32,621 | | | |
| n021 | nP19416 | F | 20,6 | YES | | NO | NO | | 42,912 | | | |
| n022 | nP19417 | F | 7,1 | YES | | NO | NO | | 44,614 | | | |
| n023 | nP19418 | M | 21,8 | YES | | NO | NO | | 33,870 | | | |
| n024 | nP19419 | F | 11,5 | YES | | NO | Be-total (4 days ago) | | 42,874 | | | |
| n025 | nP19420 | F | 15,3 | YES | | NO | NO | | | 137,984 | 2,483 | |
| n028 | nP19444 | M | 27,3 | YES | | NO | NO | 81,293 | | 169,948 | 1,736 | |
| n007 | nP19450 | M | 3,7 | YES | DSP20143 | NO | NO | | | | 3,371 | 10,338 |
| n029 | nP19454 | M | 23,9 | YES | | NO | Scalp hair supplement | 88,833 | 72,788 | | 1,887 | 19,670 |
| n030 | nP19458 | F | 25,8 | YES | | NO | Estrogen progesterone pills | 105,413 | 34,567 | 170,992 | | |
| n013 | nP19460 | M | 13,0 | YES | | NO | NO | 75,324 | 42,614 | 140,428 | | |
| n004 | nP19461 | M | 16,8 | YES | | NO | Fornistin, aircort | 58,910 | 61,173 | | | |
| n032 | nP19467 | F | 25,7 | YES | | Hypothyroidism; Thrombocytopenia | Eutirox (25 mcg/day); Estrogen progesterone pills | 38,347 | 66,260 | 155,354 | | |
| n033 | nP19469 | M | 31,5 | YES | | NO | NO | | | 123,569 | 4,953 | |
| n034 | nP19474 | M | 24,4 | YES | DSP19895 | NO | Flaminase | 105,321 | 39,754 | 147,229 | 1,611 | |
| n036 | nP19481 | F | 4,6 | YES | DSP19897 | NO | Rinotricina; Tachipirina; Grintuss; Augmentin | | | | 2,126 | |
| n038 | nP19487 | F | 28,4 | YES | DSP19895 | NO | NO | 33,419 | 55,548 | 136,999 | 7,303 | |
| n039 | nP19494 | F | 8,2 | YES | DSP20194 | NO | NO | 45,871 | | 131,988 | | |
| n041 | nP19533 | M | 14,3 | YES | DSP20331 | N/A | NO | 25,641 | 32,941 | | | |
| n043 | nP19567 | M | 15,6 | YES | DSP20086 | NO | NO | | 51,477 | | | |
| n042 | nP19568 | M | 8,7 | YES | DSP20086 | NO | NO | 24,122 | 41,499 | | | |
| n045 | nP19576 | F | 15,5 | YES | DSP20126 | N/A | N/A | | 27,694 | | | |
| n046 | nP19606 | F | 13,7 | YES | DSP20187 | NO | NO | 15,732 | | 138,989 | | |
| n047 | nP19629 | M | 16,9 | YES | | NO | Xaranel (2 months ago) | 24,653 | 26,911 | | | |
| n048 | nP19631 | F | 14,1 | YES | | NO | Nurofen; Antibiotic; Bentelan (17 days ago ) | 51,471 | | | | |
| n049 | nP19645 | M | 7,9 | YES | DSP20373 | NO | NO | | 34,550 | | | |
| n050 | nP19647 | M | 9,5 | YES | DSP20373 | NO | NO | 26,419 | 37,662 | | | |
| n009 | nP19653 | F | 20,7 | NO | | NO | Jasmin | 35,138 | 50,663 | | 0,609 | 4,768 |
| n052 | nP19660 | M | 5,6 | YES | DSP20299 | NO | NO | 43,610 | | | | |
| n053 | nP19691 | F | 13,0 | YES | DSP19689 | NO | NO | 15,433 | | | 1,188 | 5,029 |
| n054 | nP19718 | M | 3,3 | YES | DSP20351 | NO | NO | 8,562 | 33,138 | | | |
| n055 | nP19727 | M | 9,6 | YES | DSP19529 | NO | NO | 39,286 | | | | |
| n056 | nP19737 | F | 11,2 | YES | DSP20286 | NO | NO | 39,186 | 47,971 | | | |
| n057 | nP19783 | F | 7,7 | NO | DSP20360 | NO | NO | 53,634 | 31,458 | | | |
| n058 | nP19812 | M | 9,0 | YES | DSP20110 | NO | NO | 31,198 | | | 1,924 | 3,098 |
| n059 | nP19821 | M | 14,6 | NO | DSP20136 | NO | Oki (occasionally) | 75,358 | 39,431 | 151,777 | | |
| n060 | nP19822 | M | 9,4 | NO | DSP20136 | NO | NO | 65,340 | | | | |

| | | | | | | | | | | | | |
|---|---|---|---|---|---|---|---|---|---|---|---|---|
| n061 | nP19854 | F | 9,4 | NO | | NO | NO | 21,402 | | 133,184 | 1,969 | 9,006 |
| n062 | nP19856 | M | 7,5 | NO | | NO | NO | 11,887 | 39,500 | 133,261 | | |
| n063 | nP19902 | M | 7,0 | YES | DSP19901 | NO | NO | 35,037 | | | 1,047 | 4,174 |
| n070 | nP19946 | M | 19,0 | YES | DSP19944 | NO | NO | 105,553 | 43,243 | 137,483 | | |
| n071 | nP19968 | M | 9,0 | YES | DSP20257 | NO | NO | 85,425 | 63,204 | | 4,418 | |
| n072 | nP20045 | F | 17,3 | YES | | NO | NO | 85,198 | 49,554 | 139,144 | 1,342 | 5,182 |
| n073 | nP20228 | F | 10,5 | NO | DSP20230 | NO | Soltux (only tonight) | 73,459 | | 130,994 | | |
| n074 | nP20231 | F | 15,9 | NO | DSP20230 | NO | NO | 91,015 | 44,494 | 137,441 | | 3,707 |
| n075 | nP20246 | M | 12,0 | N/A | DSP19948 | NO | Nurofen | 91,917 | | | 1,537 | 24,744 |
| n076 | nP20273 | M | 24,0 | NO | DSP20271 | NO | NO | 85,733 | 40,641 | 140,939 | | |
| n077 | nP20284 | F | 14,7 | NO | DSP20282 | NO | NO | 88,089 | 64,797 | 161,857 | 0,651 | 5,337 |
| n080 | nP20341 | F | 13,8 | YES | | NO | Rynoclenil nasal spray | 110,515 | | | 0,649 | 2,239 |
| n079 | nP20343 | M | 15,7 | YES | | Hypothyroidism | NO | 102,354 | | | | 6,694 |
| n081 | nP20398 | M | 12,5 | NO | | NO | NO | 47,949 | 42,534 | | 0,627 | 11,115 |
| n082 | nP20399 | M | 16,7 | NO | | NO | NO | 115,554 | 65,604 | 165,433 | | 5,222 |

| | |
|---|---|
| **Mean age** | 14,534 |
| **SD** | 6,636 |
