## Supplementary Table 1 for "One-carbon pathway metabolites are altered in the plasma of subjects with Down syndrome: relation to chromosomal dosage"

**Supplementary Table 1. Analysis of THF ELISA assays**. Totally five THF ELISA assays were performed and each assay is shown in a different Excel sheet named Plate 1, Plate 2, Plate 3, Plate 4, Plate 5. The table "Standard" shows the two absorbance (O.D.) detected for each standard sample. In the "O.D. mean" column, the average O.D. is reported. The "%CV" column represent the Coefficient of Variability between the two O.D. measurment expressed as percentage. The "Concentration (ng/mL)" column report the known concentration of each standard sample supplied by ELISA kit, while the last column shows the transformation of standard concentration values in their logarithm. In order to build the standard curve, we plotted "O.D. mean (x)" of standards in the x-axis and "Log 10 Concentration (y)" of standards in the y-axis. The polynomial equation (y=a+bx+cx^2) reported in the graph was used to determine metabolites concentration of plasma samples ("Log 10 Concentration (y)"), using interpolation of O.D. mean values reported in "O.D. mean (x)" column in tables "DS" and "control". The final concentration of plasma samples is obtained by conversion of logarithmic function. The results are reported in "Concentration (ng/mL)" column in tables "DS" and "control". In tables "DS" and "control" the O.D. mean values which are higher or lower than the O.D. mean values of standard sample ranges are reported in red and these values were not considered for the statistical analysis.

| Standard | | | | | |
|---|---|---|---|---|---|
| **Samples** | **O.D.** | **O.D. mean (x)** | **%CV** | **Concentration (ng/mL)** | **Log10 Concentration (y)** |
| STD1 | 0,429<br>0,946 | 0,688 | 53,227 | 200 | 2,301 |
| STD2 | 0,696<br>1,341 | 1,018 | 44,736 | 100 | 2,000 |
| STD3 | 1,220<br>1,547 | 1,383 | 16,742 | 50 | 1,699 |
| STD4 | 1,195<br>1,693 | 1,444 | 24,346 | 25 | 1,398 |
| STD5 | 1,567<br>1,798 | 1,682 | 9,734 | 12,5 | 1,097 |
| STD6 | 1,734<br>1,993 | 1,863 | 9,852 | 6,25 | 0,796 |
| STD7 | 2,228<br>2,151 | 2,190 | 2,510 | 3,12 | 0,494 |
| STD8 | 2,348<br>2,446 | 2,397 | 2,895 | 0 | |

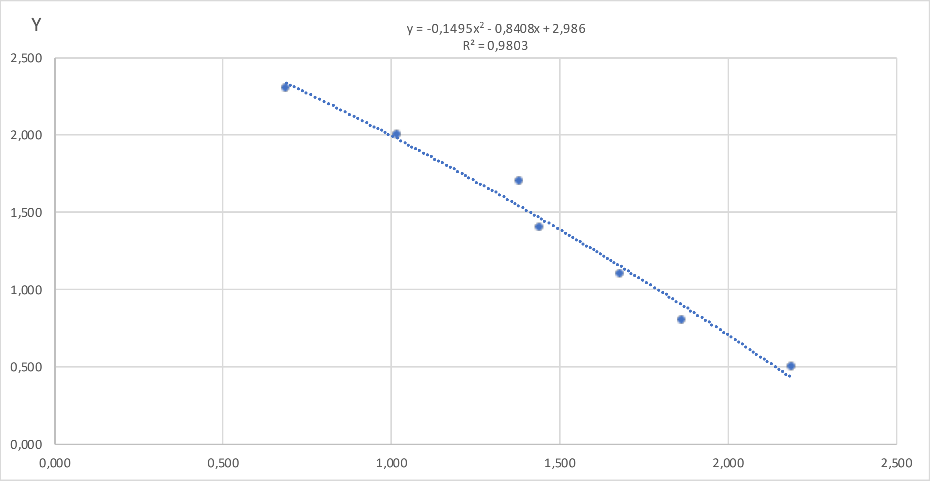


| y=a+bx+cx^2 | |
|---|---|
| a | 2,986 |
| b | -0,841 |
| c | -0,150 |
| R^2 | 0,980 |

| DS | | | | | |
|---|---|---|---|---|---|
| **Samples** | **O.D.** | **O.D. mean (x)** | **%CV** | **Log10 Concentration (y)** | **Concentration (ng/mL)** |
| DSP20203 | 1,936<br>1,900 | 1,918 | 1,334 | 0,824 | 6,666 |
| DSP20386 | 2,466<br>2,528 | 2,497 | 1,759 | | |
| DSP19970 | 1,564<br>1,183 | 1,374 | 19,611 | 1,549 | 35,389 |
| DSP19913 | 1,475<br>1,279 | 1,377 | 10,047 | 1,545 | 35,073 |
| DSP20286 | 1,577<br>1,236 | 1,406 | 17,141 | 1,508 | 32,225 |
| DSP19873 | 1,924<br>1,645 | 1,785 | 11,043 | 1,009 | 10,215 |
| DSP20330 | 2,052<br>1,701 | 1,876 | 13,214 | 0,882 | 7,619 |
| DSP20115 | 1,820<br>1,357 | 1,589 | 20,607 | 1,273 | 18,738 |
| DSP20333 | 1,582<br>1,771 | 1,677 | 7,952 | 1,156 | 14,328 |
| DSP20282 | 1,992<br>1,998 | 1,995 | 0,232 | 0,714 | 5,172 |
| DSP20047 | 1,578<br>1,690 | 1,634 | 4,838 | 1,213 | 16,329 |

| | | | | | |
|---|---|---|---|---|---|
| DSP20230 | 1,384<br>1,687 | 1,535 | 13,954 | 1,343 | 22,008 |
| DSP20078 | 1,291<br>1,618 | 1,455 | 15,904 | 1,447 | 27,966 |
| DSP20128 | 2,061<br>2,110 | 2,086 | 1,659 | 0,582 | 3,820 |
| DSP20195 | 2,004<br>1,787 | 1,895 | 8,091 | 0,855 | 7,168 |
| DSP20275 | 1,606<br>1,791 | 1,699 | 7,703 | 1,127 | 13,383 |
| DSP20021 | 1,565<br>1,565 | 1,565 | 0,006 | 1,304 | 20,160 |
| DSP20331 | 1,410<br>1,545 | 1,478 | 6,441 | 1,417 | 26,137 |
| DSP19895 | 1,814<br>1,850 | 1,832 | 1,408 | 0,944 | 8,782 |
| DSP20255 | 2,060<br>2,091 | 2,076 | 1,050 | 0,596 | 3,948 |
| DSP20008 | 1,206<br>1,173 | 1,190 | 1,958 | 1,774 | 59,422 |
| DSP20302 | 1,049<br>1,325 | 1,187 | 16,442 | 1,777 | 59,891 |
| DSP20299 | 1,186<br>0,813 | 1,000 | 26,440 | 1,996 | 99,118 |
| DSP20110 | 1,049<br>0,953 | 1,001 | 6,760 | 1,995 | 98,795 |
| DSP20373 | 1,361<br>1,287 | 1,324 | 3,939 | 1,610 | 40,764 |
| DSP20191 | 0,875<br>0,869 | 0,872 | 0,487 | 2,140 | 137,922 |
| DSP20037 | 1,254<br>1,167 | 1,210 | 5,056 | 1,749 | 56,160 |
| DSP20312 | 1,683<br>1,589 | 1,636 | 4,066 | 1,210 | 16,219 |
| DSP20367 | 1,182<br>1,092 | 1,137 | 5,595 | 1,837 | 68,730 |
| DSP20267 | 1,328<br>1,175 | 1,252 | 8,626 | 1,699 | 50,057 |

| **control** | | | | | |
|---|---|---|---|---|---|
| **Samples** | **O.D.** | **O.D. mean (x)** | **%CV** | **Log10 Concentration (y)** | **Concentration (ng/mL)** |
| nP19821 | 1,218<br>0,988 | 1,103 | 14,767 | 1,877 | 75,358 |
| nP20284 | 1,164<br>0,925 | 1,044 | 16,167 | 1,945 | 88,089 |
| nP20399 | 0,973<br>0,908 | 0,941 | 4,885 | 2,063 | 115,554 |
| nP19474 | 1,064<br>0,889 | 0,976 | 12,642 | 2,023 | 105,321 |
| nP19444 | 1,174<br>0,975 | 1,074 | 13,143 | 1,910 | 81,293 |

| Standard | | | | | |
|---|---|---|---|---|---|
| **Samples** | **O.D.** | **O.D. mean (x)** | **%CV** | **Concentration (ng/mL)** | **Log10 Concentration (y)** |
| STD1 | 0,421 | 0,364 | 22,272 | 200 | 2,301 |
| | 0,306 | | | | |
| STD2 | 0,539 | 0,643 | 22,861 | 100 | 2,000 |
| | 0,747 | | | | |
| STD3 | 1,002 | 0,862 | 23,056 | 50 | 1,699 |
| | 0,721 | | | | |
| STD4 | 1,077 | 1,020 | 7,809 | 25 | 1,398 |
| | 0,964 | | | | |
| STD5 | 1,090 | 1,146 | 6,909 | 12,5 | 1,097 |
| | 1,202 | | | | |
| STD6 | 1,707 | 1,459 | 24,010 | 6,25 | 0,796 |
| | 1,212 | | | | |
| STD7 | 1,894 | 1,853 | 3,160 | 3,12 | 0,494 |
| | 1,811 | | | | |
| STD8 | 2,220 | 2,207 | 0,887 | 0 | |
| | 2,193 | | | | |

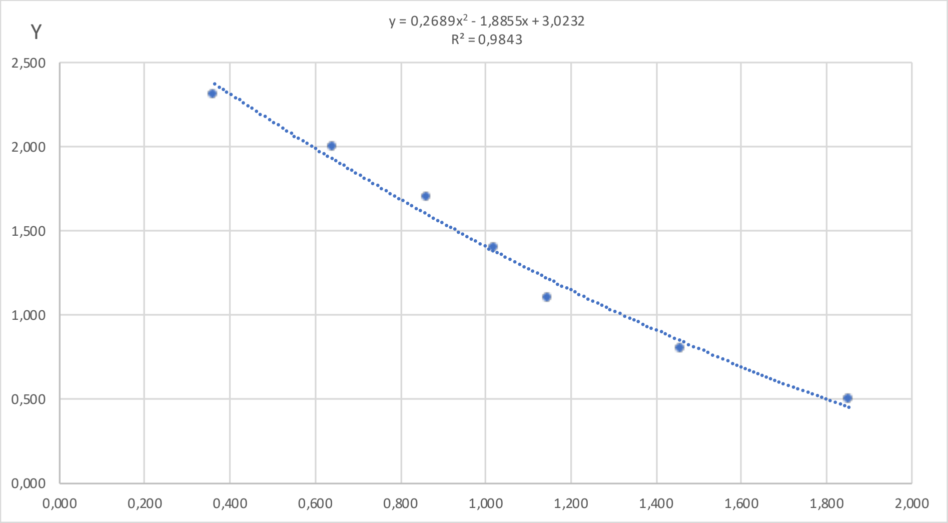


| y=a+bx+cx^2 | |
|---|---|
| a | 3,023 |
| b | -1,886 |
| c | 0,269 |
| R^2 | 0,984 |

| DS | | | | | |
|---|---|---|---|---|---|
| **Samples** | **O.D.** | **O.D. mean (x)** | **%CV** | **Log10 Concentration (y)** | **Concentration (ng/mL)** |
| DSP20091 | 1,085 | 1,0614 | 3,107 | 1,325 | 21,128 |
| | 1,038 | | | | |
| DSP20207 | 0,945 | 1,0476 | 13,890 | 1,343 | 22,031 |
| | 1,151 | | | | |
| DSP20200 | 1,332 | 1,0085 | 45,426 | 1,395 | 24,843 |
| | 0,685 | | | | |
| DSP20144 | 0,970 | 0,9515 | 2,769 | 1,473 | 29,691 |
| | 0,933 | | | | |
| DSP20374 | 0,713 | 0,7033 | 1,880 | 1,830 | 67,636 |
| | 0,694 | | | | |
| DSP20113 | 0,770 | 0,7896 | 3,453 | 1,702 | 50,357 |
| | 0,809 | | | | |
| DSP20360 | 0,853 | 0,7239 | 25,300 | 1,799 | 62,977 |
| | 0,594 | | | | |
| DSP20077 | 0,987 | 1,0594 | 9,719 | 1,328 | 21,260 |
| | 1,132 | | | | |
| DSP20096 | 1,314 | 1,2133 | 11,791 | 1,131 | 13,534 |
| | 1,112 | | | | |
| DSP20264 | 0,958 | 1,0173 | 8,192 | 1,383 | 24,172 |
| | 1,076 | | | | |
| DSP20189 | 0,993 | 1,0039 | 1,503 | 1,401 | 25,200 |
| | 1,015 | | | | |
| DSP19993 | 0,777 | 0,8032 | 4,572 | 1,682 | 48,118 |
| | 0,829 | | | | |
| DSP20381 | 1,036 | 0,9892 | 6,682 | 1,421 | 26,377 |
| | 0,942 | | | | |
| DSP20086 | 0,861 | 0,8297 | 5,401 | 1,644 | 44,054 |
| | 0,798 | | | | |
| DSP20080 | 1,115 | 0,9176 | 30,364 | 1,520 | 33,076 |
| | 0,721 | | | | |

| | | | | | |
|---|---|---|---|---|---|
| DSP20353 | 1,058<br>0,799 | 0,9286 | 19,749 | 1,504 | 31,932 |
| DSP19948 | 1,122<br>1,139 | 1,1305 | 1,052 | 1,235 | 17,192 |
| DSP20327 | 1,095<br>0,967 | 1,0311 | 8,780 | 1,365 | 23,172 |
| DSP20242 | 1,155<br>0,948 | 1,0511 | 13,929 | 1,338 | 21,802 |
| DSP20062 | 0,971<br>1,030 | 1,0006 | 4,160 | 1,406 | 25,456 |
| DSP20363 | 0,849<br>0,998 | 0,9234 | 11,465 | 1,511 | 32,461 |
| DSP20183 | 1,206<br>0,905 | 1,0555 | 20,133 | 1,333 | 21,509 |
| DSP19489 | 1,632<br>1,461 | 1,5463 | 7,789 | 0,751 | 5,631 |
| DSP20095 | 0,888<br>0,857 | 0,8722 | 2,522 | 1,583 | 38,303 |
| DSP19957 | 0,961<br>0,771 | 0,8660 | 15,489 | 1,592 | 39,090 |
| DSP20136 | 0,834<br>0,749 | 0,7913 | 7,647 | 1,700 | 50,071 |
| DSP20127 | 0,862<br>0,602 | 0,7321 | 25,123 | 1,787 | 61,230 |
| DSP20205 | 0,720<br>0,761 | 0,7404 | 3,916 | 1,775 | 59,514 |
| DSP20380 | 0,847<br>0,668 | 0,7576 | 16,779 | 1,749 | 56,119 |
| DSP20139 | 0,850<br>0,821 | 0,8352 | 2,430 | 1,636 | 43,246 |
| DSP20178 | 1,111<br>0,859 | 0,9847 | 18,106 | 1,427 | 26,746 |

| **control** | | | | | |
|---|---|---|---|---|---|
| **Samples** | **O.D.** | **O.D. mean (x)** | **%CV** | **Log10 Concentration (y)** | **Concentration (ng/mL)** |
| nP19968 | 0,555<br>0,718 | 0,637 | 18,067 | 1,932 | 85,425 |
| nP19461 | 0,748<br>0,739 | 0,743 | 0,897 | 1,770 | 58,910 |
| nP19653 | 0,942<br>0,856 | 0,899 | 6,789 | 1,546 | 35,138 |
| nP19454 | 0,607<br>0,644 | 0,626 | 4,202 | 1,949 | 88,833 |
| nP19467 | 0,801<br>0,943 | 0,872 | 11,501 | 1,584 | 38,347 |
| nP19487 | 0,862<br>0,967 | 0,914 | 8,133 | 1,524 | 33,419 |

| Standard | | | | | |
|---|---|---|---|---|---|
| **Samples** | **O.D.** | **O.D. mean (x)** | **%CV** | **Concentration (ng/mL)** | **Log10 Concentration (y)** |
| STD1 | 0,401<br>0,361 | 0,381 | 7,475 | 200 | 2,301 |
| STD2 | 0,659<br>0,779 | 0,719 | 11,800 | 100 | 2,000 |
| STD3 | 0,879<br>0,835 | 0,857 | 3,695 | 50 | 1,699 |
| STD4 | 0,985<br>0,924 | 0,954 | 4,491 | 25 | 1,398 |
| STD5 | 1,209<br>1,220 | 1,214 | 0,622 | 12,5 | 1,097 |
| STD6 | 1,559<br>1,461 | 1,510 | 4,564 | 6,25 | 0,796 |
| STD7 | 1,582<br>1,585 | 1,584 | 0,114 | 3,12 | 0,494 |
| STD8 | 1,636<br>1,598 | 1,617 | 1,634 | 0 | |

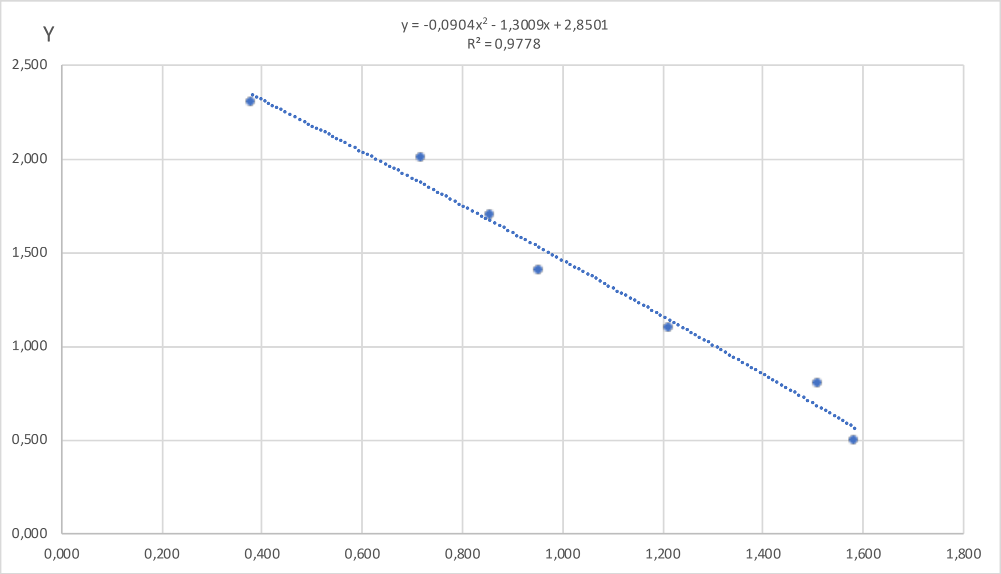


| y=a+bx+cx^2 | |
|---|---|
| a | 2,850 |
| b | -1,301 |
| c | -0,090 |
| R^2 | 0,978 |

| DS | | | | | |
|---|---|---|---|---|---|
| **Samples** | **O.D.** | **O.D. mean (x)** | **%CV** | **Log10 Concentration (y)** | **Concentration (ng/mL)** |
| DSP20359 | 1,896<br>1,757 | 1,827 | 5,385 | | |
| DSP20371 | 1,412<br>1,667 | 1,539 | 11,743 | 0,633 | 4,297 |
| DSP20376 | 0,799<br>0,797 | 0,798 | 0,200 | 1,755 | 56,851 |
| DSP20383 | 1,100<br>0,867 | 0,984 | 16,776 | 1,483 | 30,415 |
| DSP19916 | 0,707<br>0,715 | 0,711 | 0,857 | 1,879 | 75,757 |
| DSP19939 | 0,626<br>0,581 | 0,603 | 5,232 | 2,032 | 107,770 |
| DSP19952 | 0,891<br>0,707 | 0,799 | 16,338 | 1,753 | 56,646 |
| DSP19956 | 0,963<br>0,689 | 0,826 | 23,434 | 1,714 | 51,777 |
| DSP19989 | 1,916<br>1,765 | 1,840 | 5,832 | | |
| DSP20018 | 0,833<br>0,818 | 0,826 | 1,280 | 1,714 | 51,801 |
| DSP20034 | 0,758<br>0,516 | 0,637 | 26,810 | 1,984 | 96,489 |
| DSP20082 | 0,541<br>0,548 | 0,545 | 0,900 | 2,115 | 130,213 |
| DSP20092 | 0,499<br>0,613 | 0,556 | 14,485 | 2,099 | 125,646 |

| | | | | | |
|---|---|---|---|---|---|
| DSP20117 | 0,781<br>0,772 | 0,776 | 0,819 | 1,786 | 61,048 |
| DSP20121 | 0,800<br>0,688 | 0,744 | 10,570 | 1,832 | 67,964 |
| DSP20123 | 0,705<br>0,616 | 0,661 | 9,484 | 1,951 | 89,390 |
| DSP20140 | 0,861<br>0,893 | 0,877 | 2,573 | 1,640 | 43,671 |
| DSP20150 | 0,815<br>0,840 | 0,828 | 2,139 | 1,711 | 51,461 |
| DSP20155 | 0,914<br>0,743 | 0,828 | 14,581 | 1,711 | 51,364 |
| DSP20161 | 0,687<br>0,696 | 0,691 | 0,920 | 1,908 | 80,882 |
| DSP20164 | 0,809<br>0,675 | 0,742 | 12,770 | 1,835 | 68,395 |
| DSP20187 | 0,801<br>0,692 | 0,746 | 10,316 | 1,829 | 67,401 |
| DSP20194 | 0,796<br>0,569 | 0,682 | 23,546 | 1,921 | 83,292 |
| DSP20214 | 0,779<br>0,805 | 0,792 | 2,359 | 1,763 | 57,911 |
| DSP20220 | 0,844<br>0,700 | 0,772 | 13,226 | 1,792 | 61,941 |
| DSP20236 | 0,966<br>0,922 | 0,944 | 3,303 | 1,542 | 34,835 |
| DSP20257 | 1,075<br>0,788 | 0,932 | 21,796 | 1,560 | 36,268 |
| DSP20271 | 0,683<br>0,776 | 0,730 | 9,074 | 1,853 | 71,270 |
| DSP20274 | 1,040<br>1,123 | 1,082 | 5,464 | 1,337 | 21,747 |
| DSP20290 | 0,884<br>0,749 | 0,816 | 11,726 | 1,728 | 53,453 |
| DSP20294 | 0,816<br>0,511 | 0,663 | 32,548 | 1,948 | 88,622 |
| DSP20296 | 0,679<br>0,643 | 0,661 | 3,889 | 1,950 | 89,199 |
| DSP20368 | 1,183<br>1,034 | 1,109 | 9,566 | 1,297 | 19,814 |

| **control** | | | | | |
|---|---|---|---|---|---|
| **Samples** | **O.D.** | **O.D. mean (x)** | **%CV** | **Log10 Concentration (y)** | **Concentration (ng/mL)** |
| nP19460 | 0,737<br>0,688 | 0,713 | 4,842 | 1,877 | 75,324 |
| nP20231 | 0,716<br>0,595 | 0,655 | 13,071 | 1,959 | 91,015 |
| nP20045 | 0,690<br>0,661 | 0,675 | 3,065 | 1,930 | 85,198 |
| nP19946 | 0,605<br>0,614 | 0,610 | 1,101 | 2,023 | 105,553 |
| nP20273 | 0,683<br>0,663 | 0,673 | 2,119 | 1,933 | 85,733 |
| nP19458 | 0,621<br>0,599 | 0,610 | 2,560 | 2,023 | 105,413 |

| Standard | | | | | |
|---|---|---|---|---|---|
| **Samples** | **O.D.** | **O.D. mean (x)** | **%CV** | **Concentration (ng/mL)** | **Log10 Concentration (y)** |
| STD1 | 0,269 | 0,363 | 36,745 | 200 | 2,301 |
| | 0,457 | | | | |
| STD2 | 0,457 | 0,473 | 4,897 | 100 | 2,000 |
| | 0,489 | | | | |
| STD3 | 0,766 | 0,730 | 6,900 | 50 | 1,699 |
| | 0,695 | | | | |
| STD4 | 0,927 | 0,862 | 10,757 | 25 | 1,398 |
| | 0,796 | | | | |
| STD5 | 1,239 | 1,281 | 4,651 | 12,5 | 1,097 |
| | 1,324 | | | | |
| STD6 | 1,438 | 1,423 | 1,546 | 6,25 | 0,796 |
| | 1,407 | | | | |
| STD7 | 2,127 | 1,891 | 17,643 | 3,12 | 0,494 |
| | 1,655 | | | | |
| STD8 | 2,246 | 2,185 | 3,980 | 0 | |
| | 2,123 | | | | |

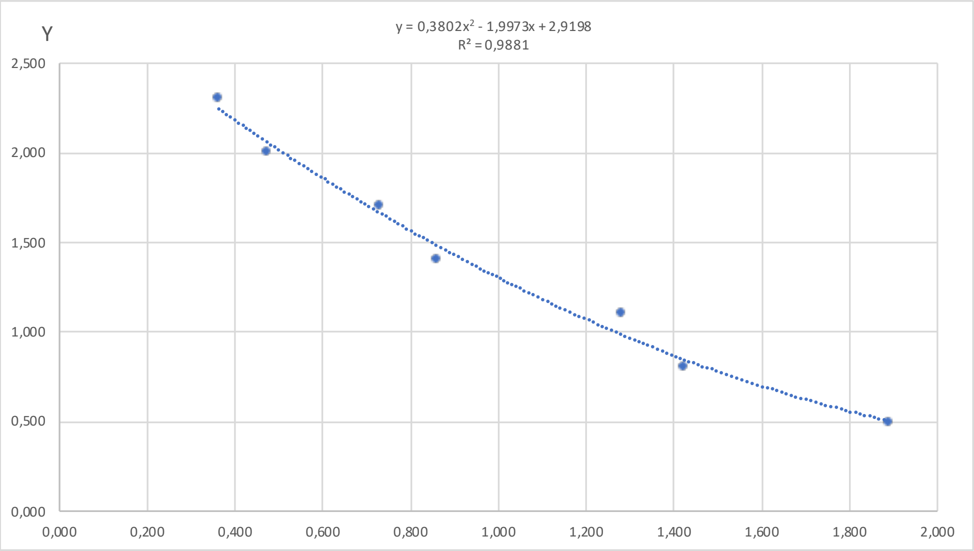


| y=a+bx+cx^2 | |
|---|---|
| a | 2,920 |
| b | -1,997 |
| c | 0,380 |
| R^2 | 0,988 |

| DS | | | | | |
|---|---|---|---|---|---|
| **Samples** | **O.D.** | **O.D. mean (x)** | **%CV** | **Log10 Concentration (y)** | **Concentration (ng/mL)** |
| DSP20379 | 1,341 | 1,347 | 0,660 | 0,919 | 8,307 |
| | 1,353 | | | | |
| DSP20387 | 2,612 | 2,620 | 0,425 | | |
| | 2,628 | | | | |
| DSP20388 | 1,018 | 0,979 | 5,559 | 1,329 | 21,316 |
| | 0,941 | | | | |
| DSP20085 | 1,227 | 1,227 | 0,073 | 1,042 | 11,014 |
| | 1,226 | | | | |
| DSP20103 | 1,138 | 1,014 | 17,310 | 1,285 | 19,296 |
| | 0,890 | | | | |
| DSP20104 | 1,643 | 1,432 | 20,823 | 0,839 | 6,908 |
| | 1,221 | | | | |
| DSP20112 | 0,897 | 0,861 | 5,980 | 1,482 | 30,352 |
| | 0,824 | | | | |
| DSP20132 | 1,282 | 1,186 | 11,477 | 1,086 | 12,193 |
| | 1,089 | | | | |
| DSP20235 | 1,915 | 1,748 | 13,495 | 0,590 | 3,894 |
| | 1,581 | | | | |
| DSP20279 | 2,437 | 2,483 | 2,629 | | |
| | 2,529 | | | | |
| DSP20350 | 0,914 | 0,864 | 8,144 | 1,478 | 30,047 |
| | 0,814 | | | | |
| DSP20366 | 1,334 | 1,072 | 34,582 | 1,216 | 16,447 |
| | 0,810 | | | | |

| control | | | | | |
|---|---|---|---|---|---|
| **Samples** | **O.D.** | **O.D. mean (x)** | **%CV** | **Log10 Concentration (y)** | **Concentration (ng/mL)** |

| | | | | | |
|---|---|---|---|---|---|
| nP19568 | 0,866<br>1,007 | 0,937 | 10,648 | 1,382 | 24,122 |
| nP19647 | 0,983<br>0,830 | 0,906 | 11,907 | 1,422 | 26,419 |
| nP19533 | 0,998<br>0,835 | 0,916 | 12,555 | 1,409 | 25,641 |
| nP19629 | 1,047<br>0,812 | 0,929 | 17,825 | 1,392 | 24,653 |
| nP19415 | 1,362<br>1,129 | 1,246 | 13,255 | 1,022 | 10,514 |

| Standard | | | | | |
|---|---|---|---|---|---|
| Samples | O.D. | O.D. mean (x) | %CV | Concentration (ng/mL) | Log10 Concentration (y) |
| STD1 | 1,194<br>1,170 | 1,182 | 1,442 | 200 | 2,301 |
| STD2 | 1,668<br>1,401 | 1,535 | 12,293 | 100 | 2,000 |
| STD3 | 2,169<br>1,922 | 2,045 | 8,545 | 50 | 1,699 |
| STD4 | 2,223<br>2,106 | 2,164 | 3,840 | 25 | 1,398 |
| STD5 | 2,214<br>2,360 | 2,287 | 4,526 | 12,5 | 1,097 |
| STD6 | 2,556<br>2,445 | 2,500 | 3,116 | 6,25 | 0,796 |
| STD7 | 2,409<br>2,616 | 2,513 | 5,836 | 3,12 | 0,494 |
| STD8 | 2,665<br>2,607 | 2,636 | 1,559 | 0 | |

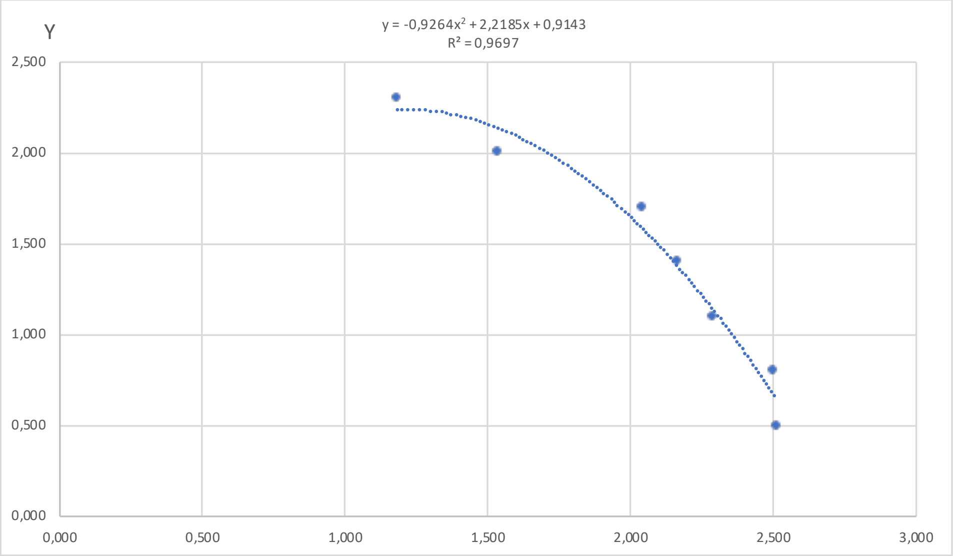


| y=a+bx+cx^2 | |
|---|---|
| a | 0,914 |
| b | 2,219 |
| c | -0,926 |
| R^2 | 0,970 |

| DS | | | | | |
|---|---|---|---|---|---|
| Samples | O.D. | O.D. mean (x) | %CV | Log10 Concentration (y) | Concentration (ng/mL) |
| DSP19559 | 2,715<br>2,552 | 2,634 | 4,370 | | |
| DSP19755 | 2,568<br>2,627 | 2,598 | 1,597 | | |
| DSP19757 | 1,624<br>2,140 | 1,882 | 19,398 | 1,808 | 64,294 |
| DSP19841 | 2,416<br>2,226 | 2,321 | 5,801 | 1,073 | 11,840 |
| DSP19897 | 1,993<br>1,840 | 1,917 | 5,662 | 1,763 | 57,962 |
| DSP19901 | 2,341<br>2,319 | 2,330 | 0,660 | 1,053 | 11,307 |
| DSP19958 | 2,594<br>2,470 | 2,532 | 3,465 | | |
| DSP20003 | 2,371<br>1,824 | 2,098 | 18,458 | 1,492 | 31,018 |
| DSP20030 | 2,143<br>1,902 | 2,023 | 8,422 | 1,612 | 40,902 |
| DSP20143 | 1,874<br>1,992 | 1,933 | 4,334 | 1,741 | 55,129 |
| DSP20156 | 2,499<br>2,538 | 2,518 | 1,081 | | |

| control | | | | | |
|---|---|---|---|---|---|
| Samples | O.D. | O.D. mean (x) | %CV | Log10 Concentration (y) | Concentration (ng/mL) |
| nP19494 | 1,991<br>1,987 | 1,989 | 0,142 | 1,662 | 45,871 |

| | | | | | |
|---|---|---|---|---|---|
| nP19606 | 2,319<br>2,201 | 2,260 | 3,709 | 1,197 | 15,732 |
| nP19631 | 1,978<br>1,931 | 1,954 | 1,685 | 1,712 | 51,471 |
| nP19660 | 1,982<br>2,026 | 2,004 | 1,561 | 1,640 | 43,610 |
| nP19691 | 2,250<br>2,278 | 2,264 | 0,883 | 1,188 | 15,433 |
| nP19718 | 2,387<br>2,386 | 2,386 | 0,005 | 0,933 | 8,562 |
| nP19727 | 1,959<br>2,108 | 2,034 | 5,176 | 1,594 | 39,286 |
| nP19737 | 1,931<br>2,138 | 2,035 | 7,174 | 1,593 | 39,186 |
| nP19783 | 2,049<br>1,834 | 1,942 | 7,836 | 1,729 | 53,634 |
| nP19812 | 2,218<br>1,974 | 2,096 | 8,242 | 1,494 | 31,198 |
| nP19822 | 1,908<br>1,845 | 1,877 | 2,353 | 1,815 | 65,340 |
| nP20228 | 1,947<br>1,723 | 1,835 | 8,656 | 1,866 | 73,459 |
| nP19854 | 2,215<br>2,164 | 2,190 | 1,655 | 1,330 | 21,402 |
| nP19856 | 2,240<br>2,400 | 2,320 | 4,851 | 1,075 | 11,887 |
| nP19902 | 2,082<br>2,049 | 2,065 | 1,145 | 1,545 | 35,037 |
| nP20246 | 1,713<br>1,780 | 1,746 | 2,698 | 1,963 | 91,917 |
| nP20341 | 1,621<br>1,701 | 1,661 | 3,408 | 2,043 | 110,515 |
| nP20343 | 1,748<br>1,648 | 1,698 | 4,157 | 2,010 | 102,354 |
| nP20398 | 2,023<br>1,929 | 1,976 | 3,338 | 1,681 | 47,949 |
