## Supplementary Table 2 for "One-carbon pathway metabolites are altered in the plasma of subjects with Down syndrome: relation to chromosomal dosage"

**Supplementary Table 2. Analysis of 5-methyl-THF ELISA assays.** Totally six 5-methyl-THF ELISA assays were performed and each assay is shown in a different Excel sheet named Plate 1, Plate 2, Plate 3, Plate 4, Plate 5, Plate 6. The table "Standard" shows the two absorbance (O.D.) detected for each standard sample. In the "O.D. mean (x)" column, the average O.D. is reported. The "%CV" column represent the Coefficient of Variability between the two O.D. measurment expressed as percentage, while the last column shows the known concentration of each standard sample supplied by ELISA kit. In order to build the standard curve, we plotted "O.D. mean (x)" of standards in the x-axis and "Concentration (ng/mL) (y)" of standards in the y-axis. The polynomial equation (y=a+bx+cx^2) reported in the graph was used to determine metabolites concentration of plasma samples ("Concentration (y)"), using interpolation of O.D. mean values reported in "O.D. mean (x)" column in tables "DS" and "control". The final concentration of plasma samples is obtained by the moltiplication of y value for the dilution factor. The results are reported in "Concentration (ng/mL)" column in tables "DS" and "control". *The Standard 8 of Plate 5 was not take into consideration in order to make the standard curve because its O.D. mean was lower than Standard 7.

| Standard | | | | |
|---|---|---|---|---|
| **Samples** | **O.D.** | **O.D. mean (x)** | **%CV** | **Concentration (ng/mL) (y)** |
| STD1 | 0,171<br>0,185 | 0,178 | 5,769 | 10 |
| STD2 | 0,300<br>0,291 | 0,296 | 2,250 | 5 |
| STD3 | 0,538<br>0,621 | 0,580 | 10,042 | 2,5 |
| STD4 | 0,913<br>0,780 | 0,846 | 11,152 | 1,25 |
| STD5 | 0,976<br>0,996 | 0,986 | 1,439 | 0,625 |
| STD6 | 1,059<br>1,134 | 1,096 | 4,873 | 0,313 |
| STD7 | 1,149<br>1,251 | 1,200 | 6,020 | 0,156 |
| STD8 | 1,290<br>1,042 | 1,166 | 15,026 | 0 |

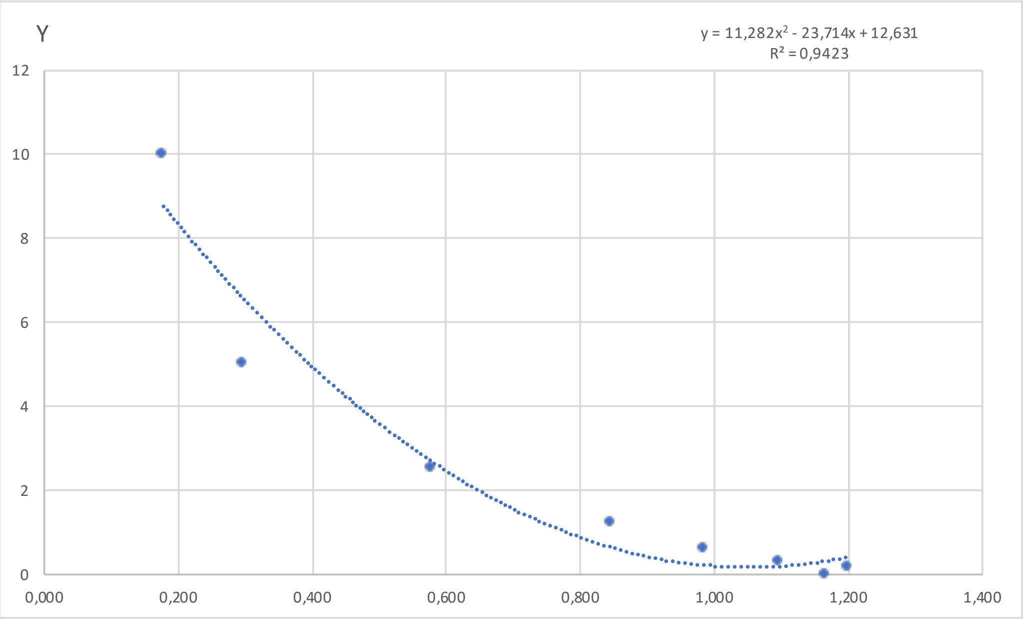


| y=a+bx+cx^2 | |
|---|---|
| a | 12,631 |
| b | -23,714 |
| c | 11,282 |
| R^2 | 0,942 |

| DS | | | | | |
|---|---|---|---|---|---|
| **Samples** | **O.D.** | **O.D. mean (x)** | **%CV** | **Concentration (y)** | **Concentration (ng/mL)** |
| DSP20203 | 0,482<br>0,470 | 0,476 | 1,806 | 3,904 | 39,042 |
| DSP20384 | 0,441<br>0,401 | 0,421 | 6,598 | 4,647 | 46,473 |
| DSP19970 | 0,330<br>0,342 | 0,336 | 2,655 | 5,936 | 59,361 |
| DSP19913 | 0,465<br>0,417 | 0,441 | 7,786 | 4,367 | 43,668 |

| | | | | | |
|---|---|---|---|---|---|
| DSP20286 | 0,277<br>0,293 | 0,285 | 3,794 | 6,788 | 67,879 |
| DSP19873 | 0,342<br>0,322 | 0,332 | 4,374 | 6,003 | 60,025 |
| DSP20380 | 0,416<br>0,373 | 0,394 | 7,584 | 5,034 | 50,337 |
| DSP20333 | 0,344<br>0,305 | 0,325 | 8,441 | 6,119 | 61,190 |
| DSP20282 | 0,449<br>0,413 | 0,431 | 5,951 | 4,510 | 45,096 |
| DSP20047 | 0,337<br>0,370 | 0,353 | 6,749 | 5,658 | 56,584 |
| DSP20230 | 0,368<br>0,402 | 0,385 | 6,145 | 5,177 | 51,766 |
| DSP20078 | 0,333<br>0,326 | 0,329 | 1,510 | 6,045 | 60,453 |
| DSP20275 | 0,416<br>0,420 | 0,418 | 0,718 | 4,685 | 46,850 |
| DSP20232 | 0,350<br>0,343 | 0,346 | 1,425 | 5,772 | 57,717 |
| DSP20331 | 0,292<br>0,290 | 0,291 | 0,622 | 6,684 | 66,836 |
| DSP19895 | 0,311<br>0,364 | 0,338 | 11,002 | 5,910 | 59,101 |
| DSP20008 | 0,321<br>0,367 | 0,344 | 9,369 | 5,806 | 58,060 |
| DSP20302 | 0,317<br>0,369 | 0,343 | 10,731 | 5,819 | 58,191 |
| DSP20299 | 0,340<br>0,353 | 0,347 | 2,809 | 5,767 | 57,668 |
| DSP20110 | 0,296<br>0,327 | 0,312 | 7,158 | 6,336 | 63,362 |
| DSP20373 | 0,328<br>0,349 | 0,339 | 4,418 | 5,893 | 58,934 |
| DSP20191 | 0,287<br>0,314 | 0,300 | 6,400 | 6,527 | 65,265 |
| DSP20267 | 0,314<br>0,345 | 0,330 | 6,665 | 6,040 | 60,403 |

| **control** | | | | | |
|---|---|---|---|---|---|
| **Samples** | **O.D.** | **O.D. mean (x)** | **%CV** | **Concentration (y)** | **Concentration (ng/mL)** |
| nP19821 | 0,466<br>0,480 | 0,473 | 2,109 | 3,943 | 39,431 |
| nP20284 | 0,290<br>0,316 | 0,303 | 6,039 | 6,480 | 64,797 |
| nP20399 | 0,272<br>0,325 | 0,298 | 12,676 | 6,560 | 65,604 |
| nP19365 | 0,358<br>0,258 | 0,308 | 22,957 | 6,398 | 63,983 |

| Standard | | | | |
|---|---|---|---|---|
| Samples | O.D. | O.D. mean (x) | %CV | Concentration (ng/mL) (y) |
| STD1 | 0,164<br>0,135 | 0,150 | 13,427 | 10 |
| STD2 | 0,284<br>0,285 | 0,285 | 0,095 | 5 |
| STD3 | 0,443<br>0,437 | 0,440 | 0,935 | 2,5 |
| STD4 | 0,635<br>0,616 | 0,625 | 2,166 | 1,25 |
| STD5 | 0,761<br>0,754 | 0,757 | 0,702 | 0,625 |
| STD6 | 0,867<br>0,856 | 0,862 | 0,887 | 0,313 |
| STD7 | 0,997<br>1,002 | 0,999 | 0,359 | 0,156 |
| STD8 | 0,925<br>0,903 | 0,914 | 1,723 | 0 |

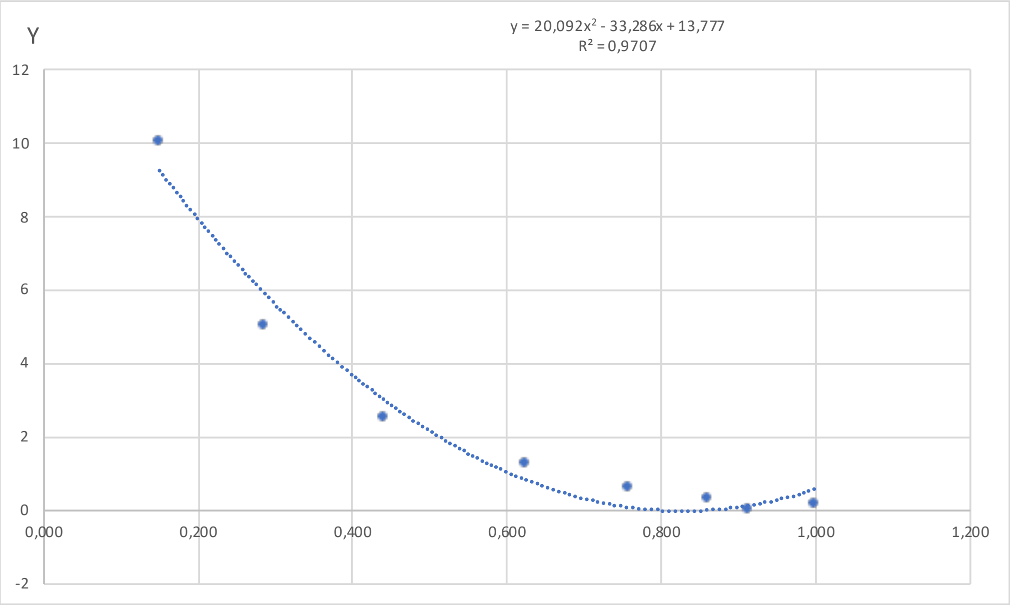


| y=a+bx+cx^2 | |
|---|---|
| a | 13,777 |
| b | -33,286 |
| c | 20,092 |
| R^2 | 0,9707 |

| DS | | | | | |
|---|---|---|---|---|---|
| Samples | O.D. | O.D. mean (x) | %CV | Concentration (ng/mL) (y) | Concentration (ng/mL) |
| DSP20091 | 0,409<br>0,400 | 0,404 | 1,561 | 3,602 | 36,020 |
| DSP20207 | 0,327<br>0,331 | 0,329 | 0,863 | 4,992 | 49,916 |
| DSP20200 | 0,332<br>0,360 | 0,346 | 5,722 | 4,673 | 46,730 |
| DSP20144 | 0,321<br>0,301 | 0,311 | 4,451 | 5,363 | 53,626 |
| DSP20374 | 0,303<br>0,307 | 0,305 | 0,924 | 5,497 | 54,966 |
| DSP20113 | 0,342<br>0,316 | 0,329 | 5,514 | 4,996 | 49,964 |
| DSP20107 | 0,305<br>0,360 | 0,332 | 11,548 | 4,933 | 49,327 |
| DSP20077 | 0,354<br>0,333 | 0,343 | 4,248 | 4,718 | 47,185 |
| DSP20096 | 0,338<br>0,385 | 0,362 | 9,125 | 4,364 | 43,639 |
| DSP20189 | 0,290<br>0,373 | 0,332 | 17,544 | 4,949 | 49,493 |
| DSP20265 | 0,303<br>0,345 | 0,324 | 9,283 | 5,099 | 50,993 |

| | | | | | |
|---|---|---|---|---|---|
| DSP20381 | 0,249<br>0,330 | 0,290 | 19,795 | 5,820 | 58,197 |
| DSP20086 | 0,281<br>0,314 | 0,298 | 7,769 | 5,647 | 56,468 |
| DSP20080 | 0,235<br>0,303 | 0,269 | 17,930 | 6,279 | 62,786 |
| DSP20353 | 0,319<br>0,277 | 0,298 | 9,956 | 5,651 | 56,506 |
| DSP19948 | 0,325<br>0,348 | 0,337 | 4,991 | 4,849 | 48,491 |
| DSP20327 | 0,246<br>0,331 | 0,288 | 20,740 | 5,850 | 58,497 |
| DSP20242 | 0,261<br>0,283 | 0,272 | 5,714 | 6,213 | 62,128 |
| DSP20062 | 0,257<br>0,260 | 0,259 | 0,770 | 6,508 | 65,085 |
| DSP20363 | 0,266<br>0,253 | 0,259 | 3,471 | 6,498 | 64,980 |
| DSP20183 | 0,293<br>0,299 | 0,296 | 1,519 | 5,690 | 56,897 |
| DSP19489 | 0,377<br>0,350 | 0,363 | 5,234 | 4,340 | 43,403 |
| DSP20095 | 0,247<br>0,279 | 0,263 | 8,787 | 6,413 | 64,129 |
| DSP19957 | 0,241<br>0,230 | 0,235 | 3,237 | 7,058 | 70,575 |
| DSP20136 | 0,218<br>0,265 | 0,241 | 13,950 | 6,912 | 69,121 |
| DSP20127 | 0,256<br>0,256 | 0,256 | 0,199 | 6,569 | 65,694 |
| DSP20205 | 0,274<br>0,299 | 0,286 | 6,109 | 5,894 | 58,937 |
| DSP20339 | 0,269<br>0,274 | 0,271 | 1,524 | 6,221 | 62,213 |
| DSP20139 | 0,245<br>0,270 | 0,258 | 6,740 | 6,537 | 65,368 |
| DSP20178 | 0,331<br>0,359 | 0,345 | 5,644 | 4,687 | 46,873 |

| **control** | | | | | |
|---|---|---|---|---|---|
| **Samples** | **O.D.** | **O.D. mean (x)** | **%CV** | **Concentration (ng/mL) (y)** | **Concentration (ng/mL)** |
| nP19968 | 0,253<br>0,282 | 0,267 | 7,685 | 6,320 | 63,204 |
| nP19461 | 0,259<br>0,293 | 0,276 | 8,832 | 6,117 | 61,173 |
| nP19653 | 0,300<br>0,352 | 0,326 | 11,231 | 5,066 | 50,663 |
| nP19454 | 0,209<br>0,243 | 0,226 | 10,402 | 7,279 | 72,788 |
| nP19467 | 0,237<br>0,270 | 0,254 | 9,291 | 6,626 | 66,260 |
| nP19487 | 0,288<br>0,317 | 0,302 | 6,794 | 5,555 | 55,548 |

| Standard | | | | |
|---|---|---|---|---|
| Samples | O.D. | O.D. mean (x) | %CV | Concentration (ng/mL) (y) |
| STD1 | 0,142<br>0,124 | 0,133 | 9,722 | 10 |
| STD2 | 0,228<br>0,241 | 0,234 | 3,984 | 5 |
| STD3 | 0,388<br>0,374 | 0,381 | 2,598 | 2,5 |
| STD4 | 0,585<br>0,598 | 0,591 | 1,454 | 1,25 |
| STD5 | 0,874<br>0,796 | 0,835 | 6,589 | 0,625 |
| STD6 | 0,964<br>0,817 | 0,890 | 11,676 | 0,313 |
| STD7 | 0,992<br>0,802 | 0,897 | 14,987 | 0,156 |
| STD8 | 0,915<br>0,913 | 0,914 | 0,145 | 0 |

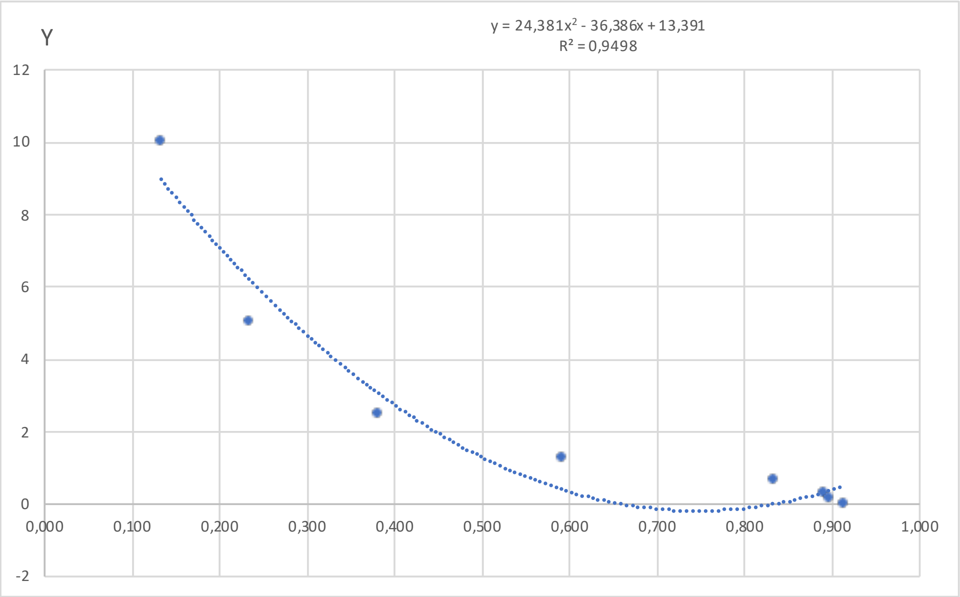


| y=a+bx+cx^2 | |
|---|---|
| a | 13,391 |
| b | -36,386 |
| c | 24,381 |
| R^2 | 0,950 |

| DS | | | | | |
|---|---|---|---|---|---|
| Samples | O.D. | O.D. mean (x) | %CV | Concentration (ng/mL) (y) | Concentration (ng/mL) |
| DSP20359 | 0,390<br>0,426 | 0,408 | 6,279 | 2,601 | 26,010 |
| DSP20371 | 0,347<br>0,261 | 0,304 | 20,157 | 4,586 | 45,858 |
| DSP20376 | 0,258<br>0,293 | 0,275 | 8,913 | 5,217 | 52,173 |
| DSP20383 | 0,264<br>0,309 | 0,287 | 11,003 | 4,962 | 49,616 |
| DSP19916 | 0,226<br>0,328 | 0,277 | 25,795 | 5,183 | 51,827 |
| DSP19939 | 0,309<br>0,330 | 0,320 | 4,708 | 4,250 | 42,503 |
| DSP19952 | 0,282<br>0,292 | 0,287 | 2,354 | 4,959 | 49,594 |
| DSP19956 | 0,315<br>0,308 | 0,312 | 1,576 | 4,415 | 44,147 |
| DSP19981 | 0,376<br>0,341 | 0,358 | 6,907 | 3,483 | 34,825 |
| DSP19989 | 0,238<br>0,269 | 0,253 | 8,536 | 5,740 | 57,399 |
| DSP20018 | 0,211<br>0,285 | 0,248 | 21,125 | 5,864 | 58,642 |

| | | | | | |
|---|---|---|---|---|---|
| DSP20034 | 0,219<br>0,288 | 0,253 | 19,259 | 5,735 | 57,351 |
| DSP20082 | 0,250<br>0,286 | 0,268 | 9,531 | 5,381 | 53,813 |
| DSP20092 | 0,214<br>0,258 | 0,236 | 13,333 | 6,167 | 61,665 |
| DSP20117 | 0,308<br>0,323 | 0,315 | 3,188 | 4,339 | 43,393 |
| DSP20121 | 0,296<br>0,307 | 0,301 | 2,453 | 4,641 | 46,410 |
| DSP20123 | 0,256<br>0,278 | 0,267 | 5,812 | 5,414 | 54,138 |
| DSP20150 | 0,285<br>0,240 | 0,263 | 12,291 | 5,519 | 55,188 |
| DSP20155 | 0,291<br>0,298 | 0,294 | 1,661 | 4,792 | 47,918 |
| DSP20161 | 0,285<br>0,269 | 0,277 | 3,967 | 5,177 | 51,766 |
| DSP20164 | 0,263<br>0,232 | 0,248 | 8,707 | 5,874 | 58,741 |
| DSP20187 | 0,287<br>0,245 | 0,266 | 11,134 | 5,444 | 54,444 |
| DSP20194 | 0,315<br>0,283 | 0,299 | 7,646 | 4,698 | 46,981 |
| DSP20214 | 0,246<br>0,334 | 0,290 | 21,412 | 4,881 | 48,808 |
| DSP20220 | 0,259<br>0,350 | 0,305 | 21,171 | 4,568 | 45,681 |
| DSP20236 | 0,304<br>0,319 | 0,312 | 3,258 | 4,419 | 44,194 |
| DSP20257 | 0,263<br>0,266 | 0,265 | 0,653 | 5,470 | 54,699 |
| DSP20271 | 0,286<br>0,298 | 0,292 | 2,773 | 4,844 | 48,442 |
| DSP20274 | 0,222<br>0,277 | 0,249 | 15,630 | 5,841 | 58,414 |
| DSP20290 | 0,242<br>0,272 | 0,257 | 8,388 | 5,648 | 56,476 |
| DSP20294 | 0,231<br>0,287 | 0,259 | 15,316 | 5,606 | 56,064 |
| DSP20296 | 0,321<br>0,344 | 0,333 | 4,802 | 3,985 | 39,847 |
| DSP20128 | 0,286<br>0,294 | 0,290 | 1,908 | 4,894 | 48,937 |

| control | | | | | |
|---|---|---|---|---|---|
| **Samples** | **O.D.** | **O.D. mean (x)** | **%CV** | **Concentration (ng/mL) (y)** | **Concentration (ng/mL)** |
| nP19460 | 0,277<br>0,361 | 0,319 | 18,733 | 4,261 | 42,614 |
| nP20231 | 0,287<br>0,333 | 0,310 | 10,378 | 4,449 | 44,494 |
| nP20045 | 0,261<br>0,313 | 0,287 | 12,864 | 4,955 | 49,554 |
| nP19946 | 0,329<br>0,304 | 0,316 | 5,590 | 4,324 | 43,243 |
| nP20273 | 0,325<br>0,332 | 0,329 | 1,606 | 4,064 | 40,641 |
| nP19458 | 0,349<br>0,371 | 0,360 | 4,358 | 3,457 | 34,567 |

| Standard | | | | |
|---|---|---|---|---|
| **Samples** | **O.D.** | **O.D. mean (x)** | **%CV** | **Concentration (ng/mL) (y)** |
| STD1 | 0,300<br>0,196 | 0,248 | 29,782 | 10 |
| STD2 | 0,490<br>0,419 | 0,455 | 11,109 | 5 |
| STD3 | 0,762<br>0,676 | 0,719 | 8,432 | 2,5 |
| STD4 | 1,231<br>1,170 | 1,200 | 3,566 | 1,25 |
| STD5 | 1,550<br>1,579 | 1,565 | 1,296 | 0,625 |
| STD6 | 1,783<br>1,649 | 1,716 | 5,500 | 0,313 |
| STD7 | 1,798<br>1,754 | 1,776 | 1,731 | 0,156 |
| STD8 | 1,726<br>1,760 | 1,743 | 1,357 | 0 |

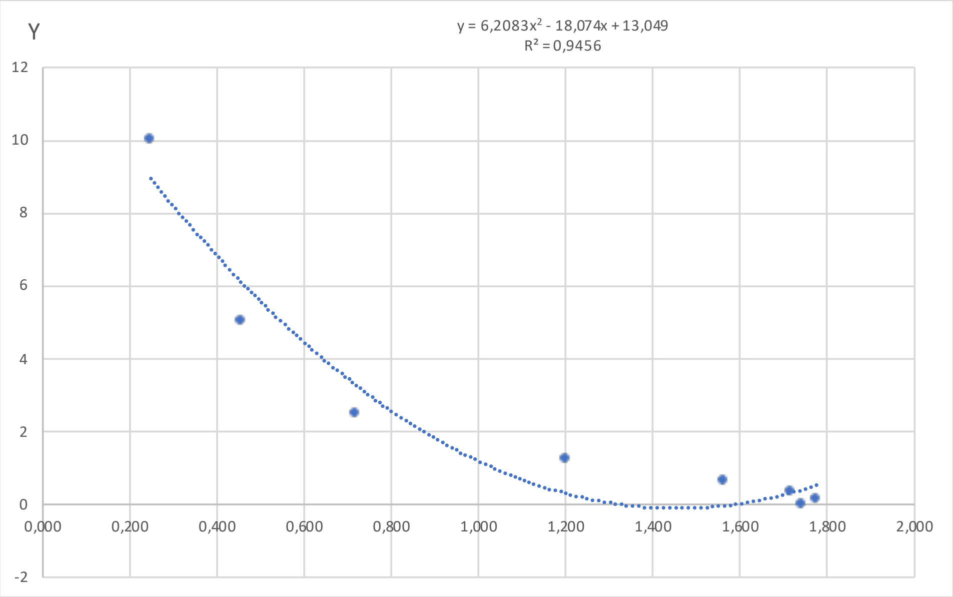


| y=a+bx+cx^2 | |
|---|---|
| a | 13,049 |
| b | -18,074 |
| c | 6,208 |
| R^2 | 0,946 |

| DS | | | | | |
|---|---|---|---|---|---|
| **Samples** | **O.D.** | **O.D. mean (x)** | **%CV** | **Concentration (ng/mL) (y)** | **Concentration (ng/mL)** |
| DSP20379 | 0,798<br>0,837 | 0,818 | 3,321 | 2,422 | 24,217 |
| DSP20388 | 0,641<br>0,744 | 0,692 | 10,477 | 3,512 | 35,123 |
| DSP20195 | 0,734<br>0,624 | 0,679 | 11,540 | 3,639 | 36,386 |
| DSP20255 | 0,687<br>0,625 | 0,656 | 6,633 | 3,866 | 38,655 |
| DSP20312 | 0,593<br>0,665 | 0,629 | 8,086 | 4,135 | 41,350 |
| DSP20264 | 0,669<br>0,651 | 0,660 | 1,931 | 3,825 | 38,255 |
| DSP20288 | 0,724<br>0,711 | 0,717 | 1,237 | 3,278 | 32,781 |
| DSP19596 | 0,635<br>0,554 | 0,595 | 9,601 | 4,494 | 44,944 |
| DSP20318 | 0,511<br>0,520 | 0,516 | 1,288 | 5,381 | 53,812 |
| DSP20104 | 0,500<br>0,505 | 0,502 | 0,641 | 5,536 | 55,358 |
| DSP20132 | 0,496<br>0,612 | 0,554 | 14,843 | 4,946 | 49,457 |
| DSP20235 | 0,685<br>0,667 | 0,676 | 1,929 | 3,667 | 36,672 |

| | | | | | |
|---|---|---|---|---|---|
| DSP20279 | 0,527<br>0,586 | 0,557 | 7,469 | 4,910 | 49,104 |
| DSP20350 | 0,511<br>0,433 | 0,472 | 11,662 | 5,904 | 59,036 |
| DSP20366 | 0,555<br>0,531 | 0,543 | 3,126 | 5,066 | 50,663 |

| control | | | | | |
|---|---|---|---|---|---|
| **Samples** | **O.D.** | **O.D. mean (x)** | **%CV** | **Concentration (ng/mL) (y)** | **Concentration (ng/mL)** |
| nP19568 | 0,656<br>0,600 | 0,628 | 6,332 | 4,150 | 41,499 |
| nP19647 | 0,669<br>0,662 | 0,666 | 0,743 | 3,766 | 37,662 |
| nP19737 | 0,528<br>0,606 | 0,567 | 9,751 | 4,797 | 47,971 |
| nP19533 | 0,665<br>0,766 | 0,716 | 9,966 | 3,294 | 32,941 |
| nP19629 | 0,733<br>0,836 | 0,784 | 9,291 | 2,691 | 26,911 |
| nP19415 | 0,694<br>0,744 | 0,719 | 4,916 | 3,262 | 32,621 |
| nP19474 | 0,662<br>0,628 | 0,645 | 3,669 | 3,975 | 39,754 |
| nP19576 | 0,785<br>0,765 | 0,775 | 1,799 | 2,769 | 27,694 |
| nP19418 | 0,720<br>0,691 | 0,706 | 2,878 | 3,387 | 33,870 |
| nP19416 | 0,617<br>0,611 | 0,614 | 0,763 | 4,291 | 42,912 |
| nP19567 | 0,561<br>0,511 | 0,536 | 6,592 | 5,148 | 51,477 |

| Standard | | | | |
|---|---|---|---|---|
| Samples | O.D. | O.D. mean (x) | %CV | Concentration (ng/mL) (y) |
| STD1 | 0,159<br>0,177 | 0,1684 | 7,619 | 10 |
| STD2 | 0,282<br>0,296 | 0,2892 | 3,367 | 5 |
| STD3 | 0,528<br>0,496 | 0,5123 | 4,445 | 2,5 |
| STD4 | 0,762<br>0,784 | 0,7728 | 2,053 | 1,25 |
| STD5 | 0,886<br>0,916 | 0,9011 | 2,306 | 0,625 |
| STD6 | 0,973<br>1,008 | 0,9904 | 2,504 | 0,313 |
| STD7 | 1,005<br>1,075 | 1,0403 | 4,785 | 0,156 |
| STD8* | 0,921<br>0,894 | 0,9075 | 2,132 | 0 |

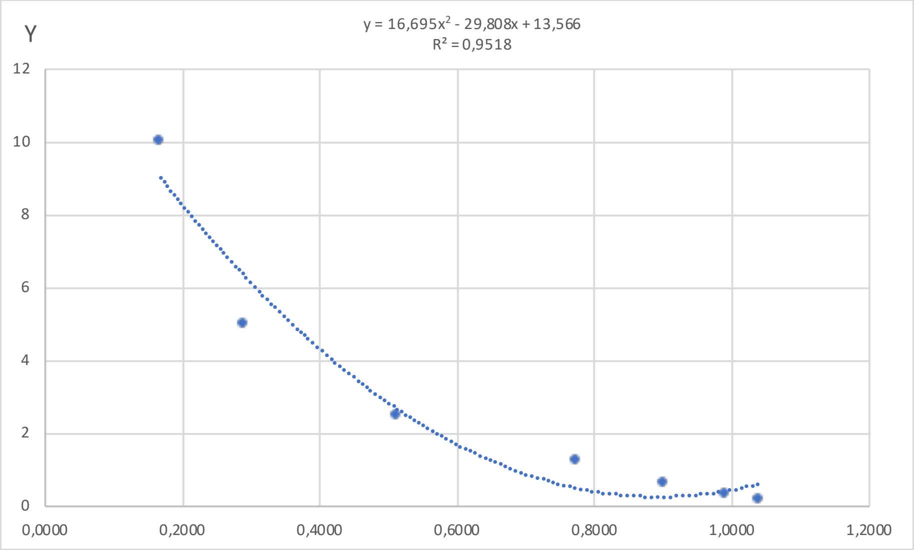


| y=a+bx+cx^2 | |
|---|---|
| a | 13,566 |
| b | -29,808 |
| c | 16,695 |
| R^2 | 0,952 |

| DS | | | | | |
|---|---|---|---|---|---|
| Samples | O.D. | O.D. mean (x) | %CV | Concentration (ng/mL) (y) | Concentration (ng/mL) |
| DSP19529 | 0,482<br>0,515 | 0,499 | 4,652 | 2,854 | 28,538 |
| DSP19759 | 0,452<br>0,455 | 0,453 | 0,430 | 3,487 | 34,871 |
| DSP19761 | 0,348<br>0,395 | 0,371 | 8,827 | 4,797 | 47,969 |
| DSP19770 | 0,364<br>0,411 | 0,387 | 8,605 | 4,524 | 45,243 |
| DSP19944 | 0,330<br>0,403 | 0,367 | 14,001 | 4,879 | 48,790 |
| DSP19966 | 0,363<br>0,372 | 0,368 | 1,698 | 4,861 | 48,611 |
| DSP20140 | 0,343<br>0,417 | 0,380 | 13,704 | 4,652 | 46,525 |
| DSP19997 | 0,443<br>0,430 | 0,437 | 2,032 | 3,734 | 37,345 |
| DSP20029 | 0,409<br>0,435 | 0,422 | 4,434 | 3,959 | 39,588 |
| DSP20068 | 0,418<br>0,417 | 0,418 | 0,084 | 4,031 | 40,311 |
| DSP20075 | 0,368<br>0,359 | 0,364 | 1,726 | 4,929 | 49,293 |
| DSP20098 | 0,378<br>0,375 | 0,376 | 0,608 | 4,710 | 47,102 |

| | | | | | |
|---|---|---|---|---|---|
| DSP20124 | 0,323<br>0,350 | 0,336 | 5,680 | 5,431 | 54,313 |
| DSP20126 | 0,424<br>0,446 | 0,435 | 3,553 | 3,752 | 37,524 |
| DSP20146 | 0,415<br>0,381 | 0,398 | 5,977 | 4,351 | 43,510 |
| DSP20157 | 0,506<br>0,471 | 0,488 | 5,044 | 2,993 | 29,930 |
| DSP20158 | 0,395<br>0,377 | 0,386 | 3,327 | 4,552 | 45,517 |
| DSP20162 | 0,399<br>0,342 | 0,371 | 10,857 | 4,808 | 48,083 |
| DSP20168 | 0,356<br>0,311 | 0,333 | 9,395 | 5,483 | 54,832 |
| DSP20212 | 0,385<br>0,394 | 0,390 | 1,653 | 4,485 | 44,851 |
| DSP20216 | 0,402<br>0,366 | 0,384 | 6,672 | 4,582 | 45,818 |
| DSP20224 | 0,404<br>0,390 | 0,397 | 2,435 | 4,367 | 43,671 |
| DSP20237 | 0,360<br>0,380 | 0,370 | 3,738 | 4,827 | 48,270 |
| DSP20240 | 0,428<br>0,428 | 0,428 | 0,056 | 3,868 | 38,685 |
| DSP20316 | 0,354<br>0,345 | 0,350 | 1,754 | 5,186 | 51,862 |
| DSP20329 | 0,322<br>0,294 | 0,308 | 6,404 | 5,974 | 59,744 |
| DSP20337 | 0,332<br>0,370 | 0,351 | 7,663 | 5,165 | 51,655 |
| DSP20347 | 0,345<br>0,353 | 0,349 | 1,469 | 5,195 | 51,950 |
| DSP20351 | 0,350<br>0,338 | 0,344 | 2,597 | 5,289 | 52,887 |
| DSP20361 | 0,340<br>0,410 | 0,375 | 13,246 | 4,743 | 47,426 |
| DSP20365 | 0,354<br>0,287 | 0,320 | 14,783 | 5,732 | 57,320 |

| **control** | | | | | |
|---|---|---|---|---|---|
| **Samples** | **O.D.** | **O.D. mean (x)** | **%CV** | **Concentration (ng/mL) (y)** | **Concentration (ng/mL)** |
| nP19783 | 0,462<br>0,492 | 0,477 | 4,430 | 3,146 | 31,458 |
| nP19645 | 0,459<br>0,452 | 0,455 | 1,123 | 3,455 | 34,550 |
| nP19419 | 0,393<br>0,411 | 0,402 | 3,140 | 4,287 | 42,874 |
| nP19417 | 0,353<br>0,429 | 0,391 | 13,752 | 4,461 | 44,614 |
| nP20398 | 0,369<br>0,438 | 0,404 | 12,083 | 4,253 | 42,534 |
| nP19718 | 0,448<br>0,483 | 0,465 | 5,336 | 3,314 | 33,138 |
| nP19856 | 0,422<br>0,423 | 0,423 | 0,210 | 3,950 | 39,500 |

| Standard | | | | |
|---|---|---|---|---|
| **Samples** | **O.D.** | **O.D. mean (x)** | **%CV** | **Concentration (ng/mL) (y)** |
| STD1 | 0,160<br>0,184 | 0,1723 | 9,848 | 10 |
| STD2 | 0,324<br>0,207 | 0,2656 | 31,147 | 5 |
| STD3 | 0,506<br>0,429 | 0,4672 | 11,704 | 2,5 |
| STD4 | 0,741<br>0,759 | 0,7500 | 1,642 | 1,25 |
| STD5 | 0,987<br>0,827 | 0,9072 | 12,483 | 0,625 |
| STD6 | 1,088<br>1,041 | 1,0643 | 3,115 | 0,313 |
| STD7 | 1,228<br>1,133 | 1,1808 | 5,675 | 0,156 |
| STD8 | 1,071<br>0,963 | 1,0171 | 7,510 | 0 |

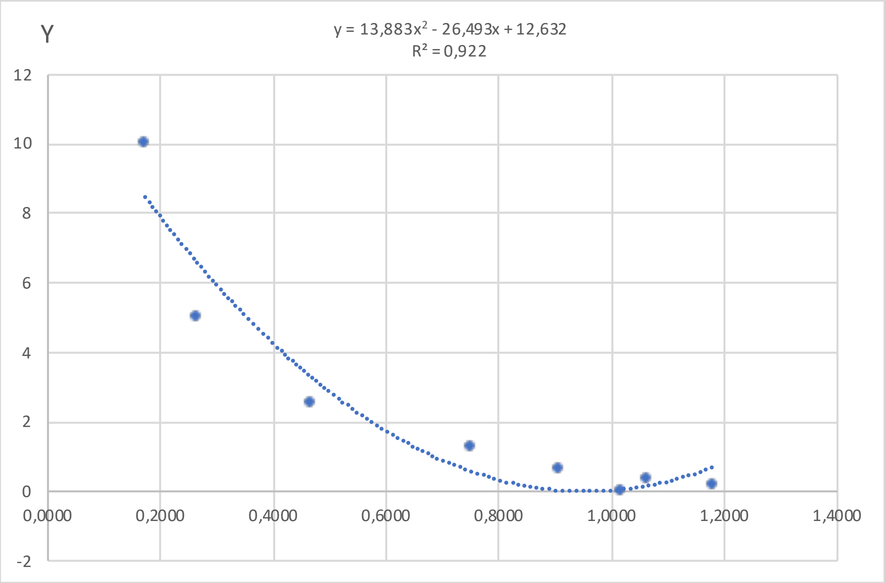


| y=a+bx+cx^2 | |
|---|---|
| a | 12,632 |
| b | -26,493 |
| c | 13,883 |
| R^2 | 0,922 |

| DS | | | | | |
|---|---|---|---|---|---|
| **Samples** | **O.D.** | **O.D. mean (x)** | **%CV** | **Concentration (ng/mL) (y)** | **Concentration (ng/mL)** |
| DSP19689 | 0,508<br>0,501 | 0,5043 | 0,986 | 2,802 | 28,021 |
| DSP19776 | 0,336<br>0,348 | 0,3420 | 2,473 | 5,196 | 51,957 |
| DSP19843 | 0,348<br>0,321 | 0,3347 | 5,725 | 5,320 | 53,202 |
| DSP19891 | 0,416<br>0,392 | 0,4038 | 4,221 | 4,198 | 41,981 |
| DSP19971 | 0,456<br>0,458 | 0,4570 | 0,251 | 3,425 | 34,246 |
| DSP20013 | 0,402<br>0,362 | 0,3816 | 7,391 | 4,544 | 45,443 |
| DSP20066 | 0,292<br>0,337 | 0,3141 | 10,113 | 5,680 | 56,797 |
| DSP20209 | 0,350<br>0,399 | 0,3743 | 9,216 | 4,661 | 46,614 |
