## Supplementary Table 3 for "One-carbon pathway metabolites are altered in the plasma of subjects with Down syndrome: relation to chromosomal dosage"

**Supplementary Table 3. Analysis of 5-formyl-THF ELISA assays**. Totally six 5-formyl-THF ELISA assays were performed and each assay is shown in a different Excel sheet named Plate 1, Plate 2, Plate 3, Plate 4, Plate 5, Plate 6. The table "Standard" shows the two absorbance (O.D.) detected for each standard sample. In the "O.D. mean" column, the average O.D. is reported. The "%CV" column represent the Coefficient of Variability between the two O.D. measurment expressed as percentage. The "Concentration (pg/mL)" column report the known concentration of each standard sample supplied by ELISA kit, while the last column shows the transformation of standard concentration values in their logarithm. In order to build the standard curve, we plotted "O.D. mean (x)" of standards in the x-axis and "Log 10 Concentration (y)" of standards in the y-axis. The polynomial equation (y=a+bx+cx^2) reported in the graph was used to determine metabolites concentration of plasma samples ("Log 10 Concentration (y)"), using interpolation of O.D. mean values reported in "O.D. mean (x)" column in tables "DS" and "control". The final concentration of plasma samples is obtained by conversion of logarithmic function. The results are reported in "Concentration (pg/mL)" column in tables "DS" and "control". In tables "DS" and "control" the O.D. mean values which are higher or lower than the O.D. mean values of standard sample ranges are reported in red and these values were not considered for the statistical analysis.

| Standard | | | | | |
|---|---|---|---|---|---|
| **Samples** | **O.D.** | **O.D. mean (x)** | **%CV** | **Concentration (pg/mL)** | **Log10 Concentration (y)** |
| STD1 | 0,053<br>0,045 | 0,049 | 11,382 | 150000 | 5,176 |
| STD2 | 0,068<br>0,064 | 0,066 | 4,899 | 30000 | 4,477 |
| STD3 | 0,130<br>0,137 | 0,134 | 3,408 | 7500 | 3,875 |
| STD4 | 0,380<br>0,225 | 0,303 | 36,290 | 1875 | 3,273 |
| STD5 | 0,607<br>0,512 | 0,559 | 12,067 | 468,8 | 2,671 |
| STD6 | 0,821<br>0,785 | 0,803 | 3,094 | 117,2 | 2,069 |
| STD7 | 1,097<br>1,108 | 1,103 | 0,736 | 0 | |

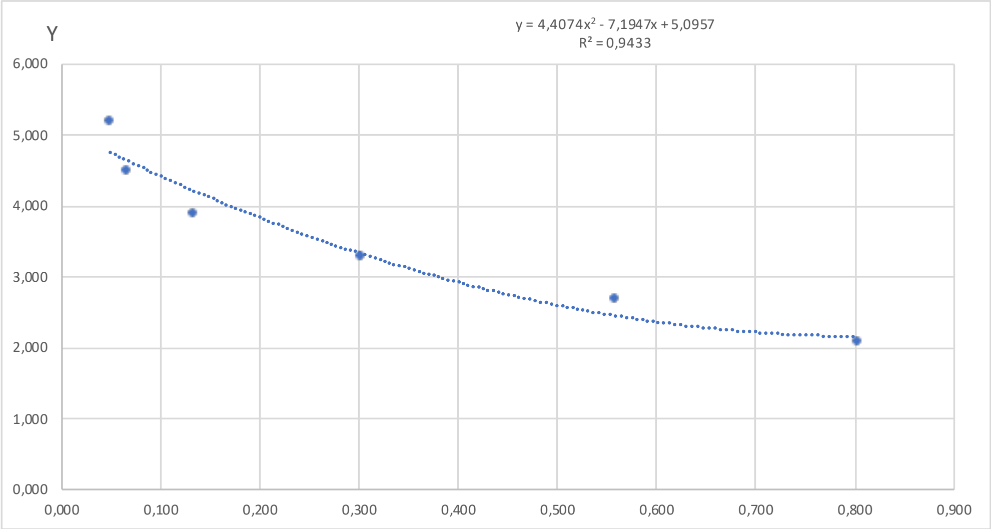


| y=a+bx+cx^2 | |
|---|---|
| a | 5,096 |
| b | -7,195 |
| c | 4,407 |
| R^2 | 0,943 |

| DS | | | | | |
|---|---|---|---|---|---|
| **Samples** | **O.D.** | **O.D. mean (x)** | **%CV** | **Log10 Concentration (y)** | **Concentration (pg/mL)** |
| DSP20384 | 0,753<br>0,744 | 0,748 | 0,852 | 2,180 | 151,279 |
| DSP19970 | 0,852<br>0,968 | 0,910 | 9,053 | | |
| DSP19913 | 0,791<br>0,870 | 0,831 | 6,780 | | |
| DSP20286 | 0,635<br>0,723 | 0,679 | 9,216 | 2,243 | 174,910 |
| DSP19873 | 0,549<br>0,653 | 0,601 | 12,195 | 2,363 | 230,856 |
| DSP20330 | 0,670<br>0,690 | 0,680 | 2,046 | 2,241 | 174,352 |
| DSP20115 | 0,702<br>0,700 | 0,701 | 0,175 | 2,218 | 165,173 |
| DSP20192 | 0,652<br>0,612 | 0,632 | 4,450 | 2,309 | 203,724 |

| | | | | | |
|---|---|---|---|---|---|
| DSP20333 | 0,943<br>0,777 | 0,860 | 13,641 | | |
| DSP20282 | 0,857<br>0,921 | 0,889 | 5,100 | | |
| DSP20047 | 0,819<br>0,856 | 0,838 | 3,145 | | |
| DSP20230 | 0,905<br>0,765 | 0,835 | 11,835 | | |
| DSP20078 | 0,645<br>0,807 | 0,726 | 15,773 | 2,196 | 156,928 |
| DSP20128 | 0,634<br>0,552 | 0,593 | 9,807 | 2,379 | 239,589 |
| DSP20195 | 0,646<br>0,667 | 0,656 | 2,267 | 2,272 | 187,206 |
| DSP20275 | 0,743<br>0,887 | 0,815 | 12,504 | | |
| DSP20232 | 0,777<br>0,932 | 0,855 | 12,794 | | |
| DSP20331 | 0,608<br>0,719 | 0,663 | 11,860 | 2,262 | 182,990 |
| DSP19895 | 0,545<br>0,830 | 0,687 | 29,255 | 2,233 | 170,896 |
| DSP20255 | 0,541<br>0,807 | 0,674 | 27,857 | 2,249 | 177,292 |
| DSP20302 | 0,720<br>0,781 | 0,751 | 5,758 | 2,178 | 150,812 |
| DSP20299 | 0,887<br>1,005 | 0,946 | 8,796 | | |
| DSP20110 | 0,775<br>0,749 | 0,762 | 2,414 | 2,172 | 148,760 |
| DSP20373 | 0,744<br>0,661 | 0,702 | 8,303 | 2,217 | 164,691 |
| DSP20191 | 0,706<br>0,669 | 0,688 | 3,857 | 2,232 | 170,757 |
| DSP20037 | 0,689<br>0,681 | 0,685 | 0,868 | 2,235 | 171,857 |
| DSP20312 | 0,585<br>0,567 | 0,576 | 2,301 | 2,414 | 259,457 |
| DSP20267 | 0,723<br>0,548 | 0,635 | 19,450 | 2,304 | 201,372 |

| control | | | | | |
|---|---|---|---|---|---|
| **Samples** | **O.D.** | **O.D. mean (x)** | **%CV** | **Log10 Concentration (y)** | **Concentration (pg/mL)** |
| nP19821 | 0,669<br>0,823 | 0,746 | 14,515 | 2,181 | 151,777 |
| nP20284 | 0,647<br>0,774 | 0,710 | 12,641 | 2,209 | 161,857 |
| nP20399 | 0,618<br>0,783 | 0,700 | 16,596 | 2,219 | 165,433 |
| nP19365 | 0,583<br>0,757 | 0,670 | 18,403 | 2,254 | 179,270 |
| nP19474 | 0,826<br>0,719 | 0,772 | 9,770 | 2,168 | 147,229 |
| nP19444 | 0,678<br>0,701 | 0,689 | 2,452 | 2,230 | 169,948 |

| Standard | | | | | |
|---|---|---|---|---|---|
| **Samples** | **O.D.** | **O.D. mean (x)** | **%CV** | **Concentration (pg/mL)** | **Log10 Concentration (y)** |
| STD1 | 0.049<br>0.049 | 0.049 | 0.069 | 150000 | 5.176 |
| STD2 | 0.057<br>0.058 | 0.058 | 1.463 | 30000 | 4.477 |
| STD3 | 0.084<br>0.080 | 0.082 | 3.664 | 7500 | 3.875 |
| STD4 | 0.185<br>0.180 | 0.182 | 1.689 | 1875 | 3.273 |
| STD5 | 0.360<br>0.405 | 0.383 | 8.427 | 468.8 | 2.671 |
| STD6 | 0.580<br>0.602 | 0.591 | 2.657 | 117.2 | 2.069 |
| STD7 | 0.818<br>0.797 | 0.808 | 1.810 | 0 | |

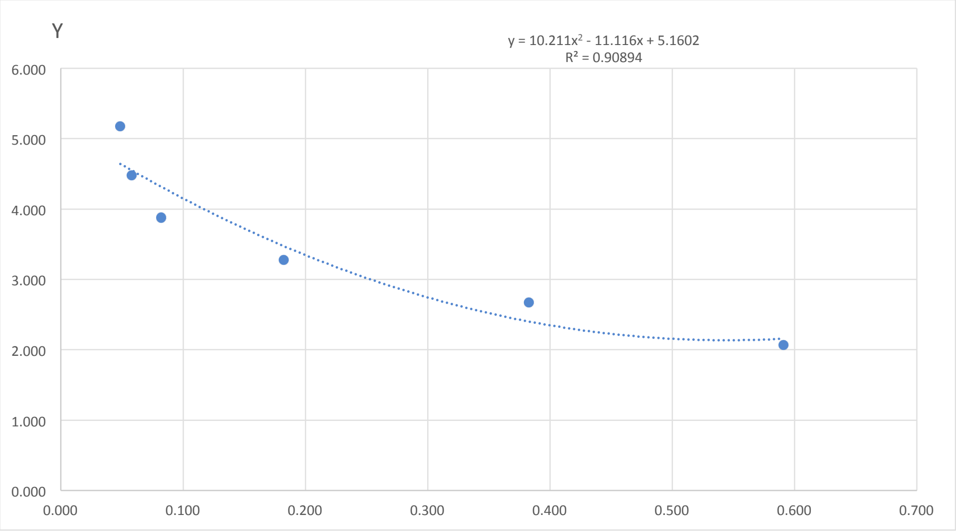


| y=a+bx+cx^2 | |
|---|---|
| a | 5.160 |
| b | -11.116 |
| c | 10.211 |
| R^2 | 0.909 |

| DS | | | | | |
|---|---|---|---|---|---|
| **Samples** | **O.D.** | **O.D. mean (x)** | **%CV** | **Log10 Concentration (y)** | **Concentration (pg/mL)** |
| DSP20091 | 0.571<br>0.546 | 0.558 | 3.142 | 2.137 | 137.060 |
| DSP20207 | 0.613<br>0.661 | 0.637 | 5.316 | | |
| DSP20200 | 0.602<br>0.699 | 0.651 | 10.541 | | |
| DSP20374 | 0.570<br>0.726 | 0.648 | 17.028 | | |
| DSP20113 | 0.698<br>0.577 | 0.637 | 13.421 | | |
| DSP20107 | 0.555<br>0.533 | 0.544 | 2.969 | 2.135 | 136.427 |
| DSP20077 | 0.484<br>0.593 | 0.539 | 14.313 | 2.135 | 136.529 |
| DSP20096 | 0.542<br>0.560 | 0.551 | 2.349 | 2.135 | 136.582 |
| DSP20264 | 0.765<br>0.770 | 0.768 | 0.439 | | |
| DSP20189 | 0.623<br>0.554 | 0.589 | 8.247 | 2.155 | 142.854 |
| DSP20265 | 0.665<br>0.668 | 0.666 | 0.317 | 2.287 | 193.446 |
| DSP20381 | 0.570<br>0.621 | 0.596 | 6.041 | | |
| DSP20086 | 0.594<br>0.538 | 0.566 | 6.957 | 2.140 | 137.955 |
| DSP20080 | 0.526<br>0.540 | 0.533 | 1.867 | 2.136 | 136.813 |
| DSP20353 | 0.594<br>0.672 | 0.633 | 8.734 | | |

| | | | | | |
|---|---|---|---|---|---|
| DSP19948 | 0.676 | 0.629 | 10.495 | | |
| | 0.582 | | | | |
| DSP20327 | 0.688 | 0.668 | 4.243 | | |
| | 0.648 | | | | |
| DSP20242 | 0.625 | 0.609 | 3.808 | | |
| | 0.592 | | | | |
| DSP20062 | 0.589 | 0.631 | 9.520 | | |
| | 0.674 | | | | |
| DSP20363 | 0.602 | 0.613 | 2.561 | | |
| | 0.624 | | | | |
| DSP20060 | 0.619 | 0.610 | 2.144 | | |
| | 0.601 | | | | |
| DSP19489 | 0.415 | 0.404 | 3.874 | 2.337 | 217.363 |
| | 0.393 | | | | |
| DSP19957 | 0.857 | 0.823 | 5.845 | | |
| | 0.789 | | | | |
| DSP20136 | 0.599 | 0.646 | 10.193 | | |
| | 0.692 | | | | |
| DSP20127 | 0.600 | 0.647 | 10.355 | | |
| | 0.695 | | | | |
| DSP20205 | 0.554 | 0.627 | 16.461 | | |
| | 0.700 | | | | |
| DSP20139 | 0.526 | 0.498 | 8.005 | 2.157 | 143.500 |
| | 0.470 | | | | |
| DSP20178 | 0.569 | 0.610 | 9.457 | | |
| | 0.651 | | | | |

| control | | | | | |
|---|---|---|---|---|---|
| **Samples** | **O.D.** | **O.D. mean (x)** | **%CV** | **Log10 Concentration (y)** | **Concentration (pg/mL)** |
| nP19968 | 0.692 | 0.734 | 8.018 | | |
| | 0.776 | | | | |
| nP19461 | 0.664 | 0.743 | 15.031 | | |
| | 0.822 | | | | |
| nP19653 | 0.477 | 0.606 | 30.049 | | |
| | 0.734 | | | | |
| nP19454 | 0.633 | 0.640 | 1.728 | | |
| | 0.648 | | | | |
| nP19467 | 0.382 | 0.470 | 26.467 | 2.191 | 155.354 |
| | 0.558 | | | | |
| nP19487 | 0.448 | 0.531 | 22.098 | 2.137 | 136.999 |
| | 0.614 | | | | |

| Standard | | | | | |
|---|---|---|---|---|---|
| Samples | O.D. | O.D. mean (x) | %CV | Concentration (pg/mL) | Log10 Concentration (y) |
| STD1 | 0.046<br>0.044 | 0.045 | 4.358 | 150000 | 5.176 |
| STD2 | 0.059<br>0.059 | 0.059 | 0.326 | 30000 | 4.477 |
| STD3 | 0.100<br>0.092 | 0.096 | 6.037 | 7500 | 3.875 |
| STD4 | 0.199<br>0.229 | 0.214 | 9.966 | 1875 | 3.273 |
| STD5 | 0.397<br>0.457 | 0.427 | 9.918 | 468.8 | 2.671 |
| STD6 | 0.561<br>0.760 | 0.661 | 21.251 | 117.2 | 2.069 |
| STD7 | 0.825<br>1.012 | 0.919 | 14.354 | 0 | |

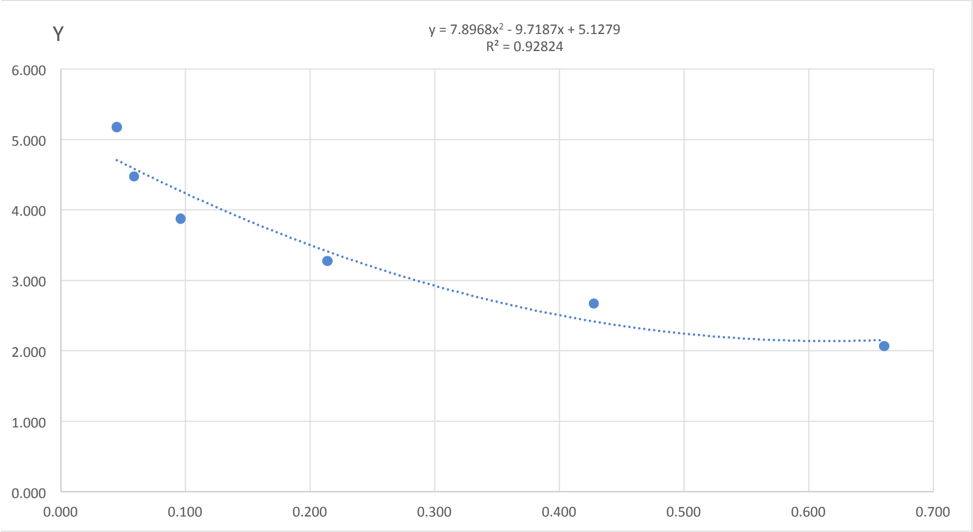


| y=a+bx+cx^2 | |
|---|---|
| a | 5.128 |
| b | -9.719 |
| c | 7.897 |
| R^2 | 0.928 |

| DS | | | | | |
|---|---|---|---|---|---|
| Samples | O.D. | O.D. mean (x) | %CV | Log10 Concentration (y) | Concentration (pg/mL) |
| DSP20359 | 0.323<br>0.345 | 0.334 | 4.579 | 2.763 | 579.917 |
| DSP20371 | 0.766<br>0.880 | 0.823 | 9.805 | | |
| DSP20376 | 0.704<br>0.816 | 0.760 | 10.421 | | |
| DSP20383 | 0.748<br>0.874 | 0.811 | 10.981 | | |
| DSP19826 | 0.648<br>0.776 | 0.712 | 12.637 | | |
| DSP19916 | 0.651<br>0.676 | 0.663 | 2.739 | | |
| DSP19939 | 0.611<br>0.621 | 0.616 | 1.171 | 2.138 | 137.301 |
| DSP19952 | 0.638<br>0.642 | 0.640 | 0.428 | 2.143 | 138.875 |
| DSP19956 | 0.514<br>0.485 | 0.500 | 4.134 | 2.243 | 175.164 |
| DSP19981 | 0.698<br>0.658 | 0.678 | 4.207 | | |
| DSP19989 | 0.589<br>0.509 | 0.549 | 10.339 | 2.172 | 148.754 |
| DSP20018 | 0.678<br>0.741 | 0.709 | 6.322 | | |
| DSP20034 | 0.583<br>0.687 | 0.635 | 11.578 | 2.141 | 138.228 |
| DSP20082 | 0.420<br>0.494 | 0.457 | 11.454 | 2.337 | 217.168 |
| DSP20092 | 0.539<br>0.572 | 0.556 | 4.125 | 2.166 | 146.494 |

| | | | | | |
|---|---|---|---|---|---|
| DSP20117 | 0.578<br>0.625 | 0.602 | 5.620 | 2.139 | 137.774 |
| DSP20121 | 0.595<br>0.587 | 0.591 | 0.902 | 2.142 | 138.767 |
| DSP20123 | 0.684<br>0.693 | 0.689 | 0.924 | | |
| DSP20140 | 0.562<br>0.700 | 0.631 | 15.437 | 2.140 | 137.905 |
| DSP20150 | 0.666<br>0.582 | 0.624 | 9.521 | 2.138 | 137.467 |
| DSP20155 | 0.620<br>0.737 | 0.678 | 12.235 | | |
| DSP20161 | 0.618<br>0.738 | 0.678 | 12.508 | | |
| DSP20164 | 0.594<br>0.654 | 0.624 | 6.770 | 2.138 | 137.490 |
| DSP20187 | 0.512<br>0.644 | 0.578 | 16.200 | 2.149 | 140.809 |
| DSP20194 | 0.533<br>0.601 | 0.567 | 8.509 | 2.156 | 143.278 |
| DSP20214 | 0.768<br>0.719 | 0.743 | 4.653 | | |
| DSP20220 | 0.676<br>0.623 | 0.650 | 5.739 | 2.147 | 140.262 |
| DSP20236 | 0.644<br>0.648 | 0.646 | 0.348 | 2.145 | 139.671 |
| DSP20257 | 0.617<br>0.608 | 0.613 | 1.047 | 2.138 | 137.314 |
| DSP20271 | 0.477<br>0.666 | 0.572 | 23.309 | 2.153 | 142.161 |
| DSP20274 | 0.480<br>0.489 | 0.484 | 1.283 | 2.273 | 187.636 |
| DSP20290 | 0.506<br>0.578 | 0.542 | 9.279 | 2.180 | 151.387 |
| DSP20294 | 0.571<br>0.465 | 0.518 | 14.547 | 2.213 | 163.255 |
| DSP20296 | 0.707<br>0.778 | 0.743 | 6.787 | | |

| control | | | | | |
|---|---|---|---|---|---|
| **Samples** | **O.D.** | **O.D. mean (x)** | **%CV** | **Log10 Concentration (y)** | **Concentration (pg/mL)** |
| nP19460 | 0.648<br>0.653 | 0.651 | 0.503 | 2.147 | 140.428 |
| nP20231 | 0.597<br>0.619 | 0.608 | 2.548 | 2.138 | 137.441 |
| nP20045 | 0.660<br>0.625 | 0.642 | 3.793 | 2.143 | 139.144 |
| nP19946 | 0.597<br>0.617 | 0.607 | 2.370 | 2.138 | 137.483 |
| nP20272 | 0.576<br>0.579 | 0.577 | 0.402 | 2.149 | 140.939 |
| nP19458 | 0.513<br>0.498 | 0.505 | 2.059 | 2.233 | 170.992 |

| Standard | | | | | |
|---|---|---|---|---|---|
| **Samples** | **O.D.** | **O.D. mean (x)** | **%CV** | **Concentration (pg/mL)** | **Log10 Concentration (y)** |
| STD1 | 0.047<br>0.047 | 0.047 | 0.084 | 150000 | 5.176 |
| STD2 | 0.064<br>0.070 | 0.067 | 6.298 | 30000 | 4.477 |
| STD3 | 0.111<br>0.124 | 0.118 | 7.789 | 7500 | 3.875 |
| STD4 | 0.270<br>0.269 | 0.270 | 0.238 | 1875 | 3.273 |
| STD5 | 0.644<br>0.696 | 0.670 | 5.480 | 468.8 | 2.671 |
| STD6 | 1.176<br>1.135 | 1.156 | 2.562 | 117.2 | 2.069 |
| STD7 | 1.545<br>1.462 | 1.503 | 3.910 | 0 | |

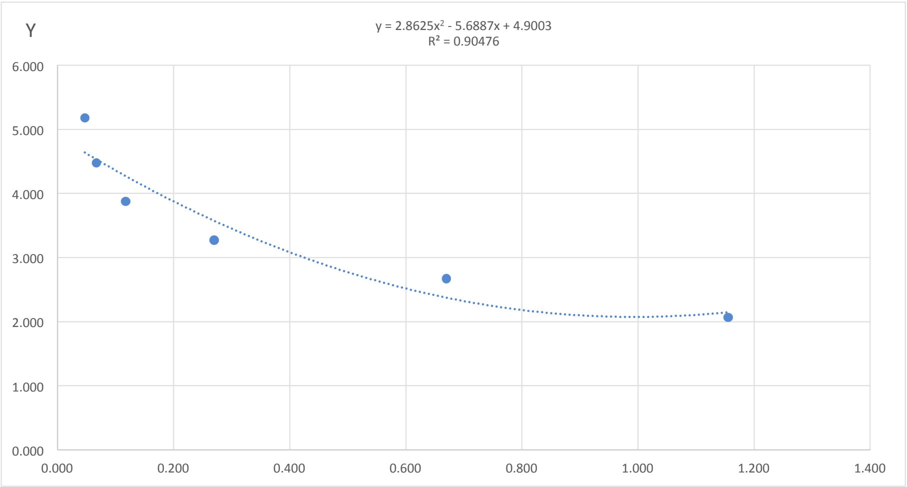


| y=a+bx+cx^2 | |
|---|---|
| a | 4.900 |
| b | -5.689 |
| c | 2.863 |
| R^2 | 0.905 |

| DS | | | | | |
|---|---|---|---|---|---|
| **Samples** | **O.D.** | **O.D. mean (x)** | **%CV** | **Log10 Concentration (y)** | **Concentration (pg/mL)** |
| DSP20379 | 1.164<br>0.898 | 1.031 | 18.218 | 2.078 | 119.681 |
| DSP20388 | 1.341<br>1.240 | 1.291 | 5.504 | | |
| DSP20001 | 1.044<br>1.186 | 1.115 | 9.021 | 2.116 | 130.635 |
| DSP20085 | 0.971<br>1.116 | 1.044 | 9.870 | 2.081 | 120.536 |
| DSP20103 | 1.266<br>1.280 | 1.273 | 0.767 | | |
| DSP20104 | 1.143<br>1.210 | 1.177 | 4.008 | | |
| DSP20112 | 1.295<br>1.085 | 1.190 | 12.484 | | |
| DSP20132 | 1.220<br>0.959 | 1.090 | 16.972 | 2.100 | 126.003 |
| DSP20235 | 1.197<br>1.128 | 1.162 | 4.169 | | |
| DSP20279 | 0.916<br>0.915 | 0.916 | 0.082 | 2.091 | 123.434 |
| DSP20350 | 1.047<br>1.159 | 1.103 | 7.198 | 2.108 | 128.335 |
| DSP20366 | 1.033<br>1.055 | 1.044 | 1.486 | 2.081 | 120.570 |

| control | | | | | |
|---|---|---|---|---|---|
| **Samples** | **O.D.** | **O.D. mean (x)** | **%CV** | **Log10 Concentration (y)** | **Concentration (pg/mL)** |
| nP19481 | 1.237<br>1.282 | 1.2596 | 2.555 | | |
| nP19367 | 1.174<br>1.145 | 1.1598 | 1.755 | | |
| nP19420 | 1.199<br>1.091 | 1.1453 | 6.665 | 2.140 | 137.984 |
| nP19469 | 1.109<br>1.037 | 1.0728 | 4.763 | 2.092 | 123.569 |

| Standard | | | | | |
|---|---|---|---|---|---|
| **Samples** | **O.D.** | **O.D. mean (x)** | **%CV** | **Concentration (pg/mL)** | **Log10 Concentration (y)** |
| STD1 | 0.055<br>0.054 | 0.054 | 1.569 | 150000 | 5.176 |
| STD2 | 0.065<br>0.074 | 0.070 | 9.952 | 30000 | 4.477 |
| STD3 | 0.114<br>0.110 | 0.112 | 2.370 | 7500 | 3.875 |
| STD4 | 0.242<br>0.227 | 0.234 | 4.406 | 1875 | 3.273 |
| STD5 | 0.484<br>0.456 | 0.470 | 4.242 | 468.8 | 2.671 |
| STD6 | 0.719<br>0.787 | 0.753 | 6.324 | 117.2 | 2.069 |
| STD7 | 1.096<br>1.071 | 1.084 | 1.610 | 0 | |

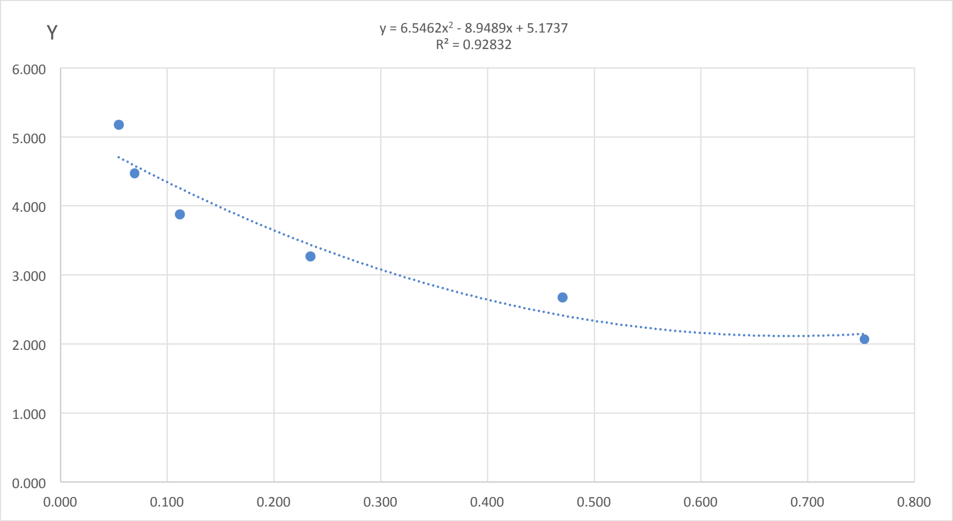


| y=a+bx+cx^2 | |
|---|---|
| a | 5.174 |
| b | -8.949 |
| c | 6.546 |
| R^2 | 0.928 |

| DS | | | | | |
|---|---|---|---|---|---|
| **Samples** | **O.D.** | **O.D. mean (x)** | **%CV** | **Log10 Concentration (y)** | **Concentration (pg/mL)** |
| DSP19529 | 0.693<br>0.608 | 0.650 | 9.277 | 2.122 | 132.577 |
| DSP19759 | 0.534<br>0.518 | 0.526 | 2.104 | 2.278 | 189.490 |
| DSP19761 | 0.850<br>0.807 | 0.829 | 3.670 | | |
| DSP19770 | 0.688<br>0.756 | 0.722 | 6.637 | 2.125 | 133.341 |
| DSP19944 | 0.738<br>0.843 | 0.790 | 9.420 | | |
| DSP19966 | 0.748<br>0.818 | 0.783 | 6.317 | | |
| DSP19997 | 0.754<br>0.703 | 0.729 | 4.947 | 2.129 | 134.494 |
| DSP20029 | 0.686<br>0.604 | 0.645 | 8.979 | 2.125 | 133.360 |
| DSP20068 | 0.834<br>0.802 | 0.818 | 2.741 | | |
| DSP20075 | 0.750<br>0.759 | 0.755 | 0.800 | | |
| DSP20098 | 0.751<br>0.788 | 0.770 | 3.319 | | |
| DSP20124 | 0.670<br>0.753 | 0.711 | 8.225 | 2.120 | 131.926 |
| DSP20126 | 0.756<br>0.796 | 0.776 | 3.679 | | |
| DSP20146 | 0.669<br>0.769 | 0.719 | 9.820 | 2.124 | 132.946 |
| DSP20157 | 0.694<br>0.647 | 0.671 | 4.892 | 2.116 | 130.740 |
| DSP20158 | 0.733<br>0.746 | 0.739 | 1.214 | 2.136 | 136.721 |

| | | | | | |
|---|---|---|---|---|---|
| DSP20162 | 0.730<br>0.710 | 0.720 | 1.916 | 2.124 | 133.053 |
| DSP20168 | 0.682<br>0.771 | 0.727 | 8.708 | 2.127 | 134.111 |
| DSP20212 | 0.658<br>0.780 | 0.719 | 11.934 | 2.124 | 132.929 |
| DSP20216 | 0.693<br>0.782 | 0.737 | 8.538 | 2.134 | 136.167 |
| DSP20224 | 0.680<br>0.981 | 0.831 | 25.673 | | |
| DSP20237 | 0.601<br>0.730 | 0.666 | 13.648 | 2.117 | 131.038 |
| DSP20240 | 0.759<br>0.651 | 0.705 | 10.854 | 2.118 | 131.337 |
| DSP20288 | 0.850<br>0.854 | 0.852 | 0.275 | | |
| DSP20316 | 0.771<br>0.715 | 0.743 | 5.314 | 2.138 | 137.485 |
| DSP20329 | 0.790<br>0.799 | 0.795 | 0.803 | | |
| DSP20337 | 0.680<br>0.717 | 0.698 | 3.752 | 2.117 | 130.851 |
| DSP20347 | 0.776<br>0.760 | 0.768 | 1.415 | | |
| DSP20351 | 0.700<br>0.678 | 0.689 | 2.290 | 2.116 | 130.470 |
| DSP20361 | 0.625<br>0.702 | 0.663 | 8.201 | 2.118 | 131.216 |
| DSP20365 | 0.633<br>0.635 | 0.634 | 0.174 | 2.131 | 135.292 |

| control | | | | | |
|---|---|---|---|---|---|
| **Samples** | **O.D.** | **O.D. mean (x)** | **%CV** | **Log10 Concentration (y)** | **Concentration (pg/mL)** |
| nP20398 | 0.864<br>0.915 | 0.889 | 4.099 | | |
| nP19856 | 0.724<br>0.719 | 0.721 | 0.492 | 2.125 | 133.261 |
| nP20343 | 0.809<br>0.949 | 0.879 | 11.301 | | |
| nP19606 | 0.560<br>0.677 | 0.619 | 13.444 | 2.143 | 138.989 |
| nP19854 | 0.703<br>0.739 | 0.721 | 3.516 | 2.124 | 133.184 |
| nP20228 | 0.715<br>0.686 | 0.701 | 2.917 | 2.117 | 130.994 |
| nP19494 | 0.717<br>0.707 | 0.712 | 0.956 | 2.121 | 131.988 |

| Standard | | | | | |
|---|---|---|---|---|---|
| **Samples** | **O.D.** | **O.D. mean (x)** | **%CV** | **Concentration (pg/mL)** | **Log10 Concentration (y)** |
| STD1 | 0.058<br>0.058 | 0.058 | 0.159 | 150000 | 5.176 |
| STD2 | 0.102<br>0.098 | 0.100 | 2.730 | 30000 | 4.477 |
| STD3 | 0.233<br>0.224 | 0.228 | 2.623 | 7500 | 3.875 |
| STD4 | 0.598<br>0.546 | 0.572 | 6.425 | 1875 | 3.273 |
| STD5 | 1.241<br>1.178 | 1.209 | 3.706 | 468.8 | 2.671 |
| STD6 | 2.067<br>2.027 | 2.047 | 1.377 | 117.2 | 2.069 |
| STD7 | 2.372<br>2.261 | 2.316 | 3.386 | 0 | |

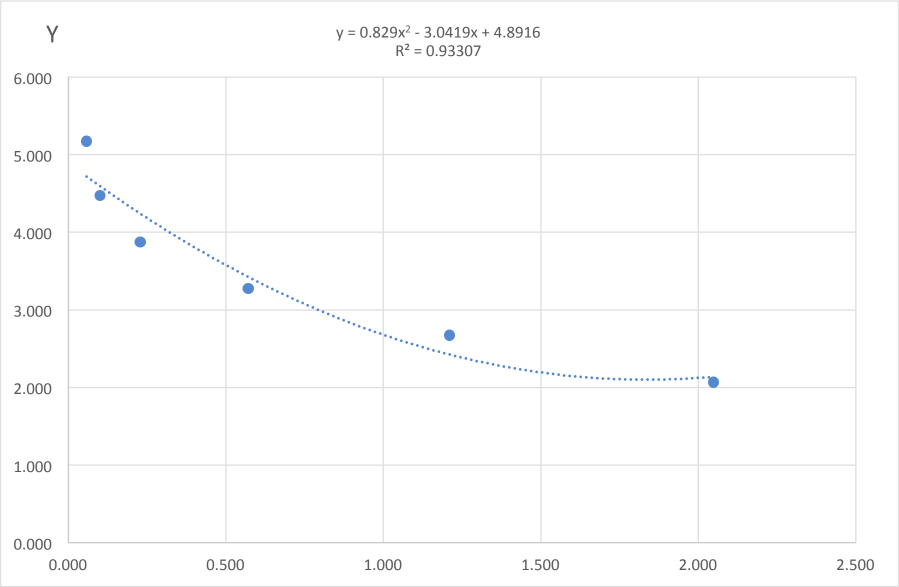


| y=a+bx+cx^2 | |
|---|---|
| a | 4.8916 |
| b | -3.0419 |
| c | 0.8290 |
| R^2 | 0.9331 |

| DS | | | | | |
|---|---|---|---|---|---|
| **Samples** | **O.D.** | **O.D. mean (x)** | **%CV** | **Log10 Concentration (y)** | **Concentration (pg/mL)** |
| DSP19971 | 1.847<br>1.863 | 1.855 | 0.589 | 2.101 | 126.324 |
| DSP20209 | 1.864<br>2.006 | 1.935 | 5.219 | 2.109 | 128.677 |
