## Supplementary Table 4 for "One-carbon pathway metabolites are altered in the plasma of subjects with Down syndrome: relation to chromosomal dosage"

**Supplementary Table 4. Analysis of SAH ELISA assays**. Totally six SAH ELISA assays were performed and each assay is shown in a different Excel sheet named Plate 1, Plate 2, Plate 3, Plate 4, Plate 5, Plate 6. The table "Standard" shows the two absorbance (O.D.) detected for each standard sample. In the "O.D. mean" column, the average O.D. is reported. The "%CV" column represent the Coefficient of Variability between the two O.D. measurment expressed as percentage. The "O.D. mean without background (x)" column report the values of O.D mean of the standard samples whithout the O.D. mean of the standard "STD8" which represent the absorbance of the diluent used to resuspend the standards and to dilute the plasma sample, while the last column shows the known concentration of each standard sample supplied by ELISA kit. In order to build the standard curve, we plotted "O.D. mean without background (x)" of standards in the x-axis and "Concentration (ng/mL) (y)" of standards in the y-axis. The polynomial equation (y=a+bx+cx^2) reported in the graph was used to determine metabolites concentration of plasma samples ("Concentration (y)"), using interpolation of O.D. mean values reported in "O.D. mean without background (x)" column in tables "DS" and "control". The final concentration of plasma samples is obtained by the moltiplication of y value for the dilution factor. The results are reported in "Concentration (ng/mL)" column in tables "DS" and "control". In tables "DS" and "control" the O.D. mean without background values which are higher or lower than the O.D. mean without background values of standard sample ranges are reported in red and these values were not considered for the statistical analysis.*The Standard 7 of Plate 3 was not take into consideration in order to make the standard curve because its O.D. mean was higher

| Standard | | | | | |
|---|---|---|---|---|---|
| **Samples** | **O.D.** | **O.D. mean** | **%CV** | **O.D. mean without background (x)** | **Concentration (ng/mL) (y)** |
| STD1 | 2.295 | 2.166 | 8.399 | 2.114 | 20 |
| | 2.037 | | | | |
| STD2 | 1.283 | 1.260 | 2.513 | 1.209 | 10 |
| | 1.238 | | | | |
| STD3 | 0.696 | 0.685 | 2.209 | 0.633 | 5 |
| | 0.674 | | | | |
| STD4 | 0.448 | 0.459 | 3.192 | 0.407 | 2.5 |
| | 0.469 | | | | |
| STD5 | 0.329 | 0.304 | 11.410 | 0.252 | 1.25 |
| | 0.280 | | | | |
| STD6 | 0.269 | 0.243 | 15.338 | 0.191 | 0.625 |
| | 0.217 | | | | |
| STD7 | 0.181 | 0.185 | 2.939 | 0.133 | 0.312 |
| | 0.188 | | | | |
| STD8 | 0.049 | 0.052 | 8.334 | | |
| | 0.055 | | | | |

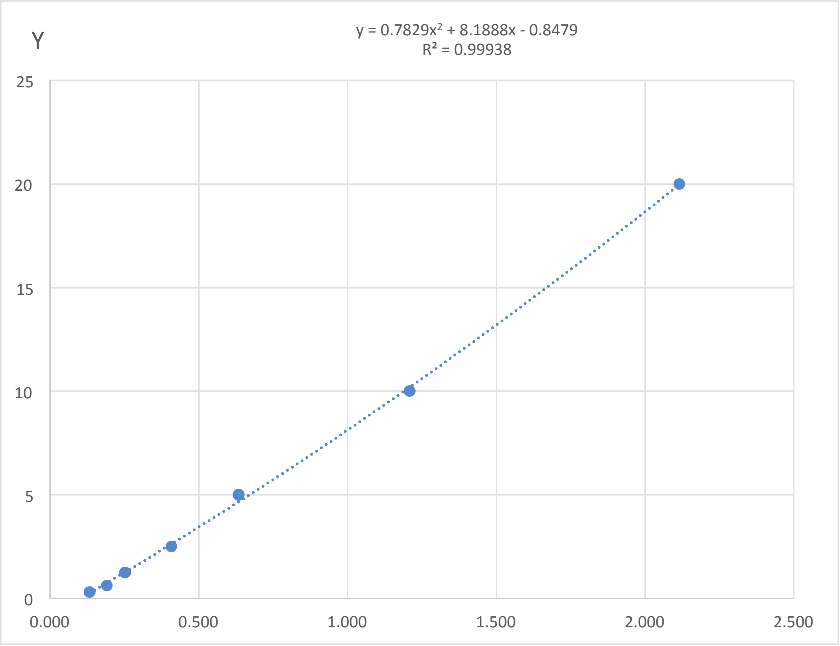


| y=a+bx+cx^2 | |
|---|---|
| a | -0.848 |
| b | 8.189 |
| c | 0.783 |
| R^2 | 0.999 |

| DS | | | | | | |
|---|---|---|---|---|---|---|
| Samples | O.D. | O.D. mean | %CV | O.D. mean without background (x) | Concentration (y) | Concentration (ng/mL) |
| DSP20203 | 0.260<br>0.275 | 0.267 | 4.114 | 0.216 | 0.954 | 4.768 |
| DSP20386 | 0.275<br>0.248 | 0.261 | 7.213 | 0.209 | 0.902 | 4.510 |
| DSP19970 | 0.241<br>0.233 | 0.237 | 2.358 | 0.185 | 0.698 | 3.488 |
| DSP19913 | 0.193<br>0.169 | 0.181 | 9.111 | 0.129 | | |
| DSP20286 | 0.163<br>0.185 | 0.174 | 9.258 | 0.122 | | |
| DSP19873 | 0.182<br>0.230 | 0.206 | 16.276 | 0.154 | 0.435 | 2.174 |
| DSP20330 | 0.200<br>0.392 | 0.296 | 45.931 | 0.244 | 1.197 | 5.984 |
| DSP20115 | 0.240<br>0.227 | 0.233 | 3.925 | 0.182 | 0.665 | 3.323 |
| DSP20192 | 0.372<br>0.233 | 0.303 | 32.363 | 0.251 | 1.255 | 6.275 |
| DSP20333 | 0.342<br>0.173 | 0.258 | 46.362 | 0.206 | 0.872 | 4.359 |
| DSP20282 | 0.273<br>0.177 | 0.225 | 29.990 | 0.173 | 0.593 | 2.965 |
| DSP20047 | 0.389<br>0.274 | 0.331 | 24.594 | 0.279 | 1.502 | 7.510 |
| DSP20230 | 0.327<br>0.279 | 0.303 | 11.126 | 0.251 | 1.260 | 6.298 |
| DSP20078 | 0.229<br>0.253 | 0.241 | 6.923 | 0.189 | 0.729 | 3.643 |
| DSP20128 | 0.336<br>0.260 | 0.298 | 17.950 | 0.246 | 1.215 | 6.076 |
| DSP20195 | 0.211<br>0.254 | 0.232 | 13.087 | 0.180 | 0.656 | 3.278 |
| DSP20089 | 0.284<br>0.193 | 0.239 | 26.920 | 0.187 | 0.712 | 3.561 |
| DSP20275 | 0.293<br>0.184 | 0.238 | 32.441 | 0.186 | 0.705 | 3.527 |
| DSP20021 | 0.224<br>0.225 | 0.225 | 0.151 | 0.173 | 0.591 | 2.957 |
| DSP20331 | 0.338<br>0.300 | 0.319 | 8.468 | 0.267 | 1.396 | 6.982 |
| DSP20174 | 0.313<br>0.195 | 0.254 | 32.997 | 0.202 | 0.839 | 4.197 |
| DSP19895 | 0.222<br>0.248 | 0.235 | 7.939 | 0.183 | 0.678 | 3.389 |
| DSP20255 | 0.219<br>0.202 | 0.211 | 5.515 | 0.159 | 0.472 | 2.358 |
| DSP20008 | 0.258<br>0.357 | 0.308 | 22.837 | 0.256 | 1.298 | 6.490 |
| DSP20302 | 0.343<br>0.350 | 0.347 | 1.319 | 0.295 | 1.634 | 8.168 |
| DSP20299 | 0.216<br>0.202 | 0.209 | 4.796 | 0.158 | 0.462 | 2.309 |
| DSP20110 | 0.315<br>0.239 | 0.277 | 19.555 | 0.225 | 1.036 | 5.182 |
| DSP20373 | 0.260<br>0.263 | 0.261 | 0.760 | 0.210 | 0.902 | 4.511 |
| DSP20191 | 0.272<br>0.209 | 0.240 | 18.674 | 0.189 | 0.725 | 3.623 |

| | | | | | | |
|---|---|---|---|---|---|---|
| DSP20037 | 0.339<br>0.230 | 0.284 | 27.049 | 0.232 | 1.098 | 5.488 |
| DSP20312 | 0.246<br>0.238 | 0.242 | 2.319 | 0.190 | 0.738 | 3.691 |
| DSP20367 | 0.209<br>0.157 | 0.183 | 20.035 | 0.131 | | |
| DSP20267 | 0.179<br>0.163 | 0.171 | 6.622 | 0.119 | | |

| control | | | | | | |
|---|---|---|---|---|---|---|
| **Samples** | **O.D.** | **O.D. mean** | **%CV** | **O.D. mean without background (x)** | **Concentration (y)** | **Concentration (ng/mL)** |
| nP19821 | 0.167<br>0.203 | 0.185 | 13.499 | 0.133 | 0.257 | 1.283 |
| nP20284 | 0.166<br>0.174 | 0.170 | 3.468 | 0.118 | | |
| nP20399 | 0.154<br>0.189 | 0.171 | 14.668 | 0.120 | | |
| nP19365 | 0.170<br>0.155 | 0.162 | 6.447 | 0.111 | | |
| nP19474 | 0.197<br>0.189 | 0.193 | 2.780 | 0.141 | 0.322 | 1.611 |
| nP19444 | 0.197<br>0.195 | 0.196 | 0.747 | 0.144 | 0.347 | 1.736 |

| Standard | | | | | |
|---|---|---|---|---|---|
| **Samples** | **O.D.** | **O.D. mean** | **%CV** | **O.D. mean without background (x)** | **Concentration (ng/mL) (y)** |
| STD1 | 1.171<br>1.347 | 1.259 | 9.913 | 1.204 | 20 |
| STD2 | 0.610<br>0.653 | 0.632 | 4.798 | 0.577 | 10 |
| STD3 | 0.339<br>0.385 | 0.362 | 9.007 | 0.307 | 5 |
| STD4 | 0.222<br>0.240 | 0.231 | 5.504 | 0.176 | 2.5 |
| STD5 | 0.163<br>0.184 | 0.173 | 8.767 | 0.118 | 1.25 |
| STD6 | 0.155<br>0.134 | 0.144 | 10.514 | 0.090 | 0.625 |
| STD7 | 0.116<br>0.170 | 0.143 | 26.738 | 0.088 | 0.312 |
| STD8 | 0.051<br>0.059 | 0.055 | 10.802 | | |

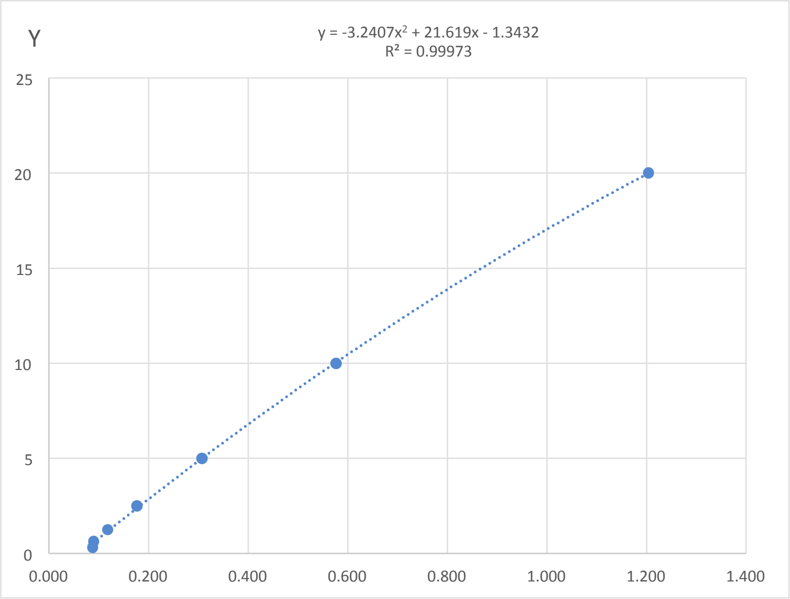


| y=a+bx+cx^2 | |
|---|---|
| a | -1.3432 |
| b | 21.6190 |
| c | -3.2407 |
| R^2 | 1.000 |

| DS | | | | | | |
|---|---|---|---|---|---|---|
| **Samples** | **O.D.** | **O.D. mean** | **%CV** | **O.D. mean without background (x)** | **Concentration (y)** | **Concentration (ng/mL)** |
| DSP20091 | 0.180<br>0.210 | 0.195 | 10.752 | 0.140 | 1.626 | 8.128 |
| DSP20207 | 0.215<br>0.225 | 0.220 | 3.273 | 0.165 | 2.131 | 10.656 |
| DSP20200 | 0.174<br>0.197 | 0.186 | 9.131 | 0.131 | 1.426 | 7.132 |
| DSP20144 | 0.190<br>0.216 | 0.203 | 8.955 | 0.148 | 1.783 | 8.916 |
| DSP20374 | 0.160<br>0.177 | 0.168 | 7.113 | 0.114 | 1.071 | 5.357 |
| DSP20113 | 0.172<br>0.214 | 0.193 | 15.098 | 0.138 | 1.580 | 7.900 |
| DSP20360 | 0.225<br>0.202 | 0.213 | 7.496 | 0.158 | 2.000 | 10.002 |
| DSP20077 | 0.319<br>0.229 | 0.274 | 23.430 | 0.219 | 3.239 | 16.194 |
| DSP20096 | 0.173<br>0.206 | 0.190 | 12.409 | 0.135 | 1.510 | 7.552 |

| | | | | | | |
|---|---|---|---|---|---|---|
| DSP20264 | 0.174<br>0.198 | 0.186 | 8.980 | 0.131 | 1.440 | 7.202 |
| DSP20189 | 0.136<br>0.171 | 0.153 | 16.088 | 0.098 | 0.755 | 3.774 |
| DSP19993 | 0.127<br>0.207 | 0.167 | 33.892 | 0.112 | 1.044 | 5.220 |
| DSP20381 | 0.151<br>0.199 | 0.175 | 19.337 | 0.120 | 1.211 | 6.055 |
| DSP20086 | 0.204<br>0.156 | 0.180 | 18.691 | 0.125 | 1.311 | 6.553 |
| DSP20080 | 0.175<br>0.237 | 0.206 | 21.427 | 0.151 | 1.845 | 9.225 |
| DSP20353 | 0.228<br>0.240 | 0.234 | 3.495 | 0.179 | 2.431 | 12.153 |
| DSP19948 | 0.261<br>0.204 | 0.233 | 17.304 | 0.178 | 2.399 | 11.994 |
| DSP20327 | 0.159<br>0.215 | 0.187 | 20.895 | 0.132 | 1.457 | 7.284 |
| DSP20242 | 0.225<br>0.151 | 0.188 | 27.768 | 0.133 | 1.480 | 7.402 |
| DSP20062 | 0.225<br>0.198 | 0.211 | 9.227 | 0.157 | 1.961 | 9.806 |
| DSP20363 | 0.148<br>0.196 | 0.172 | 19.600 | 0.117 | 1.152 | 5.761 |
| DSP19489 | 0.177<br>0.194 | 0.185 | 6.546 | 0.131 | 1.425 | 7.125 |
| DSP20095 | 0.182<br>0.151 | 0.167 | 13.077 | 0.112 | 1.037 | 5.186 |
| DSP19957 | 0.175<br>0.164 | 0.170 | 4.779 | 0.115 | 1.095 | 5.474 |
| DSP20136 | 0.157<br>0.150 | 0.154 | 3.062 | 0.099 | 0.765 | 3.824 |
| DSP20127 | 0.190<br>0.137 | 0.163 | 22.950 | 0.108 | 0.964 | 4.818 |
| DSP20205 | 0.143<br>0.141 | 0.142 | 0.779 | 0.087 | | |
| DSP20380 | 0.233<br>0.160 | 0.196 | 26.374 | 0.141 | 1.647 | 8.235 |
| DSP20139 | 0.283<br>0.204 | 0.243 | 22.970 | 0.188 | 2.614 | 13.070 |
| DSP20178 | 0.124<br>0.117 | 0.121 | 4.319 | 0.066 | | |
| DSP20325 | 0.125<br>0.139 | 0.132 | 7.658 | 0.077 | | |

| control | | | | | | |
|---|---|---|---|---|---|---|
| **Samples** | **O.D.** | **O.D. mean** | **%CV** | **O.D. mean without background (x)** | **Concentration (y)** | **Concentration (ng/mL)** |
| nP19968 | 0.162<br>0.157 | 0.159 | 1.846 | 0.105 | 0.884 | 4.418 |
| nP19461 | 0.114<br>0.134 | 0.124 | 10.911 | 0.069 | | |
| nP19653 | 0.113<br>0.137 | 0.125 | 13.264 | 0.070 | | |
| nP19454 | 0.124<br>0.144 | 0.134 | 10.490 | 0.079 | | |
| nP19467 | 0.118<br>0.140 | 0.129 | 11.977 | 0.074 | | |
| nP19487 | 0.199<br>0.175 | 0.187 | 9.126 | 0.132 | 1.461 | 7.303 |

| Standard | | | | | |
|---|---|---|---|---|---|
| **Samples** | **O.D.** | **O.D. mean** | **%CV** | **O.D. mean without background (x)** | **Concentration (ng/mL) (y)** |
| STD1 | 1.518<br>2.018 | 1.768 | 20.013 | 1.714 | 20 |
| STD2 | 0.865<br>0.985 | 0.925 | 9.167 | 0.872 | 10 |
| STD3 | 0.548<br>0.547 | 0.547 | 0.068 | 0.494 | 5 |
| STD4 | 0.344<br>0.335 | 0.340 | 1.855 | 0.286 | 2.5 |
| STD5 | 0.303<br>0.293 | 0.298 | 2.422 | 0.244 | 1.25 |
| STD6 | 0.189<br>0.204 | 0.196 | 5.330 | 0.143 | 0.625 |
| STD7* | 0.195<br>0.217 | 0.206 | 7.841 | 0.152 | 0.312 |
| STD8 | 0.048<br>0.060 | 0.054 | 16.031 | | |

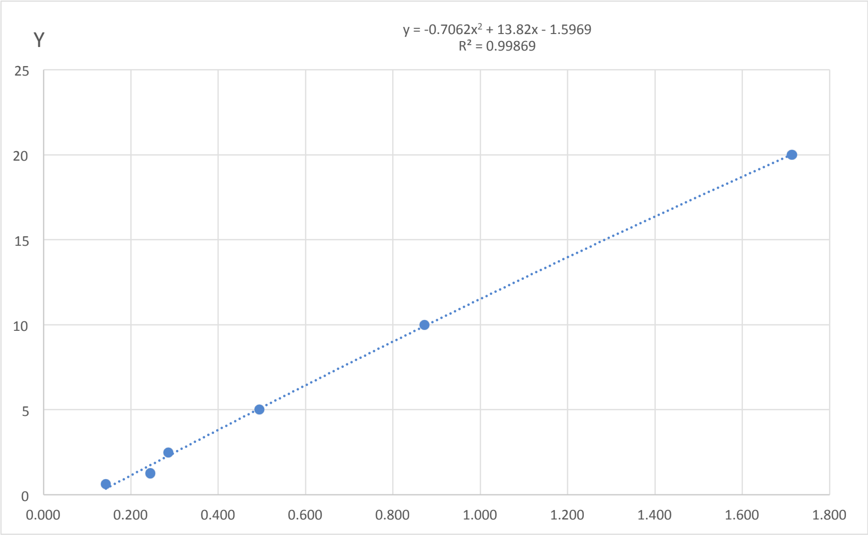


| y=a+bx+cx^2 | |
|---|---|
| a | -0.706 |
| b | 13.820 |
| c | -1.597 |
| R^2 | 0.999 |

| DS | | | | | | |
|---|---|---|---|---|---|---|
| **Samples** | **O.D.** | **O.D. mean** | **%CV** | **O.D. mean without background (x)** | **Concentration (y)** | **Concentration (ng/mL)** |
| DSP20359 | 0.290<br>0.258 | 0.274 | 8.216 | 0.220 | 2.262 | 11.308 |
| DSP20371 | 0.200<br>0.196 | 0.198 | 1.426 | 0.145 | 1.260 | 6.299 |
| DSP20376 | 0.198<br>0.128 | 0.163 | 30.370 | 0.110 | | |
| DSP20383 | 0.208<br>0.197 | 0.203 | 3.953 | 0.149 | 1.318 | 6.592 |
| DSP19916 | 0.202<br>0.160 | 0.181 | 16.572 | 0.127 | | |
| DSP19939 | 0.191<br>0.133 | 0.162 | 25.270 | 0.108 | | |
| DSP19952 | 0.219<br>0.199 | 0.209 | 6.515 | 0.155 | 1.400 | 7.000 |
| DSP19956 | 0.200<br>0.204 | 0.202 | 1.454 | 0.148 | 1.309 | 6.543 |
| DSP19989 | 0.313<br>0.322 | 0.317 | 1.860 | 0.264 | 2.827 | 14.136 |
| DSP20018 | 0.253<br>0.212 | 0.233 | 12.485 | 0.179 | 1.718 | 8.592 |
| DSP20034 | 0.220<br>0.199 | 0.210 | 6.905 | 0.156 | 1.410 | 7.052 |
| DSP20082 | 0.226<br>0.181 | 0.204 | 15.634 | 0.150 | 1.329 | 6.644 |
| DSP20092 | 0.193<br>0.184 | 0.188 | 3.301 | 0.134 | | |

| | | | | | | |
|---|---|---|---|---|---|---|
| DSP20117 | 0.186<br>0.196 | 0.191 | 3.842 | 0.137 | | |
| DSP20121 | 0.187<br>0.201 | 0.194 | 5.057 | 0.140 | | |
| DSP20123 | 0.179<br>0.218 | 0.199 | 13.655 | 0.145 | 1.263 | 6.315 |
| DSP20140 | 0.306<br>0.296 | 0.301 | 2.515 | 0.247 | 2.613 | 13.064 |
| DSP20150 | 0.267<br>0.249 | 0.258 | 4.744 | 0.204 | 2.050 | 10.252 |
| DSP20155 | 0.197<br>0.210 | 0.204 | 4.566 | 0.150 | 1.332 | 6.662 |
| DSP20161 | 0.194<br>0.200 | 0.197 | 2.270 | 0.144 | 1.245 | 6.226 |
| DSP20164 | 0.193<br>0.251 | 0.222 | 18.487 | 0.168 | 1.575 | 7.877 |
| DSP20187 | 0.178<br>0.172 | 0.175 | 2.473 | 0.121 | | |
| DSP20194 | 0.219<br>0.174 | 0.197 | 16.167 | 0.143 | 1.235 | 6.176 |
| DSP20214 | 0.212<br>0.184 | 0.198 | 9.835 | 0.145 | 1.258 | 6.289 |
| DSP20220 | 0.345<br>0.318 | 0.332 | 5.689 | 0.278 | 3.011 | 15.055 |
| DSP20236 | 0.236<br>0.222 | 0.229 | 4.252 | 0.175 | 1.667 | 8.333 |
| DSP20257 | 0.191<br>0.154 | 0.172 | 15.120 | 0.119 | | |
| DSP20271 | 0.272<br>0.186 | 0.229 | 26.578 | 0.175 | 1.667 | 8.335 |
| DSP20274 | 0.193<br>0.201 | 0.197 | 2.825 | 0.143 | 1.237 | 6.185 |
| DSP20290 | 0.214<br>0.227 | 0.220 | 4.067 | 0.166 | 1.550 | 7.752 |
| DSP20294 | 0.139<br>0.176 | 0.158 | 16.741 | 0.104 | | |
| DSP20296 | 0.152<br>0.233 | 0.192 | 29.791 | 0.138 | | |
| DSP20368 | 0.263<br>0.155 | 0.209 | 36.492 | 0.155 | 1.402 | 7.011 |

| control | | | | | | |
|---|---|---|---|---|---|---|
| **Samples** | **O.D.** | **O.D. mean** | **%CV** | **O.D. mean without background (x)** | **Concentration (y)** | **Concentration (ng/mL)** |
| nP19460 | 0.188<br>0.159 | 0.174 | 11.561 | 0.120 | | |
| nP20231 | 0.218<br>0.133 | 0.176 | 34.228 | 0.122 | | |
| nP20045 | 0.191<br>0.160 | 0.175 | 12.516 | 0.121 | | |
| nP19946 | 0.184<br>0.132 | 0.158 | 23.170 | 0.104 | | |
| nP20272 | 0.134<br>0.183 | 0.159 | 21.805 | 0.105 | | |
| nP19458 | 0.143<br>0.172 | 0.157 | 13.075 | 0.104 | | |

| Standard | | | | | |
|---|---|---|---|---|---|
| Samples | O.D. | O.D. mean | %CV | O.D. mean without background (x) | Concentration (ng/mL) (y) |
| STD1 | 1.895<br>2.069 | 1.982 | 6.202 | 1.931 | 20 |
| STD2 | 1.045<br>1.000 | 1.023 | 3.147 | 0.972 | 10 |
| STD3 | 0.642<br>0.624 | 0.633 | 1.942 | 0.582 | 5 |
| STD4 | 0.324<br>0.351 | 0.338 | 5.543 | 0.287 | 2.5 |
| STD5 | 0.227<br>0.239 | 0.233 | 3.617 | 0.182 | 1.25 |
| STD6 | 0.187<br>0.201 | 0.194 | 5.329 | 0.143 | 0.625 |
| STD7 | 0.165<br>0.162 | 0.164 | 1.387 | 0.113 | 0.312 |
| STD8 | 0.049<br>0.052 | 0.051 | 4.861 | | |

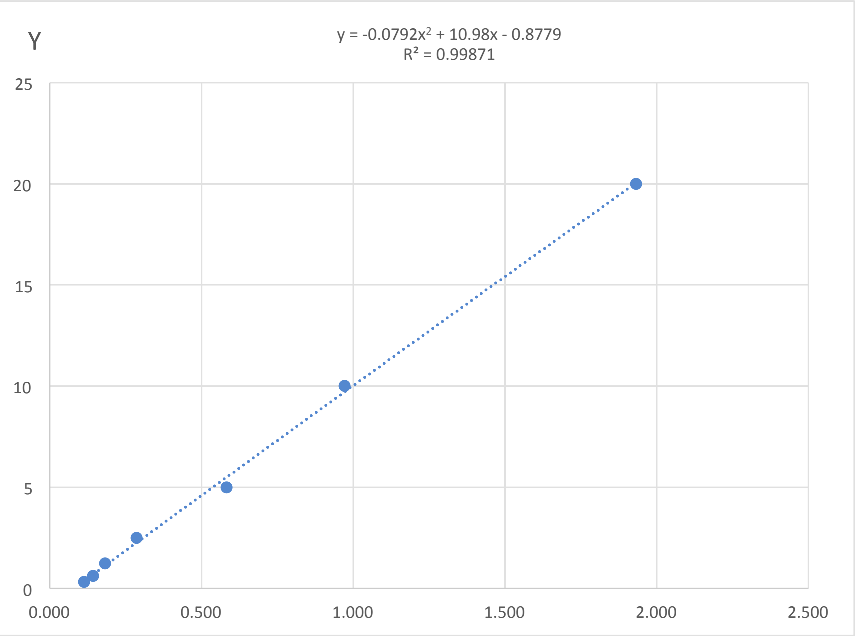


| y=a+bx+cx^2 | |
|---|---|
| a | -0.874 |
| b | 10.980 |
| c | -0.079 |
| R^2 | 0.999 |

| DS | | | | | | |
|---|---|---|---|---|---|---|
| Samples | O.D. | O.D. mean | %CV | O.D. mean without background (x) | Concentration (y) | Concentration (ng/mL) |
| DSP20379 | 0.248<br>0.323 | 0.285 | 18.553 | 0.235 | 1.699 | 8.494 |
| DSP20387 | 0.251<br>0.313 | 0.282 | 15.535 | 0.231 | 1.661 | 8.307 |
| DSP20388 | 0.208<br>0.181 | 0.195 | 9.979 | 0.144 | 0.706 | 3.528 |
| DSP20183 | 0.247<br>0.332 | 0.290 | 20.583 | 0.239 | 1.745 | 8.727 |
| DSP20085 | 0.217<br>0.218 | 0.217 | 0.216 | 0.167 | 0.954 | 4.770 |
| DSP20103 | 0.212<br>0.225 | 0.219 | 4.119 | 0.168 | 0.968 | 4.841 |
| DSP20104 | 0.174<br>0.153 | 0.163 | 9.034 | 0.112 | | |

| | | | | | | |
|---|---|---|---|---|---|---|
| DSP20112 | 0.203<br>0.202 | 0.203 | 0.286 | 0.152 | 0.792 | 3.959 |
| DSP20132 | 0.215<br>0.246 | 0.231 | 9.568 | 0.180 | 1.101 | 5.507 |
| DSP20235 | 0.211<br>0.259 | 0.235 | 14.304 | 0.185 | 1.150 | 5.751 |
| DSP20279 | 0.235<br>0.310 | 0.273 | 19.464 | 0.222 | 1.560 | 7.800 |
| DSP20350 | 0.245<br>0.197 | 0.221 | 15.358 | 0.170 | 0.991 | 4.954 |
| DSP20366 | 0.239<br>0.276 | 0.257 | 10.180 | 0.206 | 1.389 | 6.943 |

| **control** | | | | | | |
|---|---|---|---|---|---|---|
| **Samples** | **O.D.** | **O.D. mean** | **%CV** | **O.D. mean without background (x)** | **Concentration (y)** | **Concentration (ng/mL)** |
| nP19481 | 0.171<br>0.167 | 0.169 | 1.668 | 0.118 | 0.425 | 2.126 |
| nP19367 | 0.180<br>0.160 | 0.170 | 8.615 | 0.119 | 0.435 | 2.177 |
| nP19420 | 0.166<br>0.185 | 0.176 | 7.492 | 0.125 | 0.497 | 2.483 |
| nP19469 | 0.260<br>0.181 | 0.221 | 25.467 | 0.170 | 0.991 | 4.953 |

| Standard | | | | | |
|---|---|---|---|---|---|
| **Samples** | **O.D.** | **O.D. mean** | **%CV** | **O.D. mean without background (x)** | **Concentration (ng/mL) (y)** |
| STD1 | 1.715<br>1.732 | 1.723 | 0.691 | 1.680 | 20 |
| STD2 | 0.690<br>0.721 | 0.705 | 3.030 | 0.662 | 10 |
| STD3 | 0.446<br>0.478 | 0.462 | 4.974 | 0.418 | 5 |
| STD4 | 0.221<br>0.269 | 0.245 | 13.941 | 0.201 | 2.5 |
| STD5 | 0.166<br>0.178 | 0.172 | 4.808 | 0.128 | 1.25 |
| STD6 | 0.112<br>0.124 | 0.118 | 6.888 | 0.074 | 0.625 |
| STD7 | 0.086<br>0.098 | 0.092 | 9.367 | 0.048 | 0.312 |
| STD8 | 0.043<br>0.044 | 0.044 | 2.458 | | |

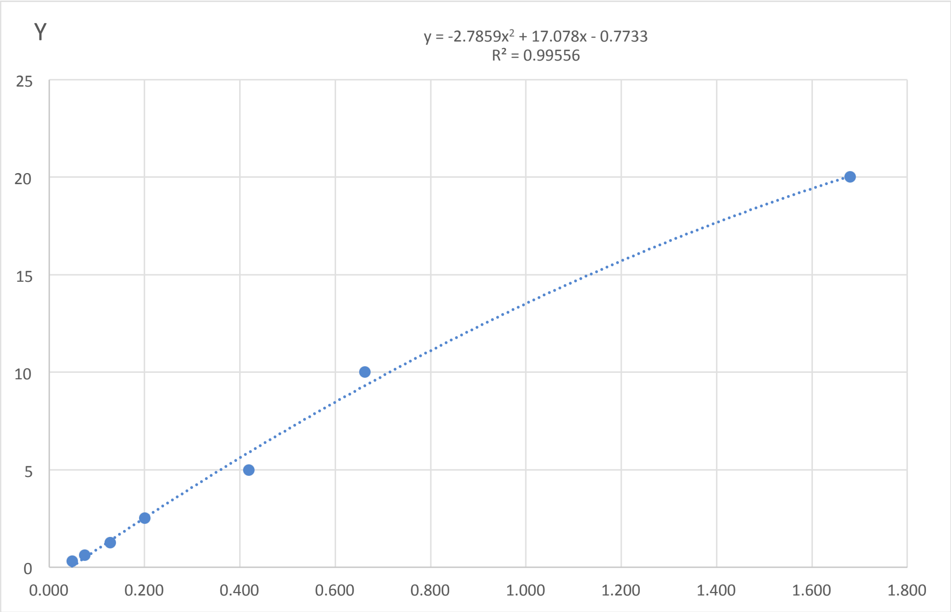


| y=a+bx+cx^2 | |
|---|---|
| a | -0.7733 |
| b | 17.078 |
| c | -2.7859 |
| R^2 | 0.9956 |

| DS | | | | | | |
|---|---|---|---|---|---|---|
| **Samples** | **O.D.** | **O.D. mean** | **%CV** | **O.D. mean without background (x)** | **Concentration (y)** | **Concentration (ng/mL)** |
| DSP19971 | 0.109<br>0.128 | 0.118 | 11.738 | 0.075 | 0.488 | 1.463 |
| DSP20209 | 0.138<br>0.088 | 0.113 | 31.002 | 0.069 | 0.398 | 1.194 |

| Standard | | | | | |
|---|---|---|---|---|---|
| Samples | O.D. | O.D. mean | %CV | O.D. mean without background (x) | Concentration (ng/mL) (y) |
| STD1 | 1.534<br>1.691 | 1.612 | 6.904 | 1.570 | 20 |
| STD2 | 0.841<br>0.778 | 0.810 | 5.516 | 0.767 | 10 |
| STD3 | 0.434<br>0.453 | 0.444 | 3.075 | 0.401 | 5 |
| STD4 | 0.251<br>0.285 | 0.268 | 8.833 | 0.226 | 2.5 |
| STD5 | 0.188<br>0.176 | 0.182 | 4.542 | 0.140 | 1.25 |
| STD6 | 0.128<br>0.133 | 0.130 | 2.798 | 0.088 | 0.625 |
| STD7 | 0.096<br>0.097 | 0.096 | 0.675 | 0.054 | 0.312 |
| STD8 | 0.041<br>0.043 | 0.042 | 2.975 | 0.000 | |

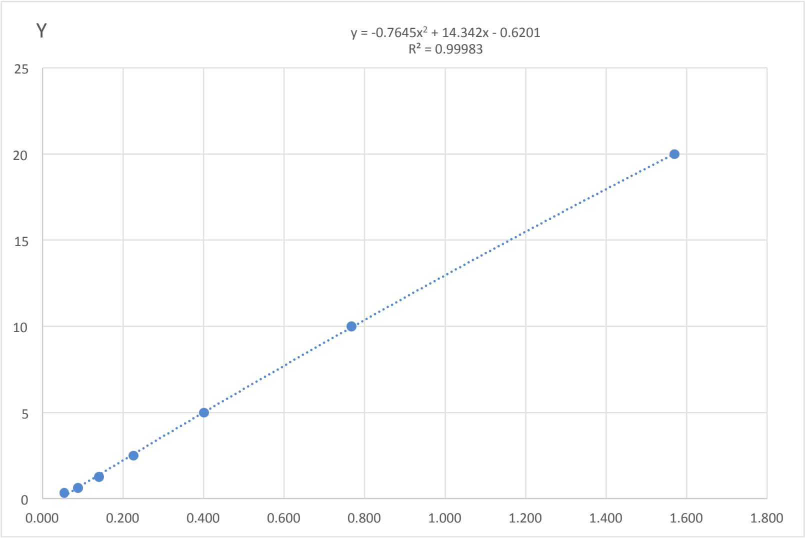


| y=a+bx+cx^2 | |
|---|---|
| a | -0.6201 |
| b | 14.3420 |
| c | -0.7645 |
| R^2 | 0.9998 |

| control | | | | | | |
|---|---|---|---|---|---|---|
| Samples | O.D. | O.D. mean | %CV | O.D. mean without background (x) | Concentration (y) | Concentration (ng/mL) |
| nP19454 | 0.115<br>0.144 | 0.130 | 15.721 | 0.088 | 0.629 | 1.887 |
| nP19653 | 0.090<br>0.110 | 0.100 | 14.642 | 0.058 | 0.203 | 0.609 |
| nP19691 | 0.114<br>0.113 | 0.113 | 0.797 | 0.071 | 0.396 | 1.188 |
| nP19812 | 0.128<br>0.133 | 0.131 | 2.819 | 0.088 | 0.641 | 1.924 |
| nP19854 | 0.120<br>0.143 | 0.132 | 12.379 | 0.089 | 0.656 | 1.969 |
| nP19902 | 0.114<br>0.106 | 0.110 | 4.914 | 0.068 | 0.349 | 1.047 |
| nP20045 | 0.115<br>0.119 | 0.117 | 2.430 | 0.075 | 0.447 | 1.342 |
| nP20246 | 0.116<br>0.127 | 0.122 | 6.329 | 0.079 | 0.512 | 1.537 |
| nP20284 | 0.092<br>0.110 | 0.101 | 12.874 | 0.059 | 0.217 | 0.651 |
| nP20341 | 0.084<br>0.117 | 0.101 | 23.231 | 0.058 | 0.216 | 0.649 |
| nP20398 | 0.093<br>0.108 | 0.100 | 10.304 | 0.058 | 0.209 | 0.627 |
| nP20399 | 0.078<br>0.099 | 0.089 | 17.249 | | | |
