## Supplementary Table 5 for "One-carbon pathway metabolites are altered in the plasma of subjects with Down syndrome: relation to chromosomal dosage"

**Supplementary Table 5. Analysis of SAM ELISA assays**. The table "Standard" shows the two absorbance (O.D.) detected for each standard sample. In the "O.D. mean" column, the average O.D. is reported. The "%CV" column represent the Coefficient of Variability between the two O.D. measurment expressed as percentage. The "1/O.D. mean (x)" column reports the inverse of O.D. mean values. The "Concentration (ng/mL) (y)" column reports the known concentration of each standard sample supplied by ELISA kit. In order to build the standard curve, we plotted "1/O.D. mean (x)" of standards in the x-axis and "Concentration (ng/mL) (y)" of standards in the y-axis. The polynomial equation (y=a+bx+cx^2) reported in the graph was used to determine metabolites concentration of plasma samples ("Concentration (ng/mL) (y)"), using interpolation of the inverse of O.D. mean values reported in "1/O.D. mean (x)" column in tables "DS" and "control". The results are reported in "Concentration (ng/mL)" column in tables "DS" and "control". In tables "DS" and "control" the O.D. mean values which are higher or lower than the O.D. mean values of standard sample ranges are reported in red and these values were not considered for the statistical analysis.

| Standard | | | | | |
|---|---|---|---|---|---|
| **Samples** | **O.D.** | **O.D. mean** | **%CV** | **1/O.D. mean (x)** | **Concentration (µg/mL) (y)** |
| STD1 | 0.103<br>0.101 | 0.102 | 0.781 | 9.804 | 25.000 |
| STD2 | 0.142<br>0.148 | 0.145 | 2.997 | 6.885 | 12.500 |
| STD3 | 0.216<br>0.195 | 0.205 | 6.899 | 4.867 | 6.250 |
| STD4 | 0.315<br>0.319 | 0.317 | 0.879 | 3.156 | 3.125 |
| STD5 | 0.472<br>0.485 | 0.478 | 1.938 | 2.091 | 1.560 |
| STD6 | 0.723<br>0.679 | 0.701 | 4.397 | 1.426 | 0.781 |
| STD7 | 0.974<br>0.952 | 0.963 | 1.577 | 1.039 | 0.390 |
| STD8 | 1.337<br>1.381 | 1.359 | 2.284 | 0.736 | 0.000 |

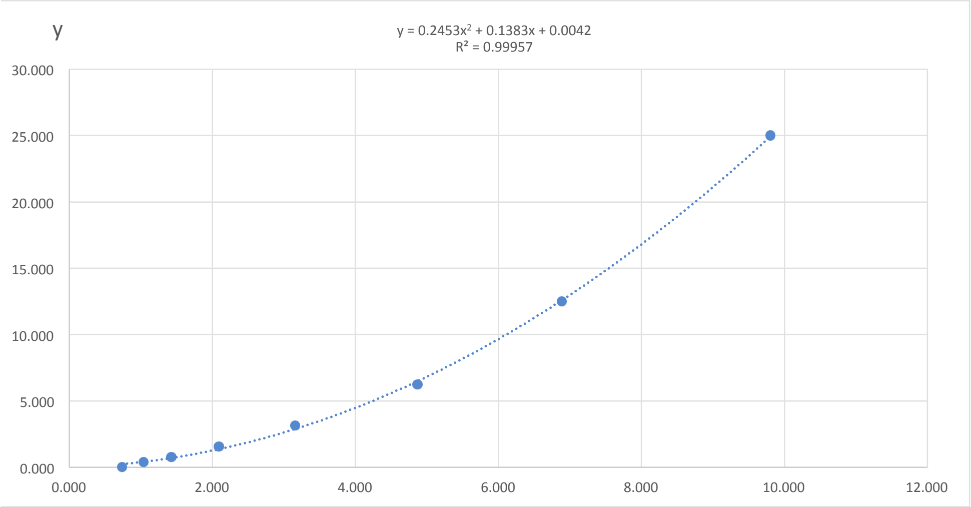


| y=a+bx+cx^2 | |
|---|---|
| a | 0.0042 |
| b | 0.1383 |
| c | 0.2453 |
| R^2 | 1.000 |

| DS | | | | | |
|---|---|---|---|---|---|
| **Samples** | **O.D.** | **O.D. mean** | **%CV** | **1/O.D. mean (x)** | **Concentration (µg/mL) (y)** |
| DSP20080 | 0.170<br>0.173 | 0.171 | 1.449 | 5.831 | 9.152 |
| DSP20086 | 0.161<br>0.183 | 0.172 | 9.120 | 5.811 | 9.090 |
| DSP20091 | 0.209<br>0.186 | 0.198 | 8.259 | 5.054 | 6.970 |
| DSP20096 | 0.220<br>0.223 | 0.222 | 0.907 | 4.507 | 5.610 |
| DSP20110 | 0.174<br>0.129 | 0.151 | 20.962 | 6.615 | 11.653 |
| DSP20127 | 0.227<br>0.169 | 0.198 | 20.876 | 5.052 | 6.964 |
| DSP20136 | 0.191<br>0.194 | 0.192 | 1.006 | 5.200 | 7.356 |
| DSP20139 | 0.142<br>0.218 | 0.180 | 29.628 | 5.553 | 8.338 |

| | | | | | |
|---|---|---|---|---|---|
| DSP20155 | 0.194 | 0.194 | 0.117 | 5.158 | 7.243 |
| | 0.194 | | | | |
| DSP20164 | 0.223 | 0.207 | 10.940 | 4.830 | 6.395 |
| | 0.191 | | | | |
| DSP20200 | 0.127 | 0.142 | 14.640 | 7.058 | 13.200 |
| | 0.156 | | | | |
| DSP20242 | 0.164 | 0.157 | 5.819 | 6.358 | 10.801 |
| | 0.151 | | | | |
| DSP20257 | 0.183 | 0.183 | 0.180 | 5.457 | 8.064 |
| | 0.183 | | | | |
| DSP20264 | 0.153 | 0.157 | 4.224 | 6.354 | 10.788 |
| | 0.162 | | | | |
| DSP20267 | 0.120 | 0.109 | 15.122 | 9.204 | 22.056 |
| | 0.097 | | | | |
| DSP20294 | 0.173 | 0.184 | 8.080 | 5.449 | 8.042 |
| | 0.194 | | | | |
| DSP20299 | 0.255 | 0.264 | 4.842 | 3.789 | 4.049 |
| | 0.273 | | | | |
| DSP20327 | 0.117 | 0.120 | 3.311 | 8.335 | 18.197 |
| | 0.123 | | | | |
| DSP20333 | 0.154 | 0.154 | 0.198 | 6.505 | 11.284 |
| | 0.154 | | | | |
| DSP20353 | 0.163 | 0.165 | 1.166 | 6.068 | 9.877 |
| | 0.166 | | | | |
| DSP20371 | 0.075 | 0.075 | 0.655 | | |
| | 0.076 | | | | |
| DSP20374 | 0.130 | 0.132 | 2.368 | 7.557 | 15.059 |
| | 0.135 | | | | |
| DSP20379 | 0.182 | 0.164 | 15.609 | 6.116 | 10.025 |
| | 0.145 | | | | |
| DSP20381 | 0.164 | 0.162 | 1.643 | 6.173 | 10.207 |
| | 0.160 | | | | |
| DSP20383 | 0.134 | 0.124 | 11.117 | 8.038 | 16.963 |
| | 0.115 | | | | |

| **control** | | | | | |
|---|---|---|---|---|---|
| **Samples** | **O.D.** | **O.D. mean** | **%CV** | **1/O.D. mean (x)** | **Concentration (µg/mL) (y)** |
| nP19450 | 0.164 | 0.161 | 2.844 | 6.215 | 10.338 |
| | 0.158 | | | | |
| nP19454 | 0.119 | 0.115 | 5.198 | 8.676 | 19.670 |
| | 0.111 | | | | |
| nP19653 | 0.280 | 0.242 | 22.491 | 4.134 | 4.768 |
| | 0.203 | | | | |
| nP19691 | 0.253 | 0.235 | 10.800 | 4.253 | 5.029 |
| | 0.217 | | | | |
| nP19812 | 0.309 | 0.305 | 1.956 | 3.280 | 3.098 |
| | 0.301 | | | | |
| nP19854 | 0.173 | 0.173 | 0.111 | 5.782 | 9.006 |
| | 0.173 | | | | |
| nP19902 | 0.249 | 0.260 | 5.696 | 3.851 | 4.174 |
| | 0.270 | | | | |
| nP20045 | 0.220 | 0.231 | 7.085 | 4.321 | 5.182 |
| | 0.243 | | | | |
| nP20231 | 0.252 | 0.277 | 12.749 | 3.614 | 3.707 |
| | 0.302 | | | | |
| nP20246 | 0.155 | 0.102 | 72.545 | 9.765 | 24.744 |
| | 0.050 | | | | |
| nP20284 | 0.223 | 0.228 | 2.782 | 4.389 | 5.337 |
| | 0.232 | | | | |
| nP20341 | 0.316 | 0.364 | 18.393 | 2.749 | 2.239 |
| | 0.411 | | | | |
| nP20343 | 0.213 | 0.202 | 7.771 | 4.948 | 6.694 |
| | 0.191 | | | | |
| nP20398 | 0.156 | 0.155 | 0.550 | 6.454 | 11.115 |
| | 0.154 | | | | |
| nP20399 | 0.247 | 0.230 | 10.276 | 4.339 | 5.222 |
| | 0.214 | | | | |
