## Supplementary Table 6 for "One-carbon pathway metabolites are altered in the plasma of subjects with Down syndrome: relation to chromosomal dosage"

**Supplementary Table 6. Descriptive analysis of DS and control group.** The values of subjects with Down syndrome (DS) are reported on the left of the table and the values of normal control subjects (control) are reported on the right of the table. For each molecule the number (n) of samples valid or missing are reported. Also the mean, modal value, standard deviation (SD), variance, range minimum and maximum concentration values are shown. Concerning the modal value, the smallest value is reported. In table A strong outliers are included in the analysis and in table B strong outliers are excluded from the analysis.

**A**

| | | DS | | | | | | | | control | | | | |
|---|---|---|---|---|---|---|---|---|---|---|---|---|---|---|
| | | Folic acid (ng/mL) | Vit B12 (pg/mL) | Hcy (µmol/L) | THF (ng/mL) | 5-methyl-THF (ng/mL) | 5-formyl-THF (pg/mL) | SAH (ng/mL) | SAM (µg/mL) | THF (ng/mL) | 5-methyl-THF (ng/mL) | 5-formyl-THF (pg/mL) | SAH (ng/mL) | SAM (µg/mL) |
| **n** | **Valid** | 143 | 155 | 42 | 108 | 140 | 80 | 94 | 24 | 41 | 34 | 21 | 20 | 15 |
| | **Missing** | 32 | 20 | 133 | 56 | 24 | 84 | 70 | 140 | 13 | 20 | 33 | 34 | 39 |
| **Mean** | | 8 | 347,37 | 9,079 | 41,574 | 50,247 | 157,028 | 6,588 | 10,308 | 57,783 | 45,369 | 145,917 | 2,180 | 8,022 |
| **Median** | | 7 | 329 | 8,35 | 33,955 | 49,766 | 138,091 | 6,298 | 9,514 | 51,471 | 42,744 | 139,144 | 1,812 | 5,222 |
| **Modal value** | | 4,7 | 345 | 8,3 | 3,820 | 24,217 | 119,681 | 4,768 | 4,049 | 8,562 | 26,911 | 123,569 | 0,609 | 2,239 |
| **SD** | | 3 | 175 | 4,1024 | 29,414 | 9,358 | 56,036 | 2,923 | 4,237 | 32,645 | 12,301 | 15,413 | 1,688 | 6,362 |
| **Variance** | | 12 | 31.105 | 16,83 | 865,199 | 87,573 | 3139,988 | 8,543 | 17,950 | 1065,665 | 151,320 | 237,570 | 2,848 | 40,481 |
| **Range** | | 18 | 1.395 | 21 | 134,102 | 46,358 | 460,237 | 15,001 | 18,007 | 106,992 | 45,877 | 55,701 | 6,693 | 22,505 |
| **Minimum** | | 2,5 | 106 | 2,1 | 3,820 | 24,217 | 119,681 | 1,194 | 4,049 | 8,562 | 26,911 | 123,569 | 0,609 | 2,239 |
| **Maximum** | | 20,6 | 1501 | 23,1 | 137,922 | 70,575 | 579,917 | 16,194 | 22,056 | 115,554 | 72,788 | 179,270 | 7,303 | 24,744 |

**B**

| | | DS | | | | | | | | control | | | | |
|---|---|---|---|---|---|---|---|---|---|---|---|---|---|---|
| | | Folic acid (ng/mL) | Vit B12 (pg/mL) | Hcy (µmol/L) | THF (ng/mL) | 5-methyl-THF (ng/mL) | 5-formyl-THF (pg/mL) | SAH (ng/mL) | SAM (µg/mL) | THF (ng/mL) | 5-methyl-THF (ng/mL) | 5-formyl-THF (pg/mL) | SAH (ng/mL) | SAM (µg/mL) |
| **n** | **Valid** | 143 | 154 | 42 | 108 | 140 | 79 | 94 | 24 | 41 | 34 | 21 | 19 | 15 |
| | **Missing** | 32 | 21 | 133 | 56 | 24 | 85 | 70 | 140 | 13 | 20 | 33 | 35 | 39 |
| **Mean** | | 8 | 339,88 | 9,079 | 41,574 | 50,247 | 151,675 | 6,588 | 10,308 | 57,783 | 45,369 | 145,917 | 1,911 | 8,022 |
| **Median** | | 7 | 329 | 8,35 | 33,955 | 49,766 | 137,955 | 6,298 | 9,514 | 51,471 | 42,744 | 139,144 | 1,736 | 5,222 |
| **Modal value** | | 4,7 | 345 | 8,3 | 3,820 | 24,217 | 119,681 | 4,768 | 4,049 | 8,562 | 26,911 | 123,569 | 0,609 | 2,239 |
| **SD** | | 3 | 150 | 4,1024 | 29,414 | 9,358 | 29,300 | 2,923 | 4,237 | 32,645 | 12,301 | 15,413 | 1,213 | 6,362 |
| **Variance** | | 12 | 22.553 | 16,83 | 865,199 | 87,573 | 858,463 | 8,543 | 17,950 | 1065,665 | 151,320 | 237,570 | 1,472 | 40,481 |
| **Range** | | 18 | 901 | 21 | 134,102 | 46,358 | 139,777 | 15,001 | 18,007 | 106,992 | 45,877 | 55,701 | 4,344 | 22,505 |
| **Minimum** | | 2,5 | 106 | 2,1 | 3,820 | 24,217 | 119,681 | 1,194 | 4,049 | 8,562 | 26,911 | 123,569 | 0,609 | 2,239 |
| **Maximum** | | 20,6 | 1007 | 23,1 | 137,922 | 70,575 | 259,457 | 16,194 | 22,056 | 115,554 | 72,788 | 179,270 | 4,953 | 24,744 |
