## Supplementary Table 7 for "One-carbon pathway metabolites are altered in the plasma of subjects with Down syndrome: relation to chromosomal dosage"

**Supplementary Table 7. Unpaired student t-test between DS and control concentration values of each metabolite.** Significant p-values (<0.05) are reported in red in the first line. T-value, degrees of freedom (df) and standard deviation (SD) are reported in the other lines. In table A strong outliers are included in the analysis and in table B strong outliers are excluded from the analysis.

**A**

| | THF | 5-methyl-THF | 5-formyl-THF | SAH | SAM |
|---|---|---|---|---|---|
| **p-value** | 0,0041 | 0,015 | 0,372 | 0,0001 | 0,1853 |
| **t-value** | 2,9137 | 2,5537 | 0,8967 | 6,5027 | 1,3499 |
| **df** | 147 | 172 | 99 | 112 | 37 |
| **SD** | 5,563 | 1,91 | 12,39 | 0,678 | 1,694 |

**B**

| | THF | 5-methyl-THF | 5-formyl-THF | SAH | SAM |
|---|---|---|---|---|---|
| **p-value** | 0,0041 | 0,015 | 0,3881 | 0,0001 | 0,1853 |
| **t-value** | 2,9137 | 2,5537 | 0,8669 | 6,8373 | 1,3499 |
| **df** | 147 | 172 | 98 | 111 | 37 |
| **SD** | 5,563 | 1,91 | 6,641 | 0,684 | 1,694 |
