## Supplementary Table 8 for "One-carbon pathway metabolites are altered in the plasma of subjects with Down syndrome: relation to chromosomal dosage"

**Supplementary Table 8. Bivariate correlation between age and each concentration levels in DS and control groups.** The table reports for each metabolite the results of the statistical analyses in DS and control groups indicated by Pearson correlation coefficient (r) and two-tailed significance (p-value) values. Significant r and p-values (< 0.05) are reported in red. In table A strong outliers are included in the analysis and in table B strong outliers are excluded from the analysis.

**A**

| | DS | | | | | | | | control | | | | |
|---|---|---|---|---|---|---|---|---|---|---|---|---|---|
| | Folic acid | Vit B12 | Hcy | THF | 5-methyl-THF | 5-formyl-THF | SAH | SAM | THF | 5-methyl-THF | 5-formyl-THF | SAH | SAM |
| r | -0,144 | -0,258 | 0,593 | 0,089 | -0,085 | 0,099 | 0,152 | 0,099 | 0,395 | 0,271 | 0,239 | 0,334 | 0,115 |
| p-value | 0,085 | 0,001 | < 0.001 | 0,362 | 0,318 | 0,383 | 0,145 | 0,644 | 0,011 | 0,121 | 0,296 | 0,151 | 0,682 |

**B**

| | DS | | | | | | | | control | | | | |
|---|---|---|---|---|---|---|---|---|---|---|---|---|---|
| | Folic acid | Vit B12 | Hcy | THF | 5-methyl-THF | 5-formyl-THF | SAH | SAM | THF | 5-methyl-THF | 5-formyl-THF | SAH | SAM |
| r | -0,144 | -0,256 | 0,593 | 0,089 | -0,085 | 0,128 | 0,152 | 0,099 | 0,395 | 0,271 | 0,239 | 0,102 | 0,115 |
| p-value | 0,085 | 0,001 | < 0.001 | 0,362 | 0,318 | 0,261 | 0,145 | 0,644 | 0,011 | 0,121 | 0,296 | 0,678 | 0,682 |
