## Supplementary Table 10 for "One-carbon pathway metabolites are altered in the plasma of subjects with Down syndrome: relation to chromosomal dosage"

**Supplementary Table 10. Results of unpaired t-test between fasting/non-fasting state and concentration values of DS and control groups.** The values of subjects with Down syndrome (DS) are reported on the left of the table and the values of normal control subjects (control) are reported on the right of the table. T-value, degrees of freedom (df), bilateral p-value and standard error (Std error) are reported for each metabolite assuming or not equal variances. Significant p-values (< 0.05) are reported in red. In table A strong outliers are included in the analysis and in table B strong outliers are excluded from the analysis.

**A**

| DS | | | | | | control | | | | | |
|---|---|---|---|---|---|---|---|---|---|---|---|
| | | t-value | df | Bilateral p-value | Std error | | | t-value | df | Bilateral p-value | Std error |
| THF | Assumed equal variances | -0,683 | 104,00 | 0,496 | 5,795 | THF | Assumed equal variances | -0,859 | 38,000 | 0,396 | 11,284 |
| | Not assumed equal variances | -0,69 | 102,14 | 0,492 | 5,733 | | Not assumed equal variances | -0,889 | 22,647 | 0,383 | 10,896 |
| 5-methyl-THF | Assumed equal variances | -0,082 | 136 | 0,935 | 1,603 | 5-methyl-THF | Assumed equal variances | -0,337 | 32 | 0,739 | 4,847 |
| | Not assumed equal variances | -0,082 | 134,88 | 0,935 | 1,601 | | Not assumed equal variances | -0,35 | 15,316 | 0,731 | 4,658 |
| 5-formyl-THF | Assumed equal variances | -0,305 | 78 | 0,761 | 12,606 | 5-formyl-THF | Assumed equal variances | 0,355 | 19 | 0,726 | 7,083 |
| | Not assumed equal variances | -0,31 | 54,421 | 0,757 | 12,386 | | Not assumed equal variances | 0,375 | 17,453 | 0,712 | 6,711 |
| SAH | Assumed equal variances | -0,262 | 90 | 0,794 | 0,622 | SAH | Assumed equal variances | 1,716 | 17 | 0,104 | 0,923 |
| | Not assumed equal variances | -0,278 | 89,358 | 0,782 | 0,587 | | Not assumed equal variances | 2,784 | 14,172 | 0,015 | 0,569 |
| SAM | Assumed equal variances | 0,676 | 21,00 | 0,507 | 1,681 | SAM | Assumed equal variances | 0,207 | 12,00 | 0,839 | 2,544 |
| | Not assumed equal variances | 0,661 | 15,72 | 0,518 | 1,720 | | Not assumed equal variances | 0,277 | 10,84 | 0,825 | 2,322 |
| Folic acid | Assumed equal variances | -1,261 | 139 | 0,209 | 0,583 | | | | | | |
| | Not assumed equal variances | -1,263 | 135,34 | 0,209 | 0,582 | | | | | | |
| Vit B12 | Assumed equal variances | -2,92 | 151 | 0,004 | 28,020 | | | | | | |
| | Not assumed equal variances | -2,954 | 139,62 | 0,004 | 27,701 | | | | | | |
| Hcy | Assumed equal variances | 1,947 | 39 | 0,059 | 1,257 | | | | | | |
| | Not assumed equal variances | 2,002 | 37,089 | 0,053 | 1,223 | | | | | | |

**B**

| DS | | | | | | control | | | | | |
|---|---|---|---|---|---|---|---|---|---|---|---|
| | | t-value | df | Bilateral p-value | Std error | | | t-value | df | Bilateral p-value | Std error |
| THF | Assumed equal variances | -0,683 | 104,00 | 0,496 | 5,795 | THF | Assumed equal variances | -0,859 | 38,000 | 0,396 | 11,284 |
| | Not assumed equal variances | -0,69 | 102,14 | 0,492 | 5,733 | | Not assumed equal variances | -0,889 | 22,647 | 0,383 | 10,896 |
| 5-methyl-THF | Assumed equal variances | -0,082 | 136 | 0,935 | 1,603 | 5-methyl-THF | Assumed equal variances | -0,337 | 32 | 0,739 | 4,847 |
| | Not assumed equal variances | -0,082 | 134,88 | 0,935 | 1,601 | | Not assumed equal variances | -0,35 | 15,316 | 0,731 | 4,658 |
| 5-formyl-THF | Assumed equal variances | 1,013 | 77 | 0,314 | 6,592 | 5-formyl-THF | Assumed equal variances | 0,355 | 19 | 0,726 | 7,083 |
| | Not assumed equal variances | 1,012 | 75,781 | 0,315 | 6,601 | | Not assumed equal variances | 0,375 | 17,453 | 0,712 | 6,711 |

| | | | | | |
|---|---|---|---|---|---|
| **SAH** | Assumed equal variances | -0,262 | 90 | 0,794 | 0,622 |
| | Not assumed equal variances | -0,278 | 89,358 | 0,782 | 0,587 |
| **SAM** | Assumed equal variances | 0,676 | 21,00 | 0,507 | 1,681 |
| | Not assumed equal variances | 0,661 | 15,72 | 0,518 | 1,720 |
| **Folic acid** | Assumed equal variances | -1,261 | 139 | 0,209 | 0,583 |
| | Not assumed equal variances | -1,263 | 135,34 | 0,209 | 0,582 |
| **Vit B12** | Assumed equal variances | -2,818 | 150 | 0,005 | 23,966 |
| | Not assumed equal variances | -2,826 | 149,51 | 0,005 | 23,896 |
| **Hcy** | Assumed equal variances | 1,947 | 39 | 0,059 | 1,257 |
| | Not assumed equal variances | 2,002 | 37,089 | 0,053 | 1,223 |

| | | | | | |
|---|---|---|---|---|---|
| **SAH** | Assumed equal variances | 1,892 | 16 | 0,077 | 0,658 |
| | Not assumed equal variances | 2,634 | 9,676 | 0,026 | 0,472 |
| **SAM** | Assumed equal variances | 0,207 | 12,00 | 0,839 | 2,544 |
| | Not assumed equal variances | 0,277 | 10,84 | 0,825 | 2,322 |
