## Supplementary Table 11 for "One-carbon pathway metabolites are altered in the plasma of subjects with Down syndrome: relation to chromosomal dosage"

**Supplementary Table 11. Results of correlation analyses between concentration levels of each molecules and the concentration levels of all the other molecules.** The values of subjects with Down syndrome (DS) are reported on the left of the table and the values of normal control subjects (control) are reported on the right of the table. Pearson's correlation (r), two tailed-significance (p-value) and the number of plasma sample analysed (n) are reported for each metabolite. Significant p-values (< 0.05) with correlation are reported in red. A bivariate correlation analysis was performed for folic acid, vitamin B12 (divided in fasting and non-fasting groups), 5-methyl-THF, 5-formyl-THF, SAH and SAM. A partial correlation analysis checking for the effect of chronological age was performed for THF and homocysteine. In table A strong outliers are included in the analysis and in table B strong outliers are excluded from the analysis.

**A**

| | | DS | | | | | | | | | control | | | | |
|---|---|---|---|---|---|---|---|---|---|---|---|---|---|---|---|
| | | Folic acid | Hcy | Vit B12 | | 5-formyl-THF | THF | 5-methyl-THF | SAH | SAM | 5-formyl-THF | THF | 5-methyl-THF | SAH | SAM |
| | | | | Fasting | Non-fasting | | | | | | | | | | |
| **5-formyl-THF** | r | -0,026 | 0,186 | -0,044 | -0,149 | | -0,148 | 0,163 | -0,294 | 0,81 | | 0,387 | 0,463 | -0,531 | -0,301 |
| | p-value | 0,837 | 0,508 | 0,794 | 0,393 | | 0,305 | 0,168 | 0,047 | 0,003 | | 0,125 | 0,095 | 0,175 | 0,612 |
| | n | 66 | 16 | 38 | 35 | | 53 | 73 | 46 | 11 | | 18 | 14 | 8 | 7 |
| **THF** | r | -0,057 | -0,217 | -0,162 | -0,143 | -0,148 | | 0,252 | 0,07 | -0,186 | 0,387 | | 0,299 | -0,257 | 0,079 |
| | p-value | 0,584 | 0,308 | 0,276 | 0,299 | 0,305 | | 0,017 | 0,529 | 0,384 | 0,125 | | 0,146 | 0,374 | 0,789 |
| | n | 96 | 25 | 47 | 55 | 53 | | 93 | 87 | 24 | 18 | | 26 | 15 | 14 |
| **5-methyl-THF** | r | 0,071 | 0,028 | 0,058 | 0,188 | 0,163 | 0,252 | | -0,029 | 0,047 | 0,463 | 0,299 | | 0,103 | 0,464 |
| | p-value | 0,434 | 0,88 | 0,643 | 0,133 | 0,168 | 0,017 | | 0,803 | 0,828 | 0,095 | 0,146 | | 0,826 | 0,294 |
| | n | 124 | 32 | 66 | 65 | 73 | 93 | | 79 | 24 | 14 | 26 | | 7 | 7 |
| **SAH** | r | 0,075 | -0,067 | 0,041 | -0,214 | -0,294 | 0,07 | -0,029 | | 0,045 | -0,531 | -0,257 | 0,103 | | 0,309 |
| | p-value | 0,508 | 0,798 | 0,806 | 0,14 | 0,047 | 0,529 | 0,803 | | 0,845 | 0,175 | 0,374 | 0,826 | | 0,329 |
| | n | 81 | 18 | 39 | 49 | 46 | 87 | 79 | | 21 | 8 | 15 | 7 | | 12 |
| **SAM** | r | 0,004 | | 0,004 | -0,628 | 0,81 | -0,186 | 0,047 | 0,045 | | -0,301 | 0,079 | 0,464 | 0,309 | |
| | p-value | 0,986 | | 0,99 | 0,029 | 0,003 | 0,384 | 0,828 | 0,845 | | 0,612 | 0,789 | 0,294 | 0,329 | |
| | n | 24 | | 11 | 12 | 11 | 24 | 24 | 21 | | 7 | 14 | 7 | 12 | |

**B**

| | | DS | | | | | | | | | control | | | | |
|---|---|---|---|---|---|---|---|---|---|---|---|---|---|---|---|
| | | Folic acid | Hcy | Vit B12 | | 5-formyl-THF | THF | 5-methyl-THF | SAH | SAM | 5-formyl-THF | THF | 5-methyl-THF | SAH | SAM |
| | | | | Fasting | Non-fasting | | | | | | | | | | |
| **5-formyl-THF** | r | 0,062 | 0,186 | -0,044 | 0,159 | | -0,148 | 0,163 | -0,294 | 0,81 | | 0,387 | 0,463 | -0,687 | -0,301 |
| | p-value | 0,624 | 0,508 | 0,794 | 0,369 | | 0,305 | 0,168 | 0,047 | 0,003 | | 0,125 | 0,095 | 0,088 | 0,612 |
| | n | 65 | 16 | 38 | 34 | | 53 | 73 | 46 | 11 | | 18 | 14 | 7 | 7 |
| **THF** | r | -0,057 | -0,217 | -0,162 | -0,173 | -0,148 | | 0,252 | 0,07 | -0,159 | 0,387 | | 0,299 | 0,22 | 0,079 |
| | p-value | 0,584 | 0,308 | 0,276 | 0,211 | 0,305 | | 0,017 | 0,529 | 0,468 | 0,125 | | 0,146 | 0,47 | 0,789 |
| | n | 96 | 25 | 47 | 54 | 53 | | 93 | 87 | 24 | 18 | | 26 | 14 | 14 |
| **5-methyl-THF** | r | 0,071 | 0,028 | 0,058 | 0,146 | 0,163 | 0,252 | | -0,029 | 0,047 | 0,463 | 0,299 | | 0,267 | 0,464 |
| | p-value | 0,434 | 0,88 | 0,643 | 0,25 | 0,168 | 0,017 | | 0,803 | 0,828 | 0,095 | 0,146 | | 0,609 | 0,294 |
| | n | 124 | 32 | 66 | 64 | 73 | 93 | | 79 | 24 | 14 | 26 | | 6 | 7 |
| **SAH** | r | 0,075 | -0,067 | 0,041 | -0,161 | -0,294 | 0,07 | -0,029 | | 0,045 | -0,687 | 0,22 | 0,267 | | 0,309 |
| | p-value | 0,508 | 0,798 | 0,806 | 0,276 | 0,047 | 0,529 | 0,803 | | 0,845 | 0,088 | 0,47 | 0,609 | | 0,329 |
| | n | 81 | 18 | 39 | 48 | 46 | 87 | 79 | | 21 | 7 | 14 | 6 | | 12 |
| **SAM** | r | 0,004 | | 0,004 | -0,628 | 0,81 | -0,159 | 0,047 | 0,045 | | -0,301 | 0,079 | 0,464 | 0,309 | |
| | p-value | 0,986 | | 0,99 | 0,029 | 0,003 | 0,468 | 0,828 | 0,845 | | 0,612 | 0,789 | 0,294 | 0,329 | |
| | n | 24 | | 11 | 12 | 11 | 24 | 24 | 21 | | 7 | 14 | 7 | 12 | |
